# Accurate transcription start sites enable mining for the cis-regulatory determinants of tissue specific gene expression

**DOI:** 10.1101/2020.09.01.278424

**Authors:** Mitra Ansariola, Valerie N. Fraser, Sergei A. Filichkin, Maria G. Ivanchenko, Zachary A. Bright, Russell A. Gould, Olivia R. Ozguc, Shawn T. O’Neil, Molly Megraw

## Abstract

Across tissues, gene expression is regulated by a combination of determinants, including the binding of transcription factors (TFs), along with other aspects of cellular state. Recent studies emphasize the importance of both genetic and epigenetic states – TF binding sites and binding site chromatin accessibility have emerged as potentially causal determinants of tissue specificity. To investigate the relative contributions of these determinants, we constructed three genome-scale datasets for both root and shoot tissues of the same *Arabidopsis thaliana* plants: TSS-seq data to identify Transcription Start Sites, OC-seq data to identify regions of Open Chromatin, and RNA-seq data to assess gene expression levels. For genes that are differentially expressed between root and shoot, we constructed a machine learning model predicting tissue of expression from chromatin accessibility and TF binding information upstream of TSS locations. The resulting model was highly accurate (over 90% auROC and auPRC), and our analysis of model contributions (feature weights) strongly suggests that patterns of TF binding sites within ∼500 nt TSS-proximal regions are predominant explainers of tissue of expression in most cases. Thus, in plants, cis-regulatory control of tissue-specific gene expression appears to be primarily determined by TSS-proximal sequences, and rarely by distal enhancer-like accessible chromatin regions. This study highlights the exciting future possibility of a native TF site-based design process for the tissue-specific targeting of plant gene promoters.

## Introduction

With the advent of genome-scale technologies, data has become increasingly available to address the intriguing question of when and where a gene will express in multi-cellular organisms. Since the entire genome of DNA sequence is identical in all of an organism’s cells, what information does RNA Polymerase II (pol-II) use to drive the transcription of many copies of a coding gene’s mRNA in one tissue or cell type, and very few copies in another? We know that “cause” is ultimately connected to a complex series of interrelated events that determine cellular state at the moments leading up to transcription initiation, including concentrations of various transcription factors (TFs) and nucleosomes, DNA methylation states, and histone modification states. Nonetheless, it is possible that DNA sequence alone contains all or most of the information necessary to determine the tissues in which the gene will strongly express.

Previous studies have largely focused on chromatin state and available TF binding sites as candidates for the primary determinants of tissue-specific gene expression, based on present mechanistic understanding of pol-II transcription initiation. The depth of understanding of pol-II promoter structure differs across multi-cellular eukaryotes (Smale and Kadonaga, 2003; Kadonaga, 2004; Thomas and Chiang, 2006; Sandelin et al., 2007; Juven-Gershon and Kadonaga, 2010; Kadonaga, 2012; Kumari and Ware, 2013) due to the timing of extensive genome-scale data availability across species. Foundational studies in Drosophila have strongly influenced concepts of ‘core’ promoter elements that reside at or immediately adjacent to the transcription start site (TSS) and regulate basal transcription, ‘proximal’ regions that extend beyond the core promoter but are also fundamentally important for transcription, and more distal ‘enhancer’ regions that are thought to regulate spatial and temporal control of transcription (Kadonaga, 2004; Ong and Corces, 2011; Spitz and Furlong, 2012; Kumari and Ware, 2013). While additional studies have broadened consideration of this paradigm over time particularly in vertebrates (Andersson, 2015; Feuerborn and Cook, 2015; Kim and Shiekhattar, 2015) and plants (Morton et al., 2014), still relatively little is known in many species about how ‘core’, ‘proximal’, and ‘enhancer’ regions are precisely defined genomically; the literature continues to focus on TF binding sites in more TSS-distal enhancer-like regions as candidate master-regulators of tissue specific gene expression (Ko et al., 2017).

The concept that ‘accessible chromatin regions’ or ‘chromatin footprints’ seem likely to pinpoint to specific regions of functionally bound TF sites, particularly in TSS-distal regions, has been presented since the advent of genome-scale open chromatin studies (Heintzman et al., 2007; Xi et al., 2007); this idea provides an attractive hypothesis that perhaps chromatin differences between tissues or cell types ‘modulate’ the patterns of TF binding sites that are available, thereby explaining gene expression differences between tissues. Many bioinformatic analyses support various forms of correlation between patterns of open chromatin and gene expression in a given tissue (Dong et al., 2012; Sheffield et al., 2013; Vera et al., 2014; Wilken et al., 2015; Rodgers-Melnick et al., 2016; Snyder et al., 2016). A recent study in plants (Ricci et al., 2019) specifically supports the idea that distal regions of open chromatin are statistically correlated with tissue-specific gene expression, and that some of these regions are likely to be enriched for relevant TF binding sites. Strikingly, however, there has been little evidence across the literature that distal accessible chromatin regions are primary drivers of tissue-specific gene expression in the case of most differentially expressed genes, or even that chromatin accessibility itself is largely determining which TFs are able to bind and functionally interact with pol-II. In fact, a recent study that includes mouse cells concludes in the title that “Accessibility of promoter DNA is not the primary determinant of chromatin-mediated gene regulation” (Chereji et al., 2019). In plants, studies comparing open chromatin landscapes across tissues and cell types (Zhang et al., 2012b; Zhang et al., 2012a; Pajoro et al., 2014; Sullivan et al., 2014; Maher et al., 2018; Lu et al., 2019) observe a surprising degree of qualitative similarity in general, including chromatin patterns surrounding differentially expressed genes, and it is not clear whether more refined, quantitative chromatin state differences may explain transcriptional program differences. The primary determinants of gene expression in a given cell type or tissue thus remain a provocative open question.

Machine learning models can potentially speak to this question by integrating genome-scale datasets of different types– including both DNA sequence and chromatin state information– in order to test whether enough information is present in these data types to predict tissue-of-expression related outcomes. Several studies have specifically contributed to this line of inquiry in the literature. An early study (Vandenbon and Nakai, 2010) used TF binding site information to test whether DNA sequences alone could predict the cell or tissue type in which a gene would be specifically expressed in 26 human and 34 mouse tissue and cell types. The study did this by training a series of models that would effectively predict whether a gene was more likely to tissue-specifically express in one tissue vs another in the same species, with the highest inter-tissue prediction success coming in at 73% auROC for human Kidney vs Fetal Liver (auROC is a performance measure of sensitivity and specificity, with a perfect model having auROC of 100% and a random classifier 50%). This study did use TSS information to define promoter regions, but high-precision genome scale TSS-sequencing data was not widely available at that time. A study from several years later incorporated genome-scale OC data (Natarajan et al., 2012) to examine 19 human cell lines, generating classifiers that predicted whether a gene would be strongly upregulated in different cell types. Median performance was reported for a variety of feature-generation techniques, the most successful technique achieved a median auROC of 73% by incorporating open chromatin information, with several of the top-performing models (out of 19 models) achieving an auROC of nearly 90%. Performance is not directly comparable with the (Vandenbon and Nakai, 2010) study, because the goal was not to predict the tissue of expression in a pair-wise setting but rather to distinguish genes that express very differently in a certain cell type than in other cell types. Features were interpreted for well-performing models, and several examples of tissue-specificity-associated TF binding locations within open chromatin were observed to be important to the models in these cases. It was clear in this study that use of chromatin information overall boosted performance, but it was very difficult to explain why just a few models performed very well while others did poorly. This study used the single annotated TSS location per gene, likely as genome-scale TSS-sequencing information was not available in all cell types of interest at that time.

Recently, the study (Agarwal and Shendure, 2020) trained a deep-learning model with the benefit of human ENCODE data that includes accurate transcription start site (TSS) locations in each cell type, as well as mRNA stability data. This study focused primarily on modeling transcript expression levels using TSS-sequencing data, but also trained a classifier to examine whether the cell type could be correctly predicted for cell-type-specifically expressing genes in human cell lines GM12878 and K562. The classifier achieved an auROC of 65% in predicting cell type from promoter sequence, but without use of TF binding site profiles of any kind. The promising model performance success of all of these outcomes supports the idea that both TF binding sites and chromatin accessibility carry considerable predictive power in classifying tissue of expression. Performance outcomes in the case of predicting tissue of expression in inter-tissue or inter-cell-line comparisons remain relatively low compared to the high 80%’s auROC that would be desirable for model interpretation. However, all of these past studies were necessarily limited to available data sets that either did not have precise TSS and chromatin information available in these same tissues, or—in cell line studies— cases where the material under examination did not come from the same individuals and did not come from normally functioning tissues. It was therefore not possible to inquire directly into the relative predictive success contributions of primary sequence information and chromatin state with the benefit of precise TSS locations in tissues or cells from the same healthy individuals.

This limitation may well have hindered predictive success in these past efforts, as precise TSS locations are necessary to correctly define promoter sequences relative to the pol-II binding site. TF binding sites are short (6-12nt) and sometimes degenerate sequences that appear throughout the genome by statistical chance. Therefore, if one does not know the actual location of each gene’s highly expressing TSS(s) in a tissue, and instead ‘guesstimates’ for each gene with a single annotated TSS (which is very likely to differ by at least 30-50 nt (nucleotides) from the actual TSS by as much as 500 nt in *Arabidopsis* (Morton et al., 2014)), then any ‘promoter sequence’ under consideration may be shifted many binding sites away from the actual TSS-proximal sequence. In this situation, not only is one unable to identify cases in which a different promoter sequence is being used to transcribe a gene in a different tissue, but one is completely unable to take advantage of accurate binding site patterns within each promoter—for example, on a very simple level one cannot even know whether a TATA site seen ∼25-35 nt upstream of a TSS is likely to be functional. Without the ability to approximate TSS location within a few nucleotides, it simply isn’t possible for a model to take advantage of precise patterns of relationships surrounding these binding sites over thousands of TSSs expressed on the genome in a given tissue sample. It will also be impossible to detect differences in these patterns for TSSs expressed in a different tissue sample, omitting an important source of information, as pol-II can utilize different transcription initiation locations for transcribing the same gene in different organs (Forrest et al., 2014; Mejía-Guerra et al., 2015). In essence, because binding sites are short and seen everywhere, important patterns in their relationships observed over thousands of TSSs (in either the same or different tissues) can simply be ‘washed away in the noise’ if each of those TSSs is randomly shifted away from its actual location by many binding sites in genomic distance.

In our study, we set out to construct a dataset that could help begin to quantitatively address the informative components in pol-II gene promoters that explain tissue of expression. The large and relatively complex yet well-studied genome of *Arabidopsis*, with many datasets from distinct tissues/organs during plant development, presented an ideal organism in which to undertake this task. We were able to construct a dataset from the same seedlings where each data component— while using relatively new technologies at the time for Transcription Start Site Sequencing (TSS-Seq) and Open Chromatin Sequencing (OC-Seq)—was able to be corroborated with other published datasets derived from similar material, indicating some stability in these data with regard to tissue/organ type, and allaying our concerns that our results might be particular to our sample material or particular to the technology/protocol that we used to produce the different dataset components. Our observations from machine learning model analysis of this dataset challenged our previous assumptions that distal chromatin-accessible TF site locations play a primary role in tissue of expression of most promoters, and suggest a possible paradigm shift in the way we generally assume plant promoters to operate. Specifically, our findings suggest that for the vast majority of differentially expressed genes in developing *Arabidopsis* organs, it is the pattern of cis-regulatory sites in the TSS-proximal DNA of these regions, regardless of chromatin state, that is most explanatory of the tissue of expression.

## Results

### TSS-Seq, RNA-Seq, and OC-Seq dataset in *Arabidopsis* roots and shoots captures chromatin state together with promoter utilization in different plant organs

Our study uses “root” and “shoot” tissues harvested from 7-day old *Arabidopsis thaliana* seedlings, dissected immediately below the hypocotyl (Figure 1). Each tissue batch was separated into three portions, to which we applied Transcription Start Site Sequencing (TSS-Seq), Open Chromatin Sequencing (OC-Seq), and RNA-Seq expression profiling protocols. We used the nanoCAGE-XL protocol (Cumbie et al., 2015b) for performing TSS-Seq, the DNase-I-SIM protocol (Filichkin and Megraw) for performing OC-Seq (Cumbie et al., 2015a), and applied a standard RNA-Seq protocol (see “Dataset Generation” section in Methods). The nanoCAGE-XL and DNase-I-SIM protocols were developed by the lab to work efficiently with relatively low-volume plant tissues such as *Arabidopsis* seedling roots and shoots; these were vetted in publication using datasets that were generated by applying other current protocols to comparable tissue samples sequenced on the Hi-Seq 2000, which sequenced to sufficient depth and coverage to support our study (see Supplementary Figure 1). We also noted that outcomes for both nanoCAGE-XL and DNase-I-SIM were remarkably consistent with other TSS-Seq datasets (Morton et al., 2014) and DNase-I-Seq datasets (Zhang et al., 2012a) generated in similarly prepared *Arabidopsis* seedling tissue samples.

**Figure 1.**
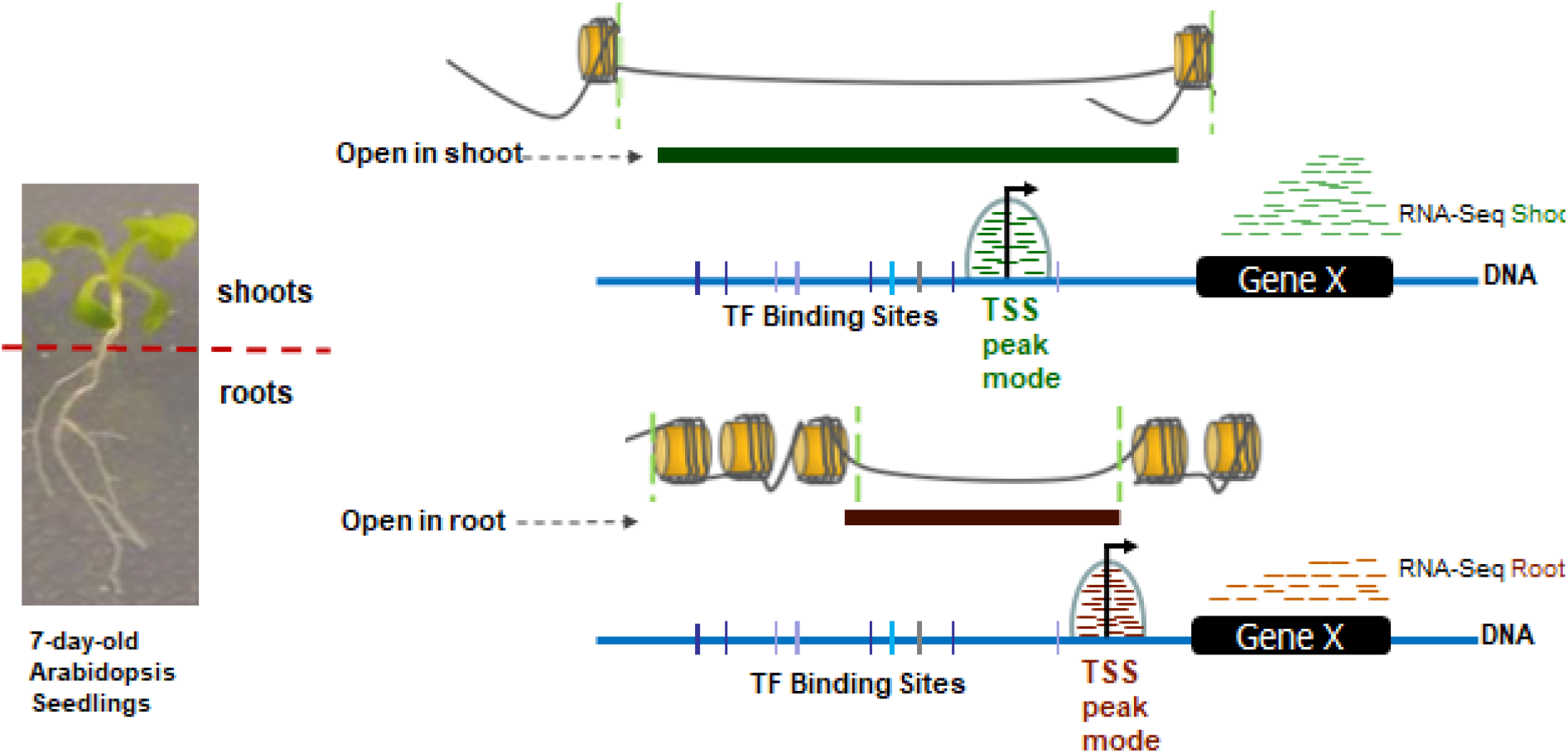
Datasets generated from 7-day-old wildtype *Arabidopsis thaliana* Columbia 0 roots and shoots. RNA-Seq reads align to annotated gene bodies to demonstrate gene expression, while OC-Seq reads highlight DNase-I Hypersensitive Sites. TSS-Seq reads align in peaks around the TSS; the mode of the peak is designated the location of the TSS.

Figure 1 illustrates the data collection goal of our study. As described in the Introduction, precise TSSs are critical to any predictive modeling effort that seeks to relate TF binding sites to gene expression outcomes, particularly those in different tissues, organs, or cell types. Clearly a reasonable estimate of highly accessible or “open” chromatin locations is also critical to our query, as it is plausible that TF binding sites within open chromatin regions are playing a large role in tissue-specific gene expression. Finally, we gathered RNA-Seq data for our samples as this form of expression profiling provides the most well-studied statistically robust estimate for gene expression levels, despite its implicit 3’ locational bias (Ross et al., 2013). We used these three data types to take an initial survey of apparent differences in gene expression program in our root and shoot samples (Supplementary Tables 1 – 4). Although we observed that TSS location and promoter accessibility are quite similar across tissues for many genes (Figure 2A), 2663 transcripts show strong expression in only one of the two tissue types (Figure 2C). Additionally, 525 differentially expressed transcripts are associated with a TSS peak in shoot only, while 707 transcripts have a TSS peak in root only. The 1431 differentially expressed transcripts with TSS peaks in both tissues results in 1632 peak pairs. Of these pairs, 222 (∼14%) have very different TSS mode locations (TSS-mode-distance > 100 bp), and 471 cases have mode locations that differ by 10 to 100 nt. Out of the 939 cases which have a very similar TSS mode location (TSS-mode-distance < 10), we looked for differences in patterns of open chromatin (% nucleotides disagreeing in chromatin state, Supplementary Figure 2). In the vast majority of cases, we were intrigued to see very little obvious difference in chromatin state—that is, we saw a large overlap in the percent of nucleotides agreeing in accessibility state. We observed that while major differences in TSS location or chromatin state might simply explain differential expression between the two tissues in perhaps 20% of the cases, in most cases the reason for differential expression could not be attributed to any obvious difference either in TF binding site usage or chromatin accessibility. We concluded that a quantitative modeling effort was necessary for further investigation.

**Figure 2.**
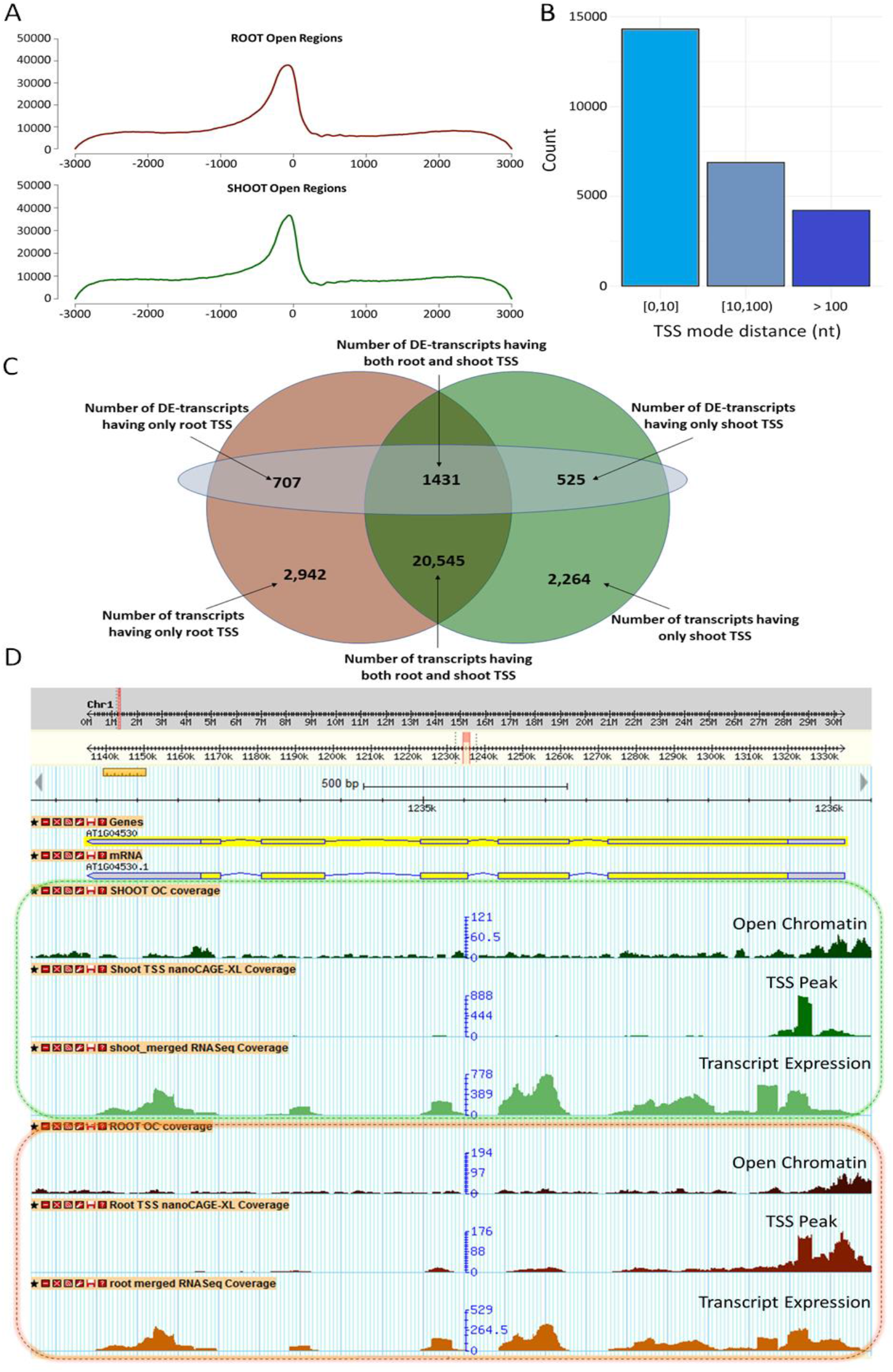
Summary of data outcomes. A) Charts comparing the general accessibility in roots (top) and shoots (bottom). B) C) Bar chart showing the difference in location between transcript-associated TSS modes in root and shoot. The majority of TSSs have similar locations in the two tissues, but ∼16% have very different locations (>100 nt). C) Numbers in the top row of the Venn diagram represent differentially expressed (DE) transcripts, separated by tissue in which the transcripts have associated TSS peaks. The numbers at the bottom represent all transcripts associated with TSS peaks, separated by tissue in which they have TSS peaks. Total number of DE transcripts = 2,663; total number of transcripts = 24,928. D) A sample gene displayed in GBrowse, showing mapped read accumulations for each data type.

### Highly expressed TSSs can be accurately modeled in each tissue type using only DNA sequence

Past machine learning studies have shown that strongly expressing TSS locations in a tissue sample can be precisely predicted from DNA sequence alone, using surrounding TF binding sites as ‘features’—that is, numerical descriptors of the genomic location that one is inquiring about with the question “Is this location a highly expressing TSS or not” (Megraw et al., 2009; Morton et al., 2014). We hypothesized that if one could accurately model highly expressing TSS locations in the root sample and in the shoot sample individually, using TF binding site (TFBS) information as features, one could then inquire into model differences in binding site patterns that are “important to root expression” vs those that are “important to shoot expression”. For this task, we selected the 3PEAT model (Morton et al., 2014), as it remains the only high-performance plant TSS peak finder to date with features that can explicitly be interpreted as representing TF:promoter binding site interactions. Additionally, the 3PEAT model had previously been applied to an *Arabidopsis* root sample grown under nearly identical conditions to those in our current study, but where the sample was generated using a different TSS-Seq protocol known as “Paired-End Analysis of Transcription start sites” or “PEAT” (Ni et al., 2010); this enabled us to understand whether our nanoCAGE-XL TSS-Seq datasets would support a similarly successful model to the PEAT root sample, which achieved an auROC in the high 90%’s. We applied the 3PEAT model to both root and shoot nanoCAGE-XL TSS-Seq samples from our study and found that we could predict strongly expressing root and shoot TSS locations in independent single-tissue-type models each with an auROC of 98%.

We then examined the TFs associated with the top-weighted features (TF binding locations most important to model success) of the trained root and shoot 3PEAT models (Supplementary Table 5, Supplemental Data Set 1), to determine whether there were any obvious root-specific or shoot-specific differences. We observed that the two models shared the majority of their top-50 most important TFs, though a few differences in the top-10 indicated the possibility of a more important role for root-development-related TFs in the root model and shoot-development-related TFs in the shoot model (Supplementary Figure 3). We then looked for quantitative evidence of root-specific vs shoot-specific TF binding site pattern usage in the two models by applying the root-trained model to the shoot model’s test set (i.e. locations that are either highly expressed TSSs or not highly expressed in shoot), and the shoot-trained model to the root model’s test set (i.e. locations that are either highly expressed TSSs or not highly expressed in root) (see Methods for details). Surprisingly, both models performed essentially identically on test sets of TSSs in the ‘other’ tissue as on test sets of TSSs from the tissue in which the model was trained (Supplementary Table 6) – with the same 98% auROC and only a negligible drop below 80% in auPRC (area under the Precision Recall Curve, a complementary performance measure). However, we found a tendency for the models to produce approximately 20% more false positives (classifying non-TSS sites as TSSs) when applied to the test set derived from the ‘other’ tissue compared to the model that was trained on that tissue. This was compensated by an 8% decrease in false negatives by the shoot-trained model on the root test set compared to the root-trained model and a 15% decrease in false negatives by the root-trained model on the shoot test set compared to the shoot-trained model (Supplementary Figure 4). Additionally, genes with TSSs that were misclassified by the shoot-trained model were statistically enriched for several root development GO-terms as compared to the full set of peaks that were tested (see Methods, Supplementary Table 7). Several additional GO-term enrichment and depletion observations, taken together with model feature-weight observations, strongly supported the idea that patterns of TF binding sites were likely to be well-explaining TSS expression in both tissues; yet the core of both models almost certainly described sequence information indicative of general transcription, as opposed to tissue-specific expression. We concluded that the 3PEAT TSS prediction concept provided an appropriate feature set that would potentially allow us to model the differences in tissue of expression based on TF binding site information, but would need to be incorporated into a model that focused on differentially expressing genes.

**Figure 3.**
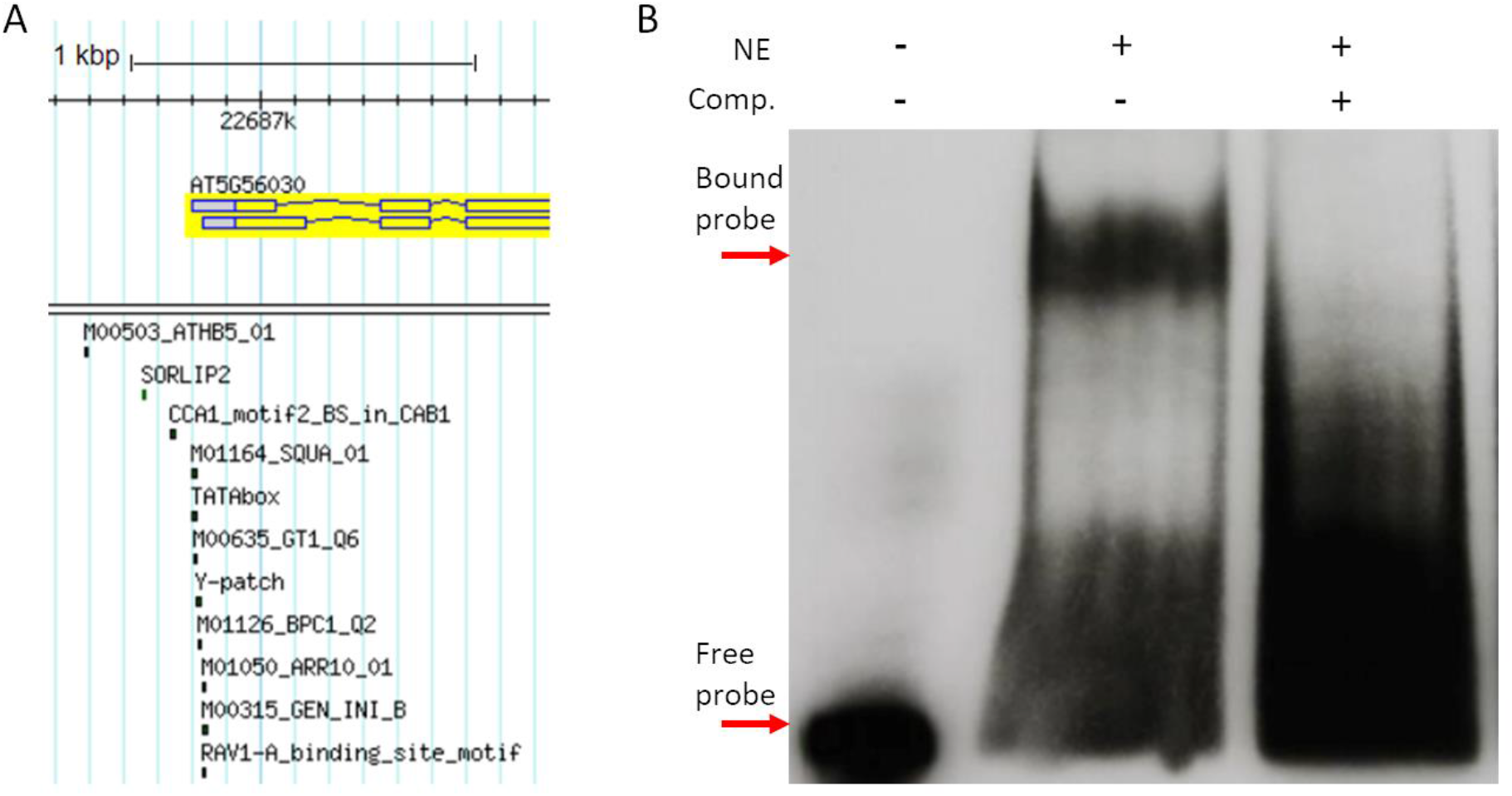
Selection and testing of putative functional binding sites. A) The HSP90.2 (AT5G56030) promoter region in GBrowse. Locations of the tested transcription factor binding sites in this promoter are displayed below the gene model. B) Y-patch electrophoretic mobility shift assay. The presence of a shifted band of probe indicates higher molecular weight than the free probe due to TF binding. The right-most lane contains >200X cold competitor to show that the shift is not an artifact.

**Figure 4.**
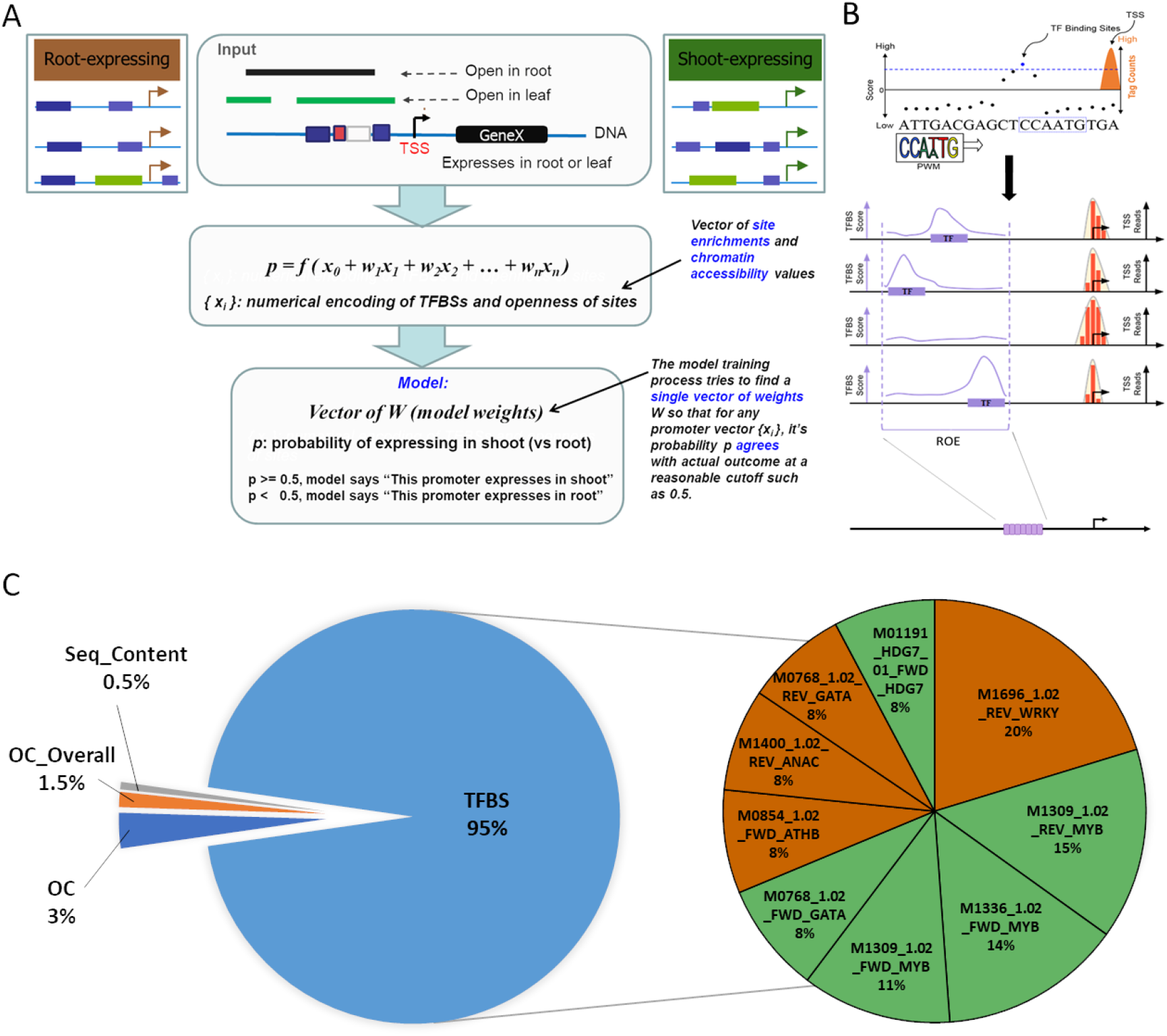
A) TEP model concept: Three data types (TSS-Seq, OC-Seq, RNA-Seq) are generated from roots and shoots; this allows us to numerically encode TFBS presence and chromatin accessibility. The numerical encoding is then used to train and test a machine learning model that outputs the probability of a transcript’s expression in root or in shoot. B) TEP-ROE feature generation: Regions of enrichment (ROEs) are detected by scanning a PWM (TF binding profile) over each promoter region and calculating where the TFBS loglikelihood scores are significantly higher than background levels. These regions are then further divided into windows for feature scoring. C) In the chart to the left, TFBS features make up 95% of the total generated. The chart on the right highlights the features that the TEP-ROE model weighted most heavily. The names of these features contain quite a bit of relevant information (e.g. M1969_1.02_REV_WRKY: M1969_1.02 is the PWM designation from the database; REV indicates that the feature is located on the opposite strand from the gene; and WRKY is the associated TF family). Green pie wedges indicate that the model deemed this feature important for expression in shoots, while orange wedges indicate importance for expression in roots.

### TSSs enable meaningful TF binding-site-based feature set construction

The original 3PEAT model investigation (Morton et al., 2014) demonstrated that precise TSS locations were key to training a highly accurate TSS prediction model, with a substantial ∼10% auROC performance drop if only annotated start sites were used. We wanted to investigate whether the 3PEAT model’s TF binding site-based feature set construction was not only the key to predictive success in explaining strong TSS expression, but also carried plausible support for explaining pol-II transcription in reality. Specifically, we wanted to test whether the putative binding sites modeled as important to a gene’s correct TSS location prediction were also likely to be TF-bound in the sample, therefore potentially functional. We also wanted to gain a basic indication of whether ‘predictive sites’ extracted from model features—that is, important TF binding site-enriched regions known in the PEAT model as “Regions of Enrichment” or ROEs (Figure 4B)—were likely to be highly sensitive to the specific dataset in terms of sample collection, TSS-Seq protocol, or informatic processing details such as selection of peak caller parameters. We selected the root sample for testing, as it was then possible to compare 3PEAT TSS models built from two different datasets using very similar tissue samples, (i) the PEAT dataset (Morton et al., 2014) and (ii) the nanoCAGE-XL dataset generated for the present study.

We began by selecting putative cis-regulatory sites from the original 3PEAT model application to the PEAT dataset for *in vitro* TF protein:DNA binding interaction testing using the following procedure (see “Functional Binding Site Selection” in Methods for details). Cis-regulatory elements considered by the model included TF binding sites as well as core promoter elements such as TATA-box which facilitate direct interactions with the pol-II complex. First, likelihood scores for individual putative binding sites that contributed to each TSS prediction (i.e. each transcript detected in the *Arabidopsis* root) were calculated using their corresponding Positional Weight Matrix (PWM) binding domain representation and their position relative to the ROE for each element. The output of this pipeline was a genome-wide “master-list” of potentially functional cis-regulatory sites. These candidates were filtered by considering only strongly expressing “narrow-peak” TSSs, which have an enriched association with developmental genes responsible for tissue-specific patterning (Morton et al., 2014), and other restrictions to generate a stringent short-list of 500 sites (i.e. sites associated with the top 20% by importance-rank according to their 3PEAT model weight, then sites with highest likelihood scores located near the center of their ROE and within 120bp of their corresponding TSS). Finally, we selected five “high-scoring” candidate sites (INI-B, TATA box, Y-Patch, PIF3-binding element, and SQUAMOSA Promoter Binding element SQUA1) for evaluation using the Electrophoretic Mobility Shift Assay (EMSA or “gel shift” assay) and nuclear extracts prepared from *Arabidopsis* roots. We included several sites from the HSP90.2 promoter, one of the few genes with top-ranked sites in our list that had a known function, as well as one site each from the promoters of ornithine carbamoyltransferase (OTC) and diacylglycerol kinase 2 (DGK2); in this first selection, we focused on sites that were located in regions of open chromatin in root, but did not account for expression level of the TF(s) corresponding to the PWM binding domain profile predicted to target the candidate sites. We observed gel-shifts for four out of the five candidate sites.

In troubleshooting the case that did not shift, we observed that PIF3, a circadian-controlled TF, had a very low level of expression as measured by the RNA-Seq outcome in the root sample of our current study. We also observed a general qualitative correlation between the intensity of the shifted band and RNA-Seq expression level with the other candidates, suggesting that TF binding would be undetectable below a certain level in nuclear extracts. We then selected 6 additional top-scoring sites in the HSP90.2 promoter (Figure 3) but filtered out any sites associated with very lowly expressed TFs as measured by our RNA-Seq sample. Of these sites, all 6 resulted in a shifted band, indicating binding. We then considered whether applying the same process to the 3PEAT model, when trained using more recent peak-calling on the nanoCAGE-XL root sample, would include these same sites as relatively important to the expression of the target gene. We repeated the process (“3PEAT-style Model Construction” in Methods) using the 3PEAT model re-trained on the nanoCAGE-XL root sample that achieved an auROC of 98% described in the section above. We observed that all of our selected sites in the HSP90.2 promoter were included in the new top sites list, although some had a lower score; the tested site in the DGK2 promoter does not appear on our new list, though this gene had a very weak TSS peak in the sample, consistent with involvement in circadian function including lack of upregulation by PIF3. OTC had a moderate TSS peak in the sample, and its site was included (Supplementary Figure 5). In considering whether additional information could be obtained by performing traditional gel-shifts for these sites using purified TF protein, we concluded that if successful this would only demonstrate the ability to bind an oligo at unrealistically high concentrations of each TF, essentially confirming the TF’s PWM binding domain description as provided by a database. In total, the gel-shift of 10/11 predicted binding sites using nuclear extracts, taken together with qualitative correspondence between RNA-Seq level of the candidate TF and darkness of the shifted band, as well as the relatively stable predictive importance of these sites across TSS-Seq datasets, provided plausible support for binding of these sites at *in vivo* TF concentrations. We concluded that 3PEAT model’s ROE-based TF binding site features represent sites that are at least potentially functionally bound in a way that promotes their target gene’s transcription by pol-II.

**Figure 5.**
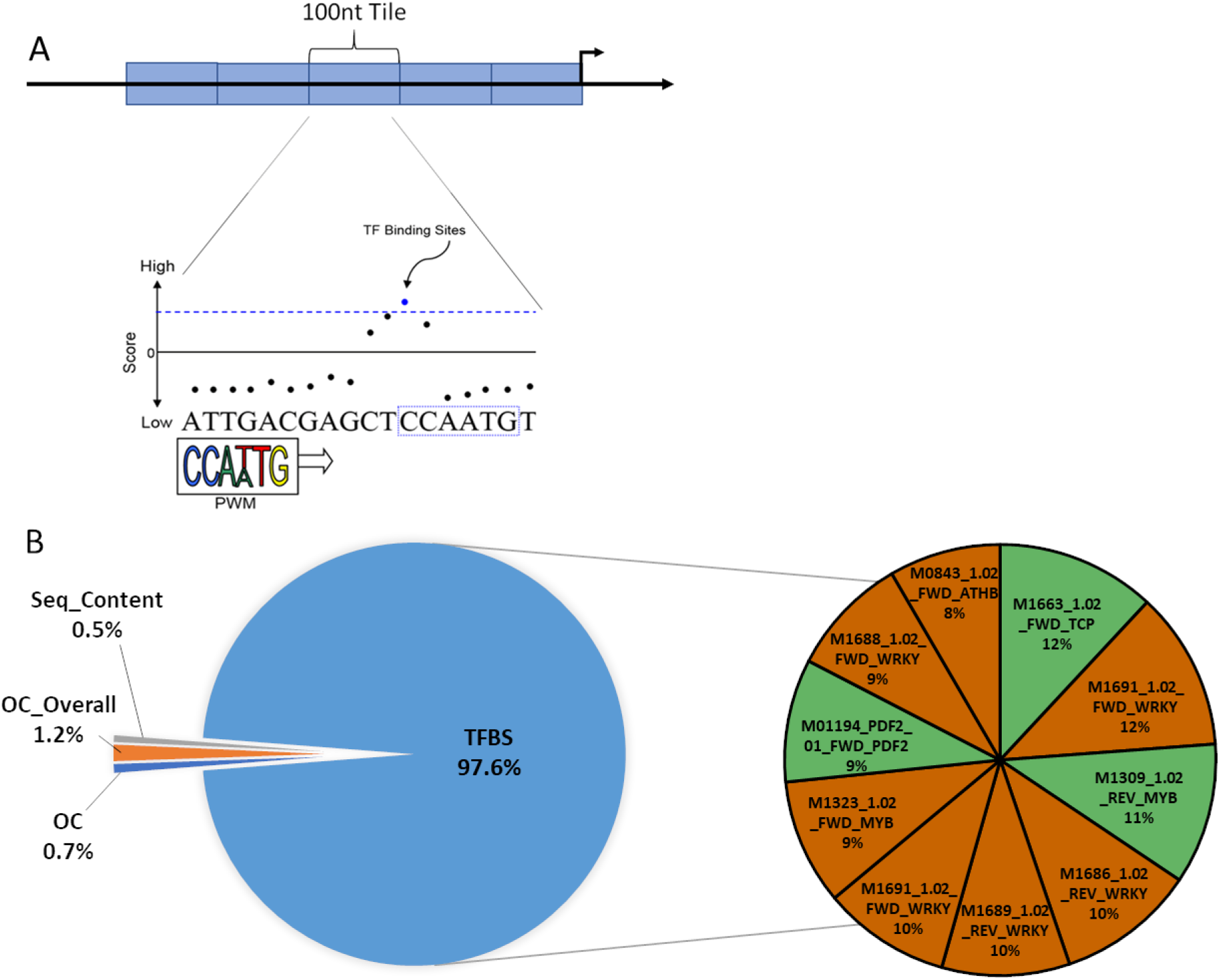
TEP-Tiled model. A) Cartoon schematic of TEP-Tiled model’s feature generation. Instead of identifying ROEs and creating features within these smaller regions, this model generates 100 nt wide tiles over the entire promoter. B) As with the TEP-ROE model, TFBS features comprise the majority of the features generated by the TEP-Tiled model (97.5%). The pie chart to the right contains the top 10 most heavily weighted features. As with the TEP-ROE model, feature names have three parts (e.g. M1691_1.02_FWD_WRKY: M1961_1.02 is the database’s identifier for the PWM; FWD means the feature is on the same strand as the gene; and WRKY is the associated TF family). Green pie wedges are features that the model deemed important for expression in shoots and the orange pie wedges are features the model deemed important for expression in roots.

### TFBS locations and their chromatin state accurately predict tissue of expression for differentially expressed genes

Building on the successful TF Regions of Enrichment feature concept of the 3PEAT model for predicting TSS location, we constructed an analogous model that we called the Tissue of Expression Prediction ROE model or TEP-ROE model. We reasoned that if patterns of TF binding site enrichments can predict the locations of strongly expressing TSSs on the genome, and high-affinity binding sites within important enrichment regions are plausibly functionally contributing to pol-II’s frequent transcription initiation at these locations, then perhaps it is patterns of TF binding sites within these regions that can help well-distinguish a tissue in which a gene will express strongly from a tissue in which it will express to a much lesser extent. But it also seemed that the general accessibility of sites in these regions could prove important, as could general sequence enrichments such as AT-content and overall degree of openness in the vicinity of the TSS. Figure 4A shows the concept of the TEP-ROE model, with details provided in Methods. Like 3PEAT, TEP-ROE is an L1-regularized logistic regression classifier that takes as input (i) the DNA sequence surrounding a TSS (TSS - 1 kb, TSS + 500 nt) and (ii) chromatin accessibility state for both tissues in this region around the TSS, and returns the predicted tissue (root or shoot) in which that TSS will express most strongly. The most important concepts for understanding and interpreting the model (Figure 4A) are that (1) each TFBS feature represents a specific genomic region in relationship to a TSS where a particular TF binding domain has a high density of high affinity binding sites, (2) each OC feature represents the “openness” (degree of accessibility) of a corresponding TFBS feature region, as a percent of accessible nucleotides in this region, (3) the two OC_overall features OC_overall_root and OC_overall_shoot represent the percent openness of the ‘proximal’ region [TSS - 500 nt, TSS + 100 nt] around a TSS in root and in shoot, (4) sequence enrichment features (e.g. GC Content) represent the percent of certain nucleotides (e.g. G and C) in a 100 nt window around the TSS, and (5) the weight that a successfully trained model gives to each of these features represents an ‘importance value’—a large weight magnitude or “top-ranked feature” indicates a feature whose value contributes heavily toward the decision about whether a TSS is predicted to express strongly in root or strongly in shoot.

We trained the TEP-ROE model on TSS locations associated with differentially expressed genes, using cross-validation for parameter selection and an independent held-out test set for reporting test performance (see “Model Training and Testing on nanoCAGE-XL TSSs” section in Methods). The model achieved an auROC of 92% and an auPRC (area under the Precision Recall Curve, an important co-indicator of performance) of 94% (Supplementary Figure 6). In order to evaluate performance stability over a wide variety of dataset divisions (training vs testing) and algorithm seedings (different initial value settings of the optimization algorithm), we re-trained the model 30 times with different ‘seeds’ (Supplementary Figure 9). We observed that auROC and auPRC model performance outcomes were tightly distributed around means which were close to our TEP-ROE model’s performance values and concluded that our TEP-ROE model’s strong performance was representative. We then examined feature stability by looking at the feature weight ranking distribution for each of the 50 top-ranked features in the model, over the 30 models used in performance stability testing (Supplementary Figure 10); we found feature rankings to be acceptably stable in the sense that each of the 50 most important features stayed within the top-ranked 50 for all other test models, and most features’ rank remained within 5-10 ranking slots of the mean in the vast majority of the test models.

**Figure 6.**
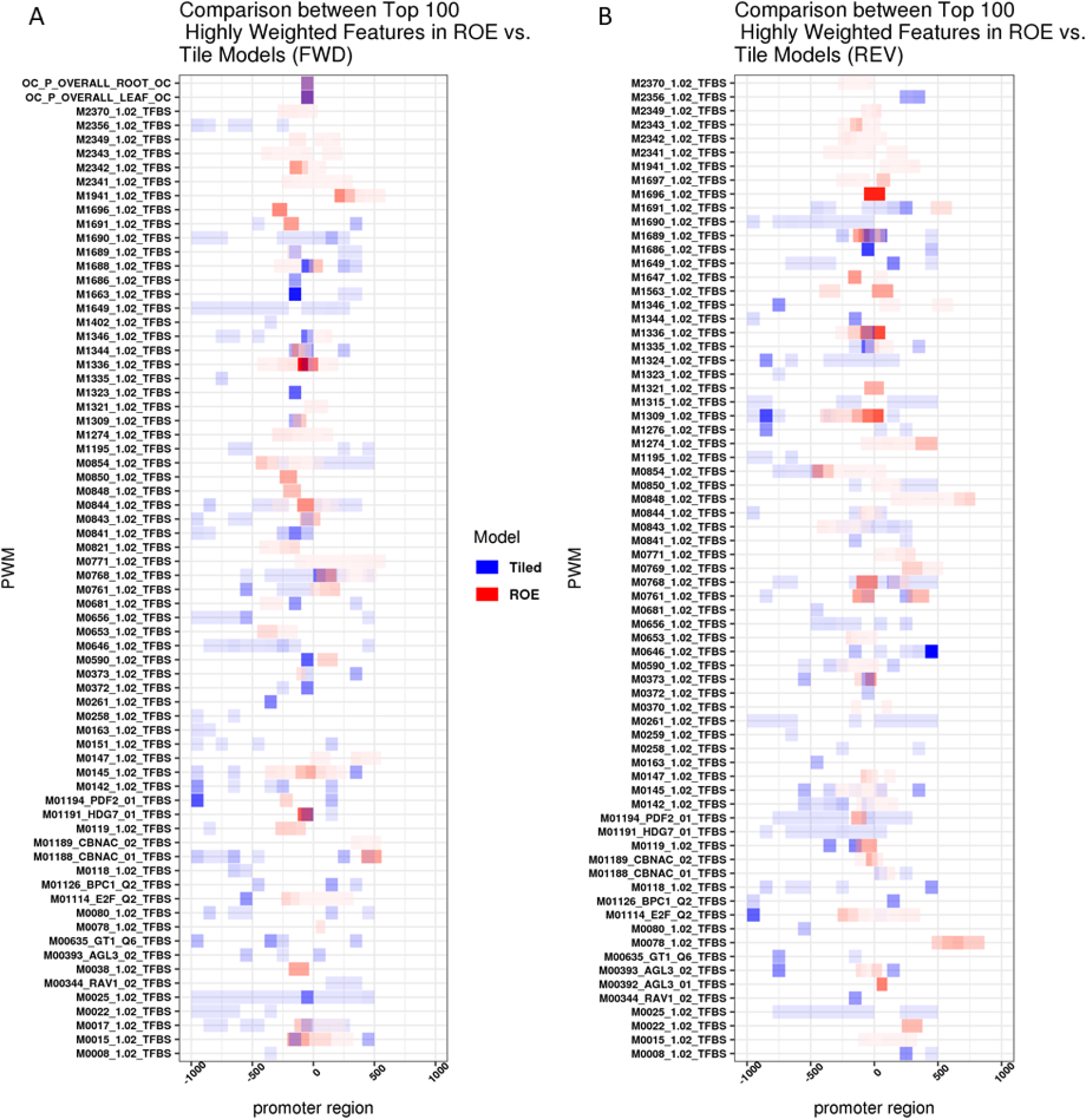
Comparisons of top PWMs between TEP models. A) Heatmap showing the differences between and shared PWMs on the same-as-gene (FWD) strand that are weighted highly by TEP-ROE (red) and TEP-Tiled (blue). B) Heatmap showing the differences between and shared PWMs on the opposite-from-gene (REV) strand that are weighted highly by TEP-ROE (red) and TEP-Tiled (blue).abase’s identifier for the PWM; FWD means the feature is on the same strand as the gene; and WRKY is the associated TF family). Green pie wedges are features that the model deemed important for expression in shoots and the orange pie wedges are features the model deemed important for expression in roots.

Performance and stability indicated that it was meaningful to interpret the TEP-ROE model, as top-weighted features were likely to be important contributors to successful tissue prediction. The two OC_overall features were high-ranked contributors (Figure 4C, Supplemental Data Set 2), with a few general sequence content features falling into the top 300. The most striking aspect of the model outcome is that aside from the OC_overall features for root and shoot, and a small number of sequence content features, TFBS features comprised all of the top ∼350 features, with the first OC feature appearing at rank 352. We performed a literature search on the top 100 TFBS feature binding domains, and found that of the 20 which had functional annotation, four had literature support for activity in the same tissue whose weight sign (positive or negative) indicated that presence of this TFBS site density made expression in this tissue more likely. The locations of these important regions fell within 500 nt of the TSS, indicating the strong predictive role of TF site densities in this proximal region. Finally, when we re-trained a version of the TEP-ROE model using only the TAIR10 annotated start site for each differentially expressing gene rather than TSS-Seq peak locations, auROC dropped to 76%. This substantial ∼15% auROC performance drop supports an important role for precise TSS locations in successful tissue of expression prediction model training.

### Promoter ‘tiling’ model offers complementary view of important feature locations

The TEP-ROE model was constructed around the concept of “Regions of Enrichment”, which are special regions that one can think of as containing “high TF binding site densities” for a particular TF with respect to all TSSs in a sample type. Since this model style focuses only on a single Region of Enrichment for each TF, and not all TFs have these high binding site densities in our root and shoot samples, some TFs and their binding sites are omitted from consideration in the TEP-ROE model. We also wondered if the TEP-ROE concept was unnecessarily “confining” important TF binding patterns that are considered by this model to locations very near to the TSS, just because this is where the highest binding site densities occur for most TFs. This lead us to ask whether a model that simply “tiled” the same region surrounding the TSS with ‘tile regions’ (Figure 5A) would achieve similar or even greater performance—and if it did, would such a model select similar TF binding site density regions as important features. We constructed the TEP-Tiled model by following an identical procedure to the TEP-ROE model, except that ROEs were replaced by a series of non-overlapping 100 nt windows tiling the entire [TSS - 1 kb, TSS + 500 nt] region under consideration (see “Promoter Tiling” section in Methods).

The TEP-Tiled model achieved nearly identical performance results (Supplementary Figure 7) to the TEP-ROE model, and its performance over a large number of seeded trials was similarly stable (Supplementary Figure 9). Building the model using only annotated TSSs from TAIR10 caused a similar ∼15% auROC performance drop to that of the ROE model. The stability of top-weighted features decreased as compared to the TEP-ROE model (Supplementary Figure 11), but this is largely to be expected because the TEP-Tiled model has many thousands of additional features (many more tiles than ROE regions) and is therefore a very highly under-constrained model; that is, there are so many more features whose importance the model must consider than there are TSSs in the root and shoot classes that (1) there are many feature combinations that can potentially help the model to perform well, and (2) the model operates at the limit of the regularization process’ ability to identify meaningful feature combinations. It is this second issue that lead to declining performance when we examined models using tiles smaller than 100 nt wide. Nonetheless, the TEP-model’s strong and stable performance provided the ability to meaningfully examine the type and location of its most important features (Figure 5B). The OC_overall features play a similarly important role, and sequence content features have similar rankings in general. The first TFBS-associated OC region appears at a lower importance rank (∼740) than in the TEP-ROE model (∼350), as TFBS features even more heavily dominated the top importance weight rankings.

**Figure 7.**
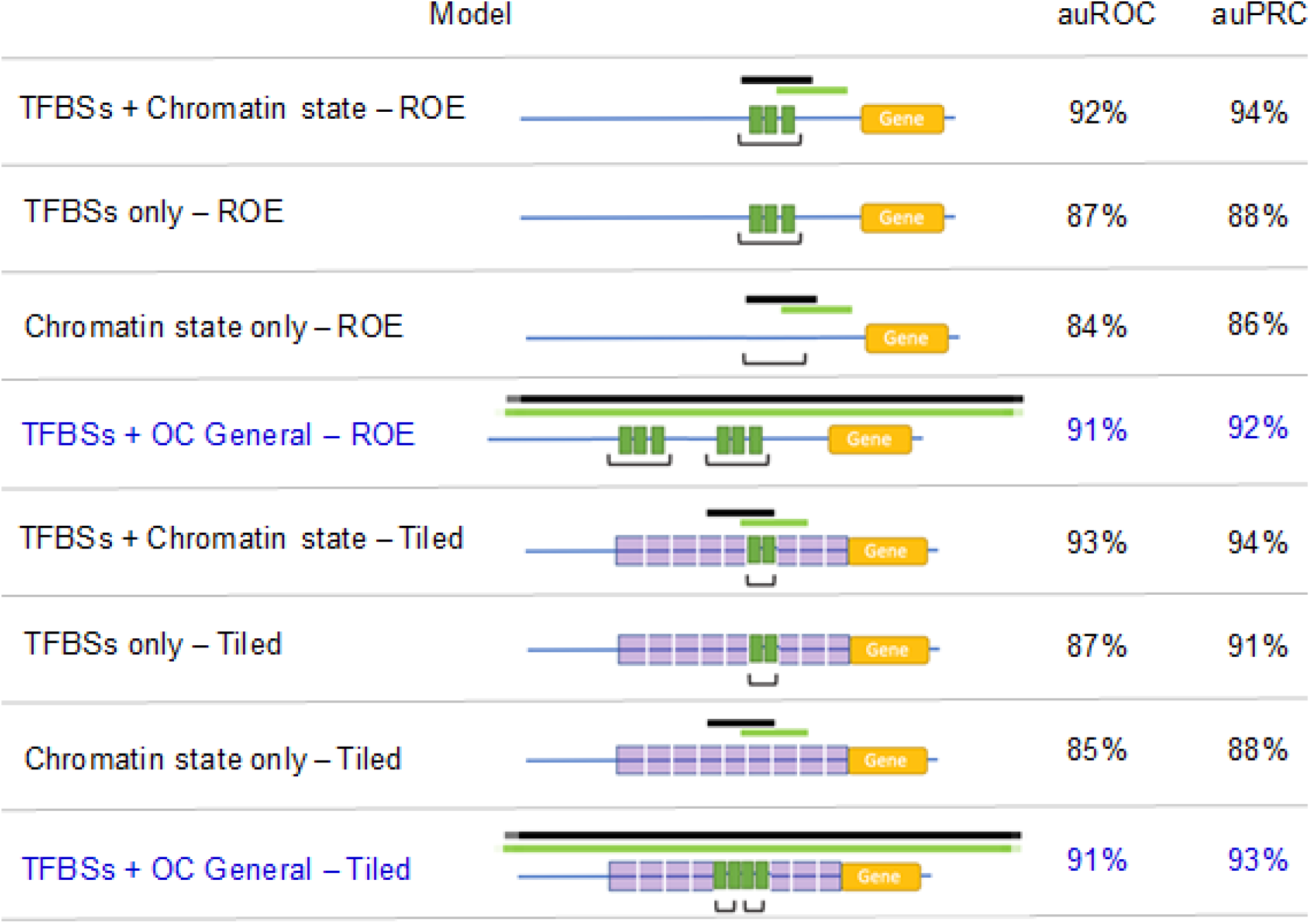
Performance summary. The removal of chromatin information from our feature set results in a performance decrease of 5-6% in both of our TEP model types. Removal of sequence-based information decreases performance by 8-9% in the TEP models. Addition of a general openness feature for the promoter region to the TFBS only TEP models (blue-highlighted lines) restores performance essentially to the same level as the original models containing all chromatin features.

To visually examine the relationships between top-ranked TFBS feature locations in the TEP-Tiled and TEP-ROE models, Figure 6 shows a heatmap overlay of the 100 top-weighted features in each model, displayed according to location with respect to the TSS. In general, for most of the TFBS features that the two models agree are in the top 100 (those rows that contain both red and blue hues), there is some form of ‘telescoping effect’ or overlap in the regions that both models consider highly important. In these cases, typically the Tiled model agrees with at least one of the locations that the ROE model considers most important for a TF binding domain type, but also gives some lesser weight to at least one additional location. This seems to suggest that much of the time, when the models agree on an important TF binding domain, there is a tendency to agree on its most important location. But, clearly, there are many ‘only red’ or ‘only blue’ rows indicating that there is agreement on inclusion of a TF binding domain feature only about one third of the time. Supplementary Table 8 provides a quantitative look at whether the two models agree on what the most important TFBS or OC features are, location aside. In considering the top 10-weighted features in each model, about 30% are shared. However, all disagreements in this case appear to result from selection of different members of the same TF family by each model (among TF-associated features), as M1691_1.02_TFBS, M1686_1.02_TFBS, M1696_1.02_TFBS are all WRKY family transcription factor binding domains. In general, 20-30% features are identical between the two models in considering up to 200 top-weighted features, and dissimilarities appear to be due at least in part to the different models’ inclusion of non-identical but relatively numerically similar binding domain profiles among TF families.

Finally, while Figure 6 shows that the most important locations in both models tend to fall within 500 nt of the TSS, the TEP-Tiled model indicates that occasionally an important TF binding domain location could be located nearly 1 kb upstream of the TSS. The most important ROEs from the TEP-ROE model all lie within 500 nt of the TSS, but we wondered if the TEP-Tiled model would select important tiles more than 1 kb upstream if given the opportunity. We retrained the TEP-tiled model using [TSS - 2 kb, TSS + 500 nt] and observed a slight performance drop, with no top-10-ranked tiles in importance falling upstream of 1 kb (Supplementary Figure 14, Supplementary Table 9). Overall, comparisons between models support the idea that the TF binding locations which contribute most to model performance—that is, to the model’s ability to correctly predict the tissue of expression—lie within about 500 nt of the TSS.

### TEP models suggest some promoters may express almost solely based on patterns of functionally bound sites

In examining the target genes of TFs that were identified by a model as very important to differential expression, we observed that the promoters of some TSSs seemed to contain high-affinity binding site densities in important regions for several different TFs. We were interested to investigate whether some promoters seemed to be “hard-coded”, in the sense that the promoter’s associated tissue of expression appeared to be entirely dictated by TF binding site patterns, with no influence from chromatin state according to a successfully trained Tissue of Expression Prediction (TEP) model. We ran an analysis to identify promoters where tissue of expression prediction was successful and nearly all of the promoter’s ‘important feature products’—meaning high-affinity TF binding site densities (TFBS features) or large chromatin accessibility values (OC features) that were associated with high model weights (feature importance values) — derived almost entirely from the presence of high-affinity TF binding site densities. We identified promoters for 18 genes that fell above the 95^th^ percentile for ‘hard-codedness’ (see Methods) using the TEP-ROE model, and 43 genes using the TEP-Tiled model (Supplementary Tables 10 and 11). Both gene sets were enriched for GO terms associated with metabolite biosynthesis and transport (Supplementary Table 12), while the TEP-Tiled model ‘hard-coded’ genes were additionally associated with development.

The models suggest that genes whose promoters are ‘hard-coded’ by TF binding site content to express differentially in roots vs shoots (or vice versa) could be preferentially involved in very basic processes that need to be performed differently in one tissue vs another during development, based on TF presence alone. This implies that chromatin state in these cases is perhaps directly modulated by one or more of the TFs involved in the important binding site patterns; we did observe that in the case of each model, at least one ‘top TF binding domain’ associated with the most important feature products was known to be involved in chromatin remodeling (Supplementary Table 12).

As a result of these inquiries, we wondered whether there existed cases of promoters such that ‘zeroing out’ a single TF binding site density would be predicted to ‘flip’ the tissue in which a gene was most highly expressed. The physical analog of this experiment would be a form of ‘insilico knockout’, where high-affinity binding sites within an important region for a TF’s influence on tissue of expression are removed or occluded, so that this TF can no longer bind in this region with respect to the current TSS. We observed that the TEP-ROE model points to 8 genes whose promoters contain ‘knockout regions’ that would cause a 50% probability shift ‘across the decision boundary’ to change the predicted tissue of strongest expression (Supplementary Table 13). The TEP-Tiled model, which has many more regions than the TEP-ROE model, points to 28 such TF-tile ‘knockout’ locations that cause at least 50% probability shifts or greater to ‘flip’ predicted tissue of strongest expression (Supplementary Table 14). For the TEP models, several of these tissue-flip-causing TF binding site density ‘knockout’ regions were also among the top TF binding domains important to ‘hard-coded’ promoters. This outcome appears to corroborate the presence of highly influential binding region locations for specific TFs that may be serving in an important ‘master regulatory’ role for the tissue of expression of some promoters. Collectively, modeling experiments suggest that it is typically pairs or larger groups of these important TF binding density regions located in spatial patterns that most heavily influence tissue of expression within the ∼500 nt upstream proximal promoter region of a transcript.

### TF site presence and location are predominant explainers of tissue of expression

Although TFBS features were highly dominant in both the TEP-ROE and TEP-Tiled models, we unexpectedly observed that the chromatin regions surrounding the important TFBS feature binding site density locations were not considered important at all by the model. In fact, TFBS feature weight values assigned by each TEP model had virtually no correlation with model weight (importance) of the corresponding region of chromatin (Supplementary Figures 15 and 16). Additionally, we observed no correlation between the importance of regions containing large TFBS densities within individual TSS promoters and chromatin openness in these regions (Supplementary Figures 17 and 18). This was perplexing; because model structure means that important TFBS features represent locations where large TFBS densities contribute strongly to expression in one tissue vs another, we had anticipated that the state of chromatin accessibility of these regions at least in some cases would be correspondingly important. We noted that the OC_overall features, representing openness of the general proximal [TSS – 500 nt, TSS + 100 nt] region, received high weight in both tissues for both models. Both models also agreed on TFBS features as overwhelmingly more important than OC features as a collection. It seemed possible then that a higher degree of general openness of this TSS-proximal region was contributing the vast majority of chromatin state information in the models.

We decided to examine whether removal of OC features entirely would seriously hurt model performance. We therefore re-trained both the TEP-ROE and TEP-Tiled models on feature sets that were identical to the original models with the exclusion of any OC feature. We observed a ∼5% drop in auROC in both cases (Figure 7), with the resulting “TFBS-Only” models performing surprisingly strongly at 87% auROC, with 88% and 91% auPRC respectively. We then wondered if a TFBS-Only model with root and shoot OC_overall features included, but no other OC features, would perform as well as the original model. We tested this idea, and both model types achieved essentially the same performance as the original TEP-ROE and TEP-Tiled models achieved with all feature types present (Figure 7). Thus, by returning only a single general measure of openness of the [TSS - 500 nt, TSS + 100 nt] proximal promoter region in each tissue to a DNA-sequence-based-features-only version of each model, performance was restored to essentially the same level as when all OC features were included.

Finally, we expected that the ‘inverse’ experiment-- removal of all TFBS features-- would not seriously hurt model performance, because there is a strong tendency of expressing promoters to be more open (Figure 2A) within ∼1 kb upstream or so of the TSS; thus, open chromatin in general should be a strong predictor of differential gene expression. We went forward with this experiment and observed that both models performed with an auROC of ∼85% (Figure 7), a ∼7% drop in auROC from the original TEP-ROE and TEP-Tiled models. This result was surprising only in that it suggests that chromatin-state patterns, at least on a regional scale, can predict tissue of expression in strongly differentially expressing cases only about as well as TFBS and other DNA sequence content information alone. By far, the highest weighted coefficients in this model were the OC_overall root and shoot coefficients, corroborating the predictive power of general chromatin openness within the [TSS – 500 nt, TSS + 100 nt] region.

While it is possible that examination of OC patterns on a binding-site-scale might contribute some additional information if it were to become technically feasible to train such a model, all outcomes in our set of experiments clearly suggest that TF binding site information within promoter DNA is the predominant explainer of the tissue of expression. In particular, outcomes support the concept that patterns of TFBS densities within the ‘more-open’ ∼500 nt proximal promoter region provide the largest influence on the tissue(s) in which a gene will preferentially express. Outcomes support a secondarily influential role for the degree to which a promoter is generally open in this region within a tissue, although of course the modeling experiments cannot inform whether this higher degree of proximal-promoter openness results from a chromatin remodeling process that is TF-dependent. Surprisingly, modeling outcomes did not support any direct association between important TF site density locations and the importance of chromatin state in these locations.

## Discussion

### Model success in predicting tissue of expression fundamentally derives from precise TSS information

The Tissue-of-Expression-Prediction or “TEP” models constructed in our study are conceptually straightforward classical machine learning models— they use L1-regularized logistic regression to find specific high-affinity TF binding site regions positioned in relationship to a TSS, along with chromatin accessibility values in these regions, that collectively identify the correct tissue of greater expression for the vast majority of differentially expressed genes in our sample set (auROC ∼90%). Previous attempts at this specific task in similarly genomically complex organisms have made good progress but achieved middling results at best (auROC ∼75%). Why then did the TEP models perform so well as compared to past models, including a recent deep learning model in cell lines (Agarwal and Shendure, 2020) that did in fact have precise genome-wide TSS-Seq information available? The construction of a first-of-its-kind dataset with generation of both Transcription Start Site sequencing data and Open Chromatin sequencing data in two different tissues of the same healthy individuals likely contributed to success. However, all of our experiments clearly indicate that the most important contributor to predictive success was modeling with accurate TSSs in each plant organ. Our outcome is consistent with similar classical machine learning studies including (Vandenbon and Nakai, 2010) in demonstrating that TF binding site information alone is capable of achieving relatively strong predictive performance, and (Natarajan et al., 2012) in confirming that chromatin accessibility information boosts inference of genes that are expressing differently in different tissues/cell-types. In relationship to the (Vandenbon and Nakai, 2010) study, which performed an identical task with auROC of 75%, the TEP models’ ∼15% auROC performance drop when only annotated start sites were used is consistent with the idea that accurate TSS information within each tissue is largely responsible for dramatic performance boost.

However, precise TSS information alone is unlikely to be solely responsible for high sensitivity and specificity given that a sophisticated DNA-sequenced-based deep learning model had the benefit of TSS-seq data but achieved only ∼65% auROC on this task; (Agarwal and Shendure, 2020) concludes in fact that the model’s performance is not boosted by the use of TSS-seq data instead of annotated start sites. Given the observations in our study, it is very likely that explicit use of important biological information such as TF binding profiles in feature set construction confers a large benefit in predicting tissue of expression. TEP model feature sets are carefully constructed to use high-affinity TF binding site densities as opposed to thresholding, and to encompass positional relationships of these densities to the TSS. Additionally, despite its relative simplicity as a classical machine learning model, regularized logistic regression is a time-tested method that still routinely outperforms deep-learning approaches in genomic classification of phenotype from transcriptomics data (Smith et al., 2020). While deep learning models hold exciting promise, it seems likely that current architectures are as-yet unable to learn TF binding site features with enough precision to take advantage of patterns in their positional relationships to each other and to the TSS.

We would hypothesize in this context that much of the “missing mass” in performance to bring auROC up near 100% with a TEP-style model is contained in the incomplete TF binding site domain profile collection presently available even in a well-characterized model species such as *Arabidopsis*. It is certainly possible that an unconsidered influence such as DNA methylation status plays a role, although this seems largely correlative with chromatin status and not necessarily definitively causal. Finally, it is possible that important micro-scale chromatin accessibility patterns are not able to be well-captured at present by our model, given that the potentially relevant set of binding site locations and their associated chromatin state is enormous as compared to the set of highly expressed TSS locations in the genome; regularization algorithms do have limits on their ability to select the most predictive features from a vast sea uninformative values using a relatively small number of examples. Yet we see little indication of this ‘micro-scale chromatin accessibility pattern’ transcriptional control concept within the fairly broad regions of accessible chromatin in the *Arabidopsis* proximal promoter.

### Accurate TSSs implicate proximal cis-regulatory regions as primary determinants of tissue-specific gene expression

Our modeling outcomes strongly suggest that DNA sequence, within about 500 nt directly upstream of the TSS, is by far the most influential feature in successfully predicting tissue expression level differences, as opposed to distal chromatin status. Specifically, our study suggests a paradigm shift in the way we generally assume plant promoters to operate: for the vast majority of differentially expressed genes in developing *Arabidopsis* organs, it is the pattern of cis-regulatory sites in the TSS-proximal DNA of these regions, regardless of chromatin state, that is most explanatory of the tissue of expression. The presence of TF binding site Regions of Enrichment, and the ability to predict both TSS location and tissue of expression primarily from binding sites within these regions, underscore the important tissue specificity role of TF binding site patterns within the [TSS – 500 nt, TSS + 100 nt] ‘proximal’ promoter region in developing *Arabidopsis* seedlings.

It is surprising that TF binding site patterns in relatively accessible proximal promoter regions could largely dictate a gene’s tissue of expression, though there is a growing body of genome-scale evidence that this may well be the case in higher eukaryotes (Vandenbon and Nakai, 2010; Huminiecki and Horbańczuk, 2017; Chereji et al., 2019). Studies such as (Maher et al., 2018) emphasize specific groups of TFs that appear to act as ‘control modules’ within different plant tissues and cell types; it may be the case that a relatively small and distinct group of TF master regulators tends to work in-concert within each tissue to help orchestrate chromatin remodeling, ensuring that promoter regions are largely accessible in the important proximal locations.

Additionally, several studies suggest the intriguing possibility that distal enhancer regions may in fact be playing a significant role in tissue specific gene expression, but that Pol-II interactions with these enhancers are dictated to a high degree by proximal promoter sequences. The (Taher et al., 2013) study entitled “Sequence signatures extracted from proximal promoters can be used to predict distal enhancers” provides substantive computational evidence for this concept. (Ong and Corces, 2011) provides a literature synthesis of studies on enhancer function in tissue specific gene regulation, noting cumulative evidence that chromatin looping between enhancer and promoter regions is likely to be dictated at least in part by specific groups of TFs. Our study is consistent with the possibility that distal enhancers are indeed playing a substantial role, but are interacting with specific patterns of TFs which bind the proximal promoter to mediate chromatin looping.

### Implications for synthetic biology: systematic design of tissue-specific promoters

A recent study (Cai et al., 2020) strongly supports the concept that the specific locational arrangements of endogenous binding sites within a plant promoter can have a dramatic effect on overall expression level. The construction and outcomes from our Tissue-of-Expression-Prediction from Regions of Enrichment or “TEP-ROE” model carry two practical implications along these lines for additionally directing strong expression in one tissue vs another. Firstly, the TEP-ROE model identifies specific TSS-proximal TF binding site regions as important to differential gene expression in each tissue sample— in our study, developing *Arabidopsis* roots and shoots. Our gel-shift analysis provides plausible support for the idea that when these model-identified patterns of high-affinity TF sites are located upstream of a specific promoter, then these sites may be bound and functional, serving in the context of surrounding sequence to preferentially upregulate gene expression in a particular tissue. Secondly, when certain TF binding densities are given a zero-coefficient or ‘removed’ from a promoter, this can produce a large shift across the decision boundary, indicating a model prediction that removal of high-affinity sites in this region would change the tissue in which a gene expresses most strongly.

In other words, in-silico “knockouts” identify TF binding regions that alter the predicted tissue in which the gene is differentially expressed. There are hundreds of cases in which a single TF-region ‘knockout’ is predicted to cause such a shift, and thousands of cases in which a ‘double-knockout’ is predicted to cause such a shift. Taken together, these results suggest strong potential for tissue-specific promoter design. For this application, rather than focusing on differentially expressed genes, one would re-train a TEP-ROE model to classify the tissue of expression for genes that expressed very highly in one tissue and very little in the other, in terms of absolute transcript counts. This would allow identification of specific high-affinity TF binding sites that, when removed from the context of a certain promoter, change the tissue of expression for a gene entirely. In summary, our model presents the exciting possibility that tissue-specific synthetic promoters can be systematically constructed using endogenous cis-regulatory sites whose presence/absence in specific locations leads to a predicted shift in tissue of expression.

## Methods

### Plant materials and sample preparation

*Arabidopsis thaliana* ecotype Columbia 0 seeds were sterilized in a solution of 50% (v/v) bleach solution with 0.1% Tween 20 for 10 minutes, then rinsed extensively with sterile deionized water. Sterilized 100 micron nylon mesh was placed on top of solidified medium (30 mM sucrose, 4.2 g Murashige and Skoog medium (PhytoTech Labs), and 0.8 % Phytagar, pH adjusted to 5.8 with KOH) in large petri plates (Genesee Scientific). Following a 4 day vernalization period in water, sterilized seeds were suspended in a 0.75% agar solution and were transferred to each plate in two dense rows (∼500 seeds per row) in a laminar flow hood under sterile conditions. Seedlings were grown vertically in a Conviron PGR15 growth chamber at 21°C under a 12:12 hour light:dark cycle (50% humidity, and 250 mol/m2/s light intensity). At seven days, the seedlings were harvested and divided into three batches. For each batch, seedlings were dissected using a surgical blade and the root tissue was separated from the shoot tissue. For our purposes here, shoot tissues include the hypocotyl, cotelydons, and any stems and true leaves that had developed by the time of collection. Each batch of seedlings was handled identically, and harvested tissues were flash-frozen in liquid nitrogen, then stored at ^-^80°C until needed for the following protocols.

### Dataset generation

#### Sequencing

TSS-Seq was performed for both root and shoot samples as described in (Cumbie et al., 2015b) using the nanoCAGE-XL protocol in conjunction with the HiSeq-2000 sequencing platform. For DNase-seq, chromatin from isolated nuclei was digested with DNase I and libraries were prepared for both root and shoot samples according to the DNase-I-SIM protocol (Filichkin and Megraw) as published in (Cumbie et al., 2015a). RNA was isolated from both root and shoot samples for RNA-Seq using the RNeasy kit (Qiagen). Samples were analyzed for quality on the Bioanalyzer 2100 (Agilent) and only RNA with a RIN > 9.0 was used. Single-end libraries were sequenced on the Illumina HiSeq-2000 sequencing platform in triplicate.

#### Read preprocessing and alignment

CapFilter software (Cumbie et al., 2015b) was used to pre-process all nanoCAGE-XL TSS sequence files prior to alignment, removing library artifacts such as extra guanines at the beginning of the reads. For all three sequencing experiments, single-end reads were aligned to the TAIR10 reference genome (Lamesch et al., 2012), using Bowtie version 2.0 (Langmead and Salzberg, 2012) with the parameter settings ‘-v 0 -m 1 -a best strata’ (uniquely mapped reads with only one mismatch allowed).

#### nanoCAGE-XL TSS-Seq data processing

After cap-filtering and aligning the TSS-Seq reads for the root and shoot samples, the JAMM peak finder (Ibrahim et al., 2015) was used to identify TSS read clusters. For TSS datasets in our study, the fragment size and bin size were both set to 10. The output of JAMM is a list of peaks along with their genomic coordinates. The coverage subcommand from the bedtools software suite (Quinlan and Hall, 2010) was used with the parameter settings -s (requiring same-strandedness) and -d (for reporting depth at each position) to retrieve the number of aligned reads in each peak region for peak annotation. An R script was developed to process the aligned reads within peak regions and generate peak information, such as the number of aligned reads in peak, TSS peak mode location, and mode read count. Each TSS peak was assigned to the closest TAIR10 annotated transcript, and peaks which fell within 250 bps upstream of the annotated translation start site and contained more than 50 read counts were selected for use.

#### DNase I SIM data processing

After alignment to the genome, the F-Seq peak-calling software (Boyle et al., 2008) was used to identify DNase-I hypersensitive sites (DHSs), as in (Cumbie et al., 2015a). We chose this OC-Seq peak caller because it provides compatible output with the original DNASE-I ENCODE data, and is therefore comparable with DNase-I peak usage by the machine learning modeling studies discussed in the Introduction and Discussion sections. Subsequent peak callers have more parameters that can be tuned, but all peak callers are smoothing algorithms that have limitations and tradeoffs in signal processing parameter selection. F-Seq was run with a specified feature length of 300 and a minimum DHS length of 50 nt.

#### RNA-Seq data processing and Differential Expression Analysis

Individual transcript abundance was determined using the RSEM software package (Li and Dewey, 2011) for each root and shoot RNA-seq sample. RSEM enables accurate transcript quantification using its built-in bowtie2 alignment by building the reference sequence from a user-provided genome annotation and calculating expression for each isoform. We used *rsem-prepare-reference* and *rsem-calculate-expression* for preparing reference sequence and computing transcript abundances, respectively. We then used EBseq (Leng et al., 2013), which is included in the RSEM package and is robust to outliers, in order to detect differentially expressed transcripts (*rsem-run-ebseq*).

### TSS-Seq data quality analysis

#### Sequence depth analysis

To determine whether our sampling depth was sufficient to accurately represent gene expression in root and shoot samples, we performed saturation analysis on our nanoCAGE-XL data. This analysis was performed as described in (Morton et al., 2014). Supplementary Figure 1 contains the results of this analysis.

### 3PEAT-style Models

#### Model construction

Using our nanoCAGE-XL peak data, we constructed models to predict whether a given genomic location is a TSS for each tissue type. Starting with annotated peaks, we removed those with less than 100 reads per peak or less than 30 reads at the peakmode, and kept only those peaks labeled as being a TSS, within the 5’UTR region, or within 500 nucleotides of the TSS. We then generated the model features from TAIR10 sequences surrounding these peak regions using the TFBScanner (Morton and Megraw, 2014). The filtered peaks were then used to train and test 3PEAT models exactly as described in (Morton et al., 2014).

#### Functional Binding Site Selection

The TSBS sequence features, their model-assigned weights, and the TSS probabilities generated from the root-trained model were then used to construct a table of putative functional binding sites and their associated metrics. This table was then filtered to retain only Narrow Peak promoters. Next, the data was sorted by model output probability, descending total feature score, descending model weight, descending false negative rate, ascending false positive rate, peak count read, and the absolute value of relative location. TFBS sites were selected from the top 500 rows of this sorted table for wet-lab validation of functional binding.

#### EMSA

Nuclear extracts were purified from *Arabisodpsis thaliana* Columbia 0 roots following a slightly modified protocol from (Staiger et al., 1991). We spun our cellular lysates at 2200 x *g* for 1 minute at 4°C before passing the supernatant through a series of progressively finer meshes (100 micron, 60 micron, 30 micron). Nuclei were washed and pelleted at 2200 x *g* for 10 minutes at 4°C. Halt Protease Inhibitor Cocktail (Thermo Fisher Scientific) was used in place of the KCl in Buffer B. After dialysis, samples were prepared with the Qubit Protein Assay Kit (Invitrogen) per manufacturer instructions and total protein concentration was measured on a Qubit Fluorometer (Invitrogen). DNA probes were designed using selected PWMs and the flanking sequences from the associated TSS. Both the template strand and its reverse complement were labeled using the Pierce Biotin 3’ End Labeling Kit (Thermo Fisher Scientific) and the complementary oligonucleotides were annealed. EMSAs were performed using the LightShift Chemiluminescent EMSA kit (Thermo Fisher Scientific). Briefly, 15 μL of nuclear extract (containing 4 μg total protein) was incubated together at room temperature for 30 min with 80fmol of biotin-labeled probes in 25 μL reaction mixtures containing 1X Binding Buffer, 10 mM DTT, 40 ng/μL poly(dI-dC), and 2% (v/v) glycerol and then separated on 6% native polyacrylamide gels in Tris-borate-EDTA buffer containing 45 mM Tris, 45 mM boric acid, and 1 mM EDTA, pH 8.3. Unlabeled probe was used as cold competitor in 300X excess. Labeled probes were detected using the Pierce Chemiluminescence Detection Kit (Thermo Fisher Scientific) according to manufacturer instructions and visualized on an Azure c600 imager (Azure Biosystems).

#### Evaluation of Peak Classification in Same-Tissue and Other-Tissue Datasets

Two 3PEAT style models were constructed—one trained on 80% of the nanoCAGE-XL peaks from root and the other on 80% of the peaks from shoot. Models were tested on the other 20% of the peaks from both the same tissue and the opposite tissue. Then, we determined which peaks were misclassified by the models. For each test set, we divided the peaks which were classified incorrectly into four groups: “root-misclassified” (misclassified only by the root-trained model), “shoot-misclassified” (misclassified only by the shoot-trained model), “both-misclassified” (misclassified by both models), and “any-misclassified” (includes all peaks misclassified by at least one of the models). For each test-set-model pair, we compiled a list of genes that had at least one TSS peak which was misclassified by only that model. We observed that these lists greatly overlapped with the other list of their respective tissue-type, therefore we chose to exclude genes that were common to both lists. For each list of genes described above, we performed a GO enrichment analysis using GOATOOLS (Klopfenstein et al., 2018), a Python-based automated gene ontology enrichment analyzer. We used the genes represented in the full test set pertaining to the list as a population for comparison, limiting the scope of the analysis to “Biological Process” ontology terms, and limiting results to those with p < 0.05. Additionally, a simple Plant-Ontology enrichment analysis (Jaiswal et al., 2005) was performed by creating a list of terms subordinate to each of the terms of interest, “root system” (PO:0025025), and “shoot system” (PO: 0009006). Genes with peaks misclassified exclusively by one of the models, which were annotated with terms subordinate to one of the terms of interest were considered to be matches to those terms. A Fisher’s exact test was performed comparing the proportion of matches in genes misclassified by one model exclusively vs. genes misclassified by either model, with a cutoff of p < 0.05 for significant enrichment or depletion.

### Tissue of Expression Prediction Modelling

### Construction of TEP models

#### Feature Generation

Each TFBS feature represents an approximation of cumulative binding affinity that a particular TF has for a specific genomic region. Each open chromatin feature represents a percentage of nucleotides within the associated region that are open. In the model versions that used TFBS features, each ROE region as determined in previous section is divided into five overlapping sub-widows and two flanking windows as described in (Morton et al., 2014). The TFBS features are cumulative log-likelihood scores for each ROE sub-window on the same and opposite strands (as the gene in consideration). The chromatin features are computed as the percentage overlap between the open regions and each ROE sub-window. Only log-likelihood scores greater than zero are considered as potential binding sites and contribute to the sum, therefore the minimum value for a TFBS feature is 0 (this is a case where none of the nucleotides in the region represent a potential binding site with a greater-than-zero log-likelihood score). In the Tiled model, the entire region from 1 kb upstream to 500bp downstream of the TSS mode is divided into non-overlapping windows of 100bp in width. The TFBS features are computed as cumulative log-likelihood scores within each tile for both strands. The open chromatin features for Tiled model are computed as a percentage overlap between the open regions and each tile. In addition to TFBS and chromatin features, we added sequence content features such as GCcontent (CG% within 100 bp upstream of TSS mode), CAcontent (CA% within 100 bp upstream of TSS mode), GAcontent (GA% within 100 bp upstream of TSS mode), ATcontent, and general promoter openness. ATcontent features were computed for each 20bp tiles within −200 to +40 bps from the TSS mode location.

##### POSITIONAL WEIGHT MATRIX SET

Position Weight Matrices (PWMs) for TFs in *Arabidopsis thaliana* downloaded from TRANSFAC (Wingender, 2008), JASPAR (Bryne et al., 2008), AGRIS (Davuluri et al., 2003), and CIS-BP (Weirauch et al., 2014) databases. We developed a software program in Python in order to compute the element-wise distance between PWM pairs; pairs with a distance less than or equal to our empirical determined threshold of 0.9 were determined to be redundant.

##### REGIONS OF ENRICHMENT AND TF SELECTION

TSS peaks identified as described above in root and shoot were collected (approximately 50,000 peaks) and 6 kb sequences were extracted (TSS - 3 kb, TSS + 3 kb, centered at each TSS mode) from the TAIR10 reference genome. As with the TSS-prediction models described earlier in the methods, the TFBS Scanner suite (Morton and Megraw, 2014; Morton et al., 2014) was used to scan each PWM over the extracted sequences, computing log-likelihood scores over these regions. Regions of enrichment (ROEs) were defined on both the forward and reverse strands by identifying the highest scoring region (the region with the largest sum of positive log-likelihood scores) for each PWM across all promoter examples (Morton et al., 2014). PWMs with cumulative log-likelihood score peaks up to 1 kb from the TSS mode were considered as regions of enrichment for that PWM. The ROEs for each PWM were computed using an updated version of ROEFinder software written in R. Our TEP-ROE model construction process is detailed in Supplementary Figure 19.

##### PROMOTER TILING

Using TSS peaks identified as described in previous sections, 6 kb sequences (TSS - 3 kb, TSS + 3 kb, centered at each TSS mode) were extracted from the TAIR10 reference genome. Sequences located from 1000 bp upstream of TSS mode up to 500 bp downstream of the TSS mode were divided into 100-nt-wide, non-overlapping tiles. The PWMs were then scanned over each tile to compute cumulative TFBS log-likelihood scores and percent overlap with open chromatin region in root and shoot tissues. Our TEP-Tiled model construction is detailed in Supplementary Figure 20.

##### FEATURE SCALING

Open chromatin features, as described in the feature generation section, all share the same 0-1 range and are interpretable as an “openness proportion” without modification. TFBS features, however, are computed as a sum of positive log-likelihood scores over all nucleotides in the region, where each nucleotide is taken as the starting point of a potential TF binding site; the log-likelihood score is computed at this site using (1) the PWM associated with the TFs binding domain, and (2) a local background nucleotide distribution model. The maximum possible value for a TFBS feature is region length multiplied by the maximum possible log-likelihood value PWM_scoreMax_ of any binding site (i.e. the score of the PWMs consensus sequence); this value represents a theoretical ‘maximal’ case in which every nucleotide in a region represents the consensus sequence of the PWM. To normalize TFBS features such that each feature conceptually approximates a proportion of the maximum binding affinity, each TFBS feature is divided by its region length. This puts the mean TFBS feature value on the same order of magnitude as the mean OC feature value and allows for comparison between feature regions of unequal length.

#### Model Training and Testing using nanoCAGE-XL TSSs

We constructed two classes of TSSs for use in training and testing of the ROE and Tiled models. The “root” class (class 0) consists of TSSs associated with transcripts that are strongly expressed in roots as compared to shoot, and the “shoot” class (class 1) consists of TSSs associated with transcripts that are strongly expressed in shoots as compared with roots. For our purposes, we defined “strongly expressed” as having an RNA-Seq data log_2_ fold-change value greater than 3. Additionally, TSSs present in both classes were screened to ensure that each TSS peak contained at least 300 reads in the tissue of its class label and each TSS-associated transcript had an RNA-Seq expression value in both roots and shoots of at least 30 TPM. This ensured that both values used for fold-change comparisons were reliably above background noise (Zavolan, 2015). The remaining TSS peaks were then randomly partitioned into 80% training and 20% independent held-out test sets. Each data set contains balanced number of labeled classes. The Python Scikit-learn library (Pedregosa et al., 2011) was used to implement L1-regularized logistic regression. L1-model weights were tuned on the training set with 5-fold cross-validation (Supplementary Figure 8), during which a range of parameter values was examined on the test partition. The average of parameter values resulting in the best performance across the folds was selected for final testing on the independent held-out test set (2.67 for TEP-ROE and 2.29 for TEP-Tiled). All auROC and auPRC values are reported on the independent held-out test set for each model.

### Tissue of Expression Modeling Analyses

#### Model Stability Assessment

In order to evaluate the stability of our TEP models, we performed two types of assessments. First, we re-ran our TEP models 30 times on training and test sets with randomized 80/20 partitioning. For our second assessment, we removed the top N PWMs from our feature list (n = 5, 25, 45, etc). We used the new feature sets to re-train and test the model. The results from both of these assessments can be found in Supplementary Figures 10, 11, 12 and 13.

#### Comparison to model using TAIR10 annotated TSSs

To investigate the importance of using precise, experimentally-obtained TSS locations in modeling, we generated a feature set as described above in the Feature Generation section of the Methods using only annotated TAIR10 TSS locations. We then trained the TEP-ROE and TEP-Tiled models on these annotated TSSs on nanoCAGE-XL data, as described in the Model Training and Testing Methods section above.

#### Hard-coded promoter analyses

For our final TEP models constructed using nanoCAGE-XL data (TEP-ROE and TEP-Tiled, including all feature types), promoter examples which were correctly classified with a high probability (≥0.9) were selected for our “hard-coded promoter” analysis. The sets of TFBS and chromatin feature values for each of these promoters were extracted, and for each set the following formula was applied:

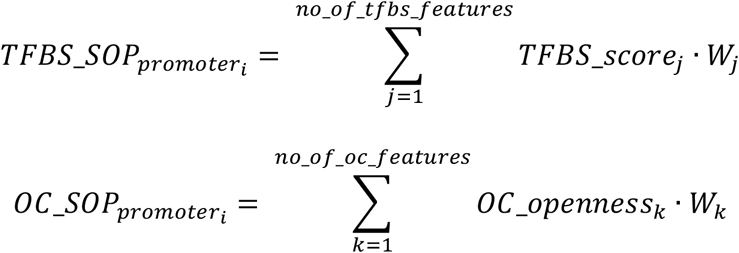

*W* is the model weight vector for each feature after training, *j* and *k* are the number of TFBS features and OC features, respectively, and *i* is the number of promoters. TFBS_SOP and OC_SOP are the sum of products for TFBS features and OC features, respectively. The distribution of the sum of products was computed for TFBS features and for chromatin features, and the 5% tails of these distributions were considered for detecting putative hard-coded promoters (Supplementary Figure 21). Promoters for which TFBS_SOP fell above the 95th percentile and OC_SOP fell below the 5th percentile were labeled as putatively hard-coded. Additionally, we extracted the features with the largest products for each of the TSSs for further investigation. GO-enrichment analysis compared to the entire genome was performed for the genes with putatively hard-coded promoters using GOATOOLS (Klopfenstein et al., 2018).

#### In silico knockouts

The output of the classifier function for our trained L1-regularized logistic regression models is a probability between zero and one, which represents the predicted likelihood of differential expression in the two tissue types. Probabilities greater than 0.5 are labeled as “class 1” or “shoot”, and probabilities less than 0.5 are labeled as “class 0” or “root”. Logistic regression is a Generalized Linear Model, where the probability of belonging to class 1 is a function of sum of products of feature weights by feature values as follows:

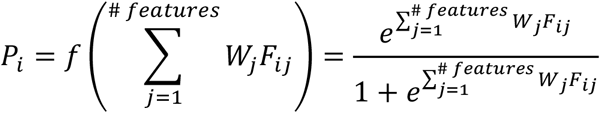

where *Pi* is the probability that Promoter *i* belongs to class 1 (shoot), W*j* is model weight, and F*ij* is the *j*th feature value for promoter *i*. Negative values of the feature product sum yield P_*i*_ (shoot) < 0.5 (a root classification), and positive values of this feature product sum yield P_*i*_ (shoot) > 0.5 (a shoot classification). Our *in silico* knockout process “zeroes out” selected feature values and then computes the new model-predicted probability for a promoter. In the “single-knockout” experiment, only one feature was removed from the equation at a time, in order to determine effect on probability outcome for every promoter. Supplementary Tables 13 and 14 report cases with the largest probability ‘shifts’ across the 0.5 decision boundary, indicating predicted high-probability ‘flips’ in tissue of greater expression upon *in silico* knockout of a single TF-feature region.

## Data and Model Availability

The full dataset of mapped, annotated nanoCAGE-XL TSS peaks, DNase I SIM peaks, and RNA-Seq expression levels for root and shoot samples in our study is available on GBrowse at http://megraw.cgrb.oregonstate.edu/suppmats/TissueOfExpressionPredictionDatasets. All raw datasets, processed datasets, and model coefficient files are also made available for download. Model training and evaluation pipelines are available upon request; these are designed to run on a Sun Grid Engine computing cluster, and require user familiarity with the Unix/Linux operating system, Make, Java, R, and Python; the pipelines cannot be supported by the authors on other hardware systems.

## Accession Numbers

All raw reads have been deposited in the National Center for Biotechnology Information Sequence Read Archive repository under the following accession numbers: OC-Seq (DNase I SIM) – PRJNA285928; TSS-Seq (nanoCAGE-XL) – PRJNA658605; RNA-Seq – PRJNA658596.

## Supplementary Data

**The following materials are available in the online version of this article**.

**Supplementary Figure 1: TSS-Seq root and shoot sampled read depth saturation analysis**

**Supplementary Figure 2: Chromatin accessibility surrounding shared TSS mode in root and shoot**

**Supplementary Figure 3: Proportion of top features shared by 3PEAT root and shoot models**

**Supplementary Figure 4: False positive and false negative rates for 3PEAT root and shoot models**

**Supplementary Figure 5: EMSA evaluations of putatively functional binding sites**

**Supplementary Figure 6: TEP-ROE model performance**

**Supplementary Figure 7: TEP-Tiled model performance**

**Supplementary Figure 8: Cross-validation ROC curves for ROE and Tiled models**

**Supplementary Figure 9: TEP model performance comparison**

**Supplementary Figure 10: Feature rank variability TEP-ROE model**

**Supplementary Figure 11: Feature rank variability TEP-Tiled model**

**Supplementary Figure 12: Feature removal performance plot for TEP-ROE model**

**Supplementary Figure 13: Feature removal performance plot for TEP-Tiled model**

**Supplementary Figure 14: Top-weighted feature comparison between TEP-Tiled and enhancer model**

**Supplementary Figure 15: TEP-ROE rank correlation plot**

**Supplementary Figure 16: TEP-Tiled rank correlation plot**

**Supplementary Figure 17: ROE feature products vs openness**

**Supplementary Figure 18: Tiled feature products vs openness**

**Supplementary Figure 19: TEP-ROE model construction**

**Supplementary Figure 20: TEP-Tiled model construction**

**Supplementary Figure 21: Feature product sums for hard-codedness evaluation**

**Supplementary Table 1: Number of TSS peaks and their mapped locations**

**Supplementary Table 2: Number of TSS peaks and covered number of transcripts**

**Supplementary Table 3: Basic statistics on TSS-seq, RNA-seq and OC data**

**Supplementary Table 4: Basic RNA-seq expression statistics**

**Supplementary Table 5: Top weighted features for 3PEAT root and shoot models**

**Supplementary Table 6: Cross-tissue model performance**

**Supplementary Table 7: Cross-tissue 3PEAT-style GO-analysis for misclassified TSSs**

**Supplementary Table 8: TEP-Tiled vs TEP-ROE models top feature comparison**

**Supplementary Table 9: Top 30 features of enhancer-covering Tiled model**

**Supplementary Table 10: Putatively “hardcoded” promoter examples from TEP-ROE**

**Supplementary Table 11: Putatively “hardcoded” promoter examples from TEP-Tiled**

**Supplementary Table 12: GO enrichment analyses for “hardcoded” promoter examples**

**Supplementary Table 13: “In-silico knockout” results for TEP-ROE model**

**Supplementary Table 14: “In-silico knockout” results for TEP-Tiled model**

**Supplemental Data Set 1: (3PEAT_Model_weights**.**xlsx) 3PEAT logistic regression coefficients for 3PEAT root and shoot TSS location prediction models**

**Supplemental Data Set 2: (TEP_Model_weights**.**xlsx) TEP logistic regression coefficients for ROE and Tiled Tissue-of-Expression-Prediction models**

## Acknowledgements

Data generation for the study was supported by an NIH K99-R00 Pathway to Independence Award GM097188 to M.M. Algorithm design and computational analysis for the study was supported by an NSF CAREER Award 1750698 to M.M.

## Author Contributions

MM designed the study. MI and SF carried out laboratory experiments for TSS-Seq, OC-Seq, and RNA-Seq dataset generation. MA performed algorithm implementation. MA and VF contributed to the design of data analysis experiments and to the evaluation of results. MA and VF worked together to select EMSA assay sites for examination, and VF carried out EMSA assays aided by OO. VF, RG, ZB, and SO contributed portions of the data analysis. OO and ZB contributed to data preparation for public distribution. MM, MA, and VF wrote the manuscript, all authors contributed to editing of the manuscript and approved the final version of the manuscript. We thank Dr. Uwe Ohler for ideas and discussions with M.M that inspired the conception of this study. We thank Dr. Ashok Prasad for his idea to investigate the existence of “hard-coded” promoters. We thank Dr. Jason Cumbie for his help in preliminary evaluation of dataset quality, Natalie Brewer for her help in the PWM literature search, and Jordan Holdaway and Teresa Tran for their help in tissue preparation for the study.

