## Supplementary_Materials for "Accurate transcription start sites enable mining for the cis-regulatory determinants of tissue specific gene expression"

### Supplementary Figures

#### Supplementary Figure 1: TSS-Seq root and shoot sampled read depth saturation analysis

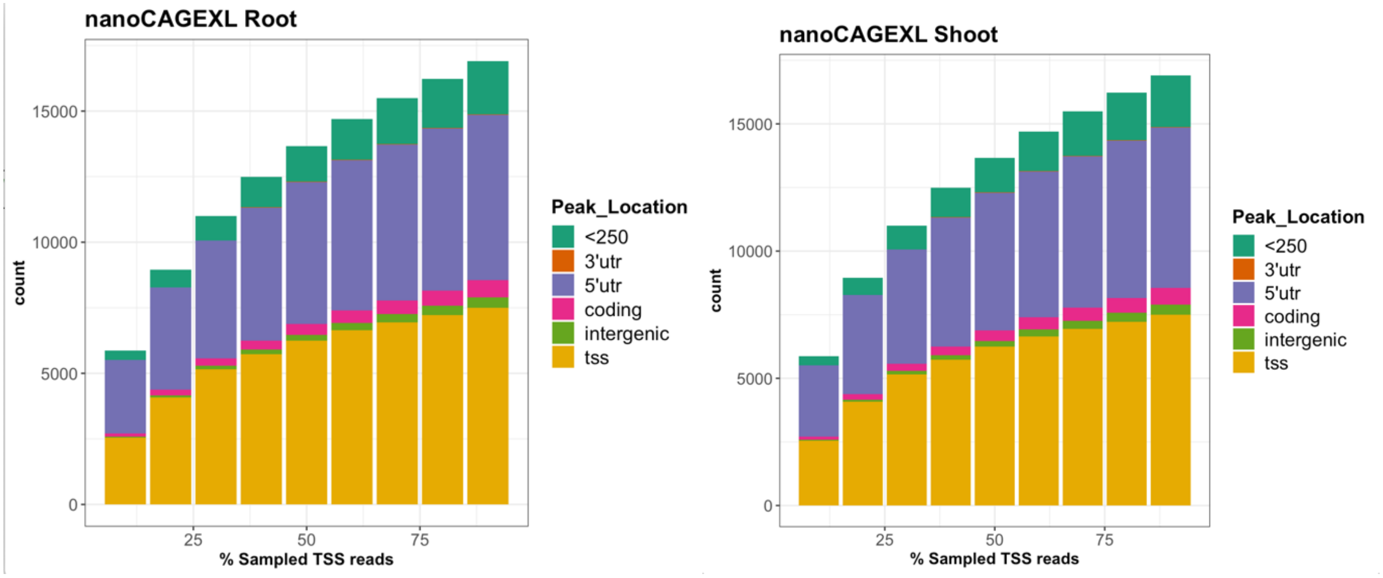

**Supplementary Figure 1:** Sequencing depth analysis of nanoCAGE-XL root and shoot data to determine whether sequencing depth accurately represented gene expression in our pooled root and shoot samples. Starting with stringently-mapping reads, reads were randomly sampled and peaks called according to our TSS peak calling procedure. As subsample size increases, fewer new genes are observed in each dataset. In each dataset, the change in number of genes has leveled off to nearly zero as the percentage of reads sampled increases from 90% to 100%. This indicates that the sampling depth we achieved in terms of gene coverage was within a few percent of the maximum sampling depth that could be achieved.

**Supplementary Figure 2: Chromatin accessibility surrounding shared TSS mode in root and shoot**

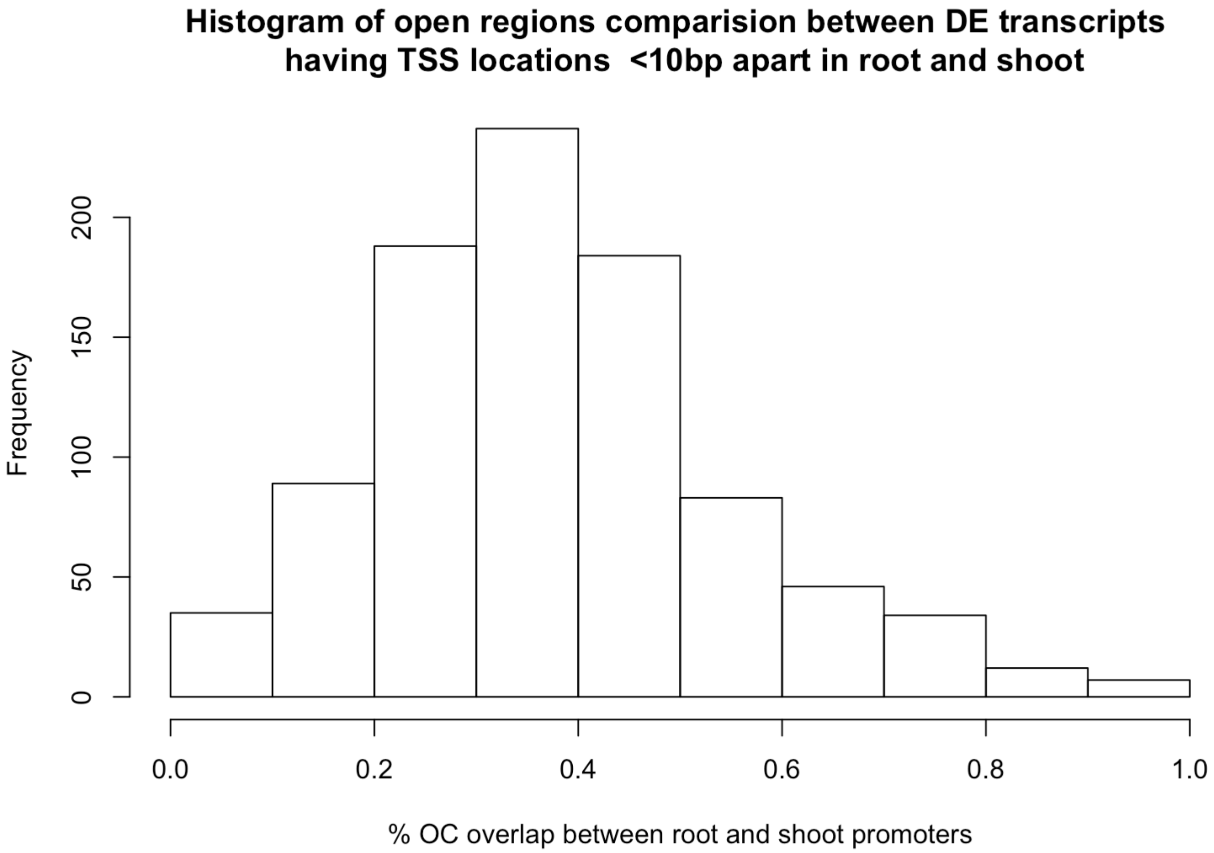

**Supplementary Figure 2:** Histogram showing the percent overlap in chromatin openness between the differentially expressed root and shoot transcripts with mapped TSS modes in close proximity (less than 10 nt apart). Percent overlap is computed as the percentage of nucleotides in the region surrounding each TSS [TSS - 3 kb, TSS + 3 kb] that agree in chromatin accessibility state (open vs closed).

**Supplementary Figure 3: Proportion of top features shared by 3PEAT root and shoot models**

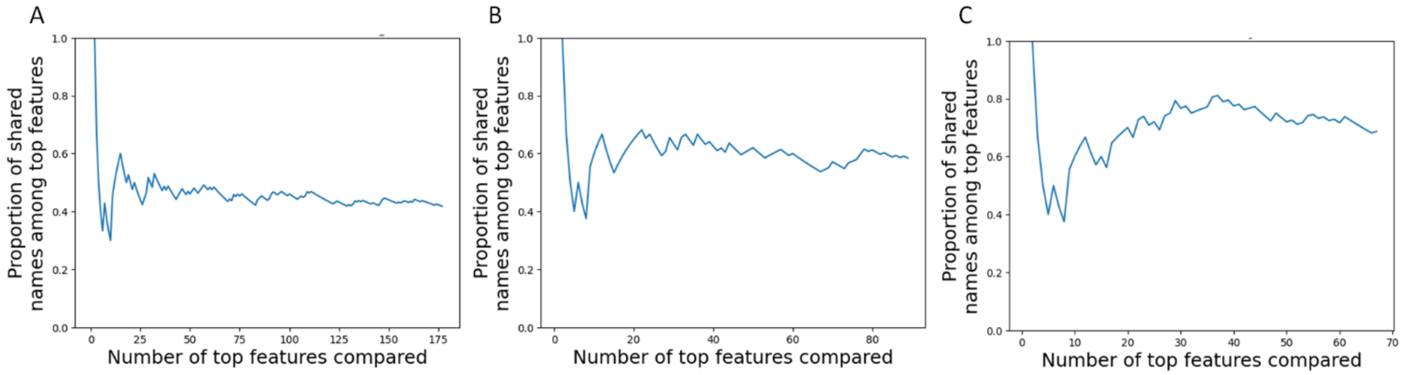

**Supplementary Figure 3:** Proportion of shared names in the top N features of both root and shoot 3PEAT models, calculated as $P = \frac{|TopN_{root} \cup TopN_{shoot}|}{N}$, i.e., the number of shared names among the top N names in both lists divided by N. A) Feature names including PWM (Positional Weight Matrix), strand and TSS. B) Feature names including PWM and strand. C) Feature names including PWM only.

**Supplementary Figure 4: False positive and false negative rates for 3PEAT root and shoot models**

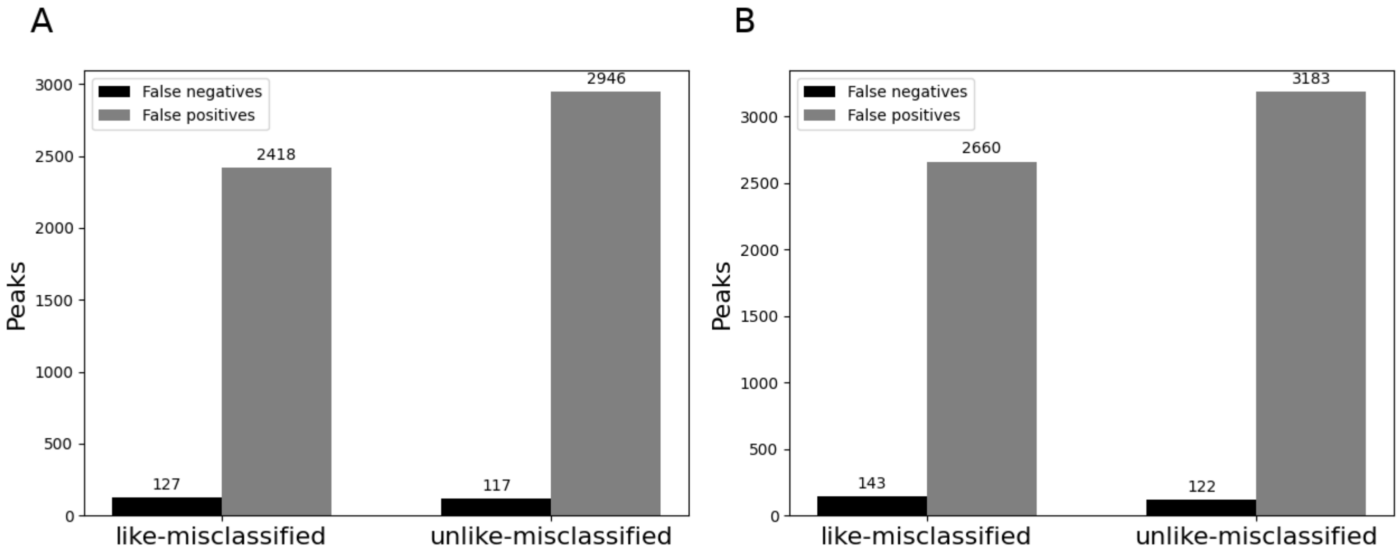

**Supplementary Figure 4:** The "like-misclassified" peaks are peaks that were misclassified by the model trained on the same tissue that the peaks were expressed in (root or shoot). The "unlike-misclassified" peaks were misclassified by the model trained on the other tissue. A) Peaks misclassified from the root test-set. B) Peaks misclassified from the shoot test-set model.

#### Supplementary Figure 5: EMSA evaluations of putatively functional binding sites

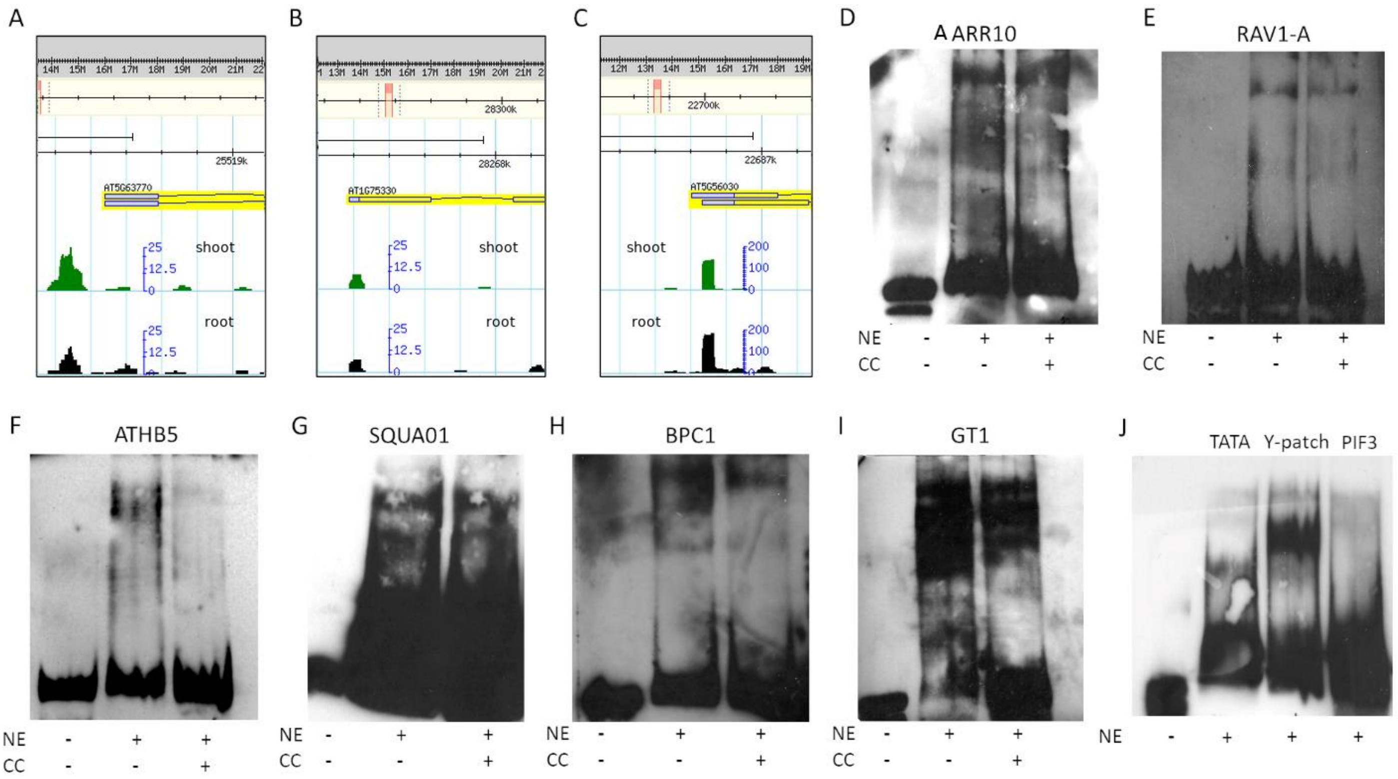

**Supplementary Figure 5:** A) GBrowse screenshot highlighting the small TSS-Seq peaks for DGK2. B) GBrowse screenshot highlighting the small TSS-Seq peaks for OTC. C) GBrowse screenshot of the large TSS-Seq peaks for HSP90.2, illustrating why we selected our second set of sites for testing from the promoter of this gene. D-I) EMSAs testing selected sites from the HSP90.2 promoter. NE = Nuclear Extract from 7-day old *Arabidopsis thaliana* Col0 roots; CC = cold competitor, >200X. J) EMSA showing binding of three sites. TATA from the HSP90.2 promoter and Y-Patch from the OTC promoter show a shift resulting from binding. PIF3 from the DGK2 promoter does not. The RNA-Seq abundance for PIF3 in the root sample was 0.1 TPM; TBP, however, had 45 TPM. No cold competitor was used for this particular gel.

#### Supplementary Figure 6: TEP-ROE model performance

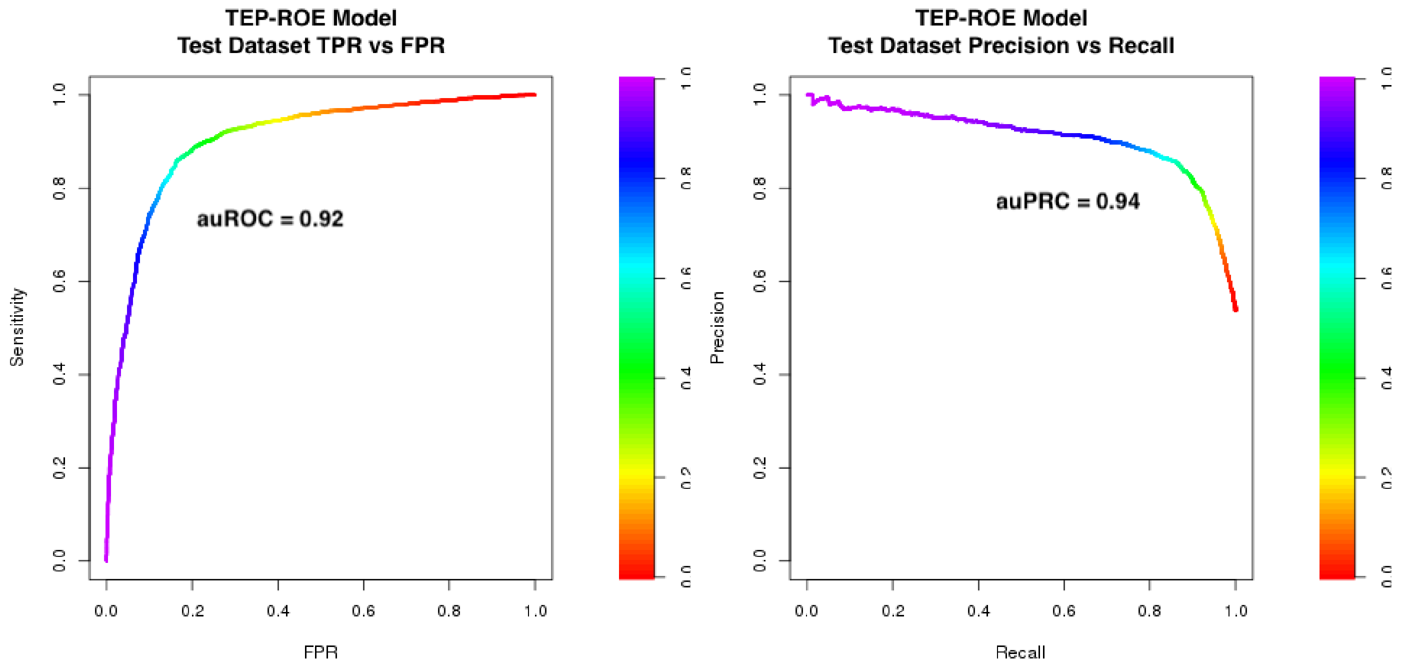

**Supplementary Figure 6:** Plots displaying true positive rate vs false positive rate (ROC) and precision vs recall (PRC) show the performance of the TEP-ROE model on an independent, held-out test set. The color gradient shows the probability threshold at each point on an auROC or auPRC curve (i.e. the probability threshold that produces the given FPR, TPR/Sensitivity values, or the given Recall, Precision values, at that point on the curve). The area under Sensitivity-Specificity curve indicates auROC performance value, and the area under Precision-Recall curve indicates the auPRC performance value. A 'perfect model' is associated with an auROC and auPRC equal to 1.0 (100% of the area is under the curve). A model that places examples into a class at-random will have an auROC and auPRC equal to 0.5 (50% of the area is under the curve).

#### Supplementary Figure 7: TEP-Tiled model performance

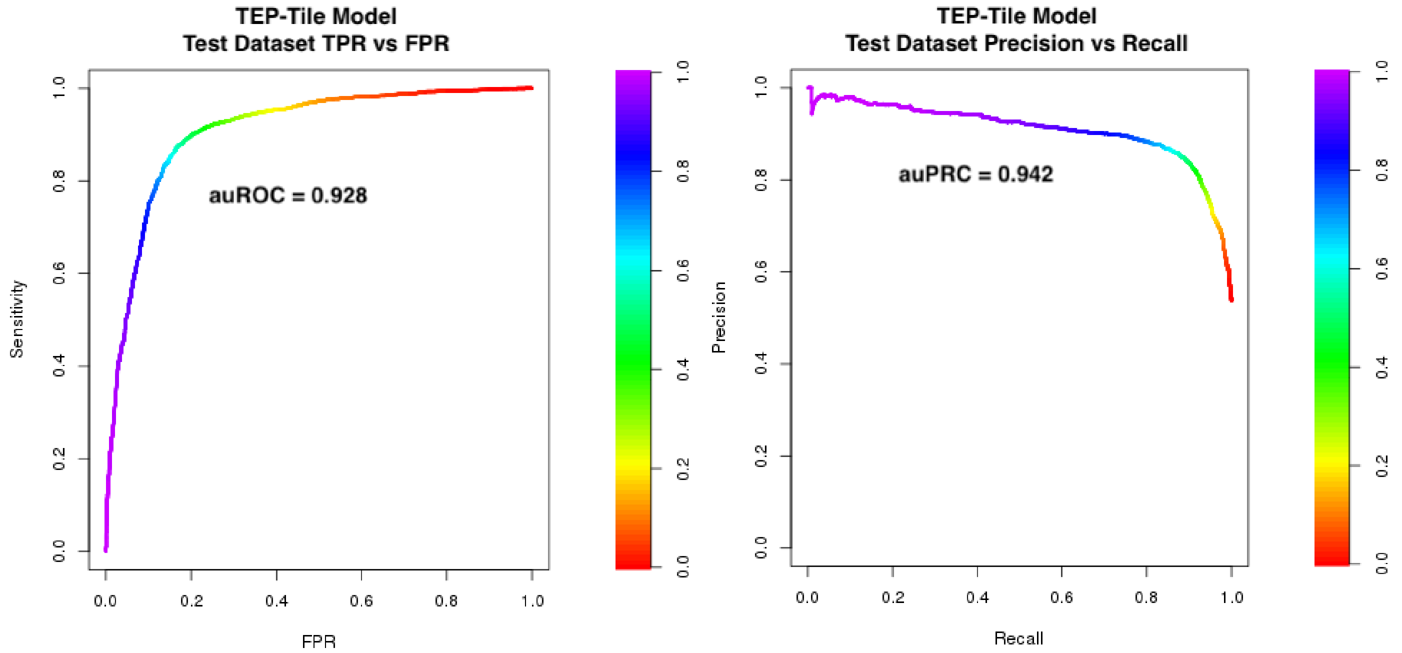

**Supplementary Figure 7:** Plots displaying true positive rate vs false positive rates (ROC) and precision vs recall (PRC) show the performance of the TEP-Tiled model on an independent, held-out test set. The color gradient shows the probability threshold at each point on an auROC or auPRC curve (i.e. the probability threshold that produces the given FPR, TPR/Sensitivity values, or the given Recall, Precision values, at that point on the curve). The area under Sensitivity-Specificity curve indicates auROC performance value, and the area under Precision-Recall curve indicates the auPRC performance value. A 'perfect model' is associated with an auROC and auPRC equal to 1.0 (100% of the area is under the curve). A model that places examples into a class at-random will have an auROC and auPRC equal to 0.5 (50% of the area is under the curve).

**Supplementary Figure 8: Cross-validation ROC curves for ROE and Tiled models**

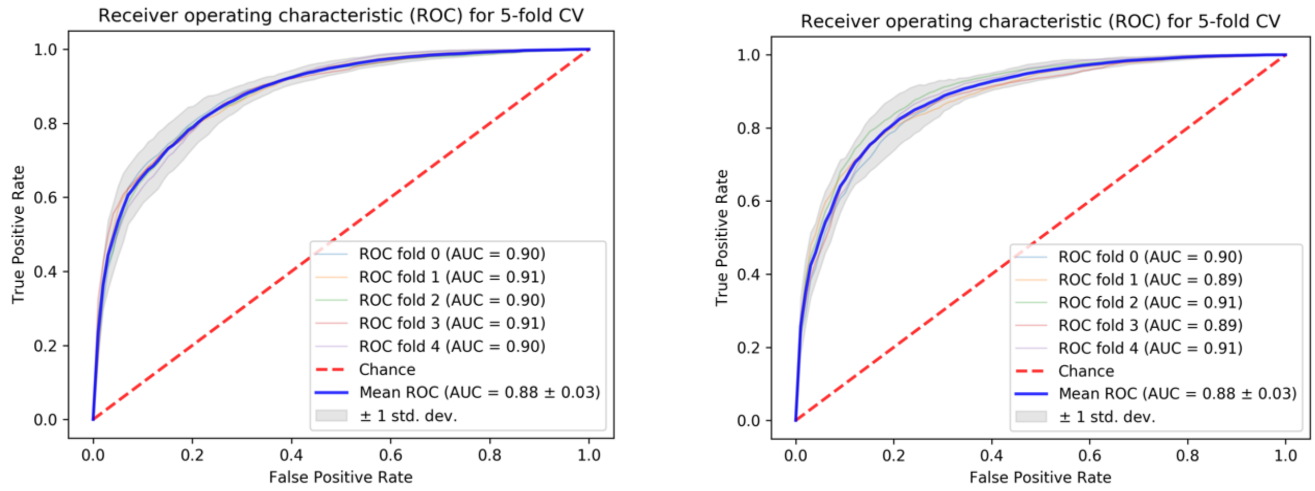

**Supplementary Figure 8:** 5-fold cross-validation was performed to determine the optimal regularization parameter for the TEP L1-regularized logistic regression models (Left: TEP-ROE, Right: TEP-Tiled). The training samples were divided into 5 partitions of equal size; a model was trained on 4 partitions, and auROC was computed on the 5$^{th}$ test partition over an L1 parameter range; the optimally performing L1 parameter was selection for this partitioning. This process was performed on all 5 possible partitionings. Plots show the auROC curve on each fold in the ROE model (Left) and the Tiled model (right). (The average of optimally performing L1 parameters over the 5 cross-validation partitions for each model was then used for the final model, trained on the entire cross-validation set and tested on the independent held-out test set for reporting of model performance results in the main manuscript).

#### Supplementary Figure 9: TEP model performance comparison

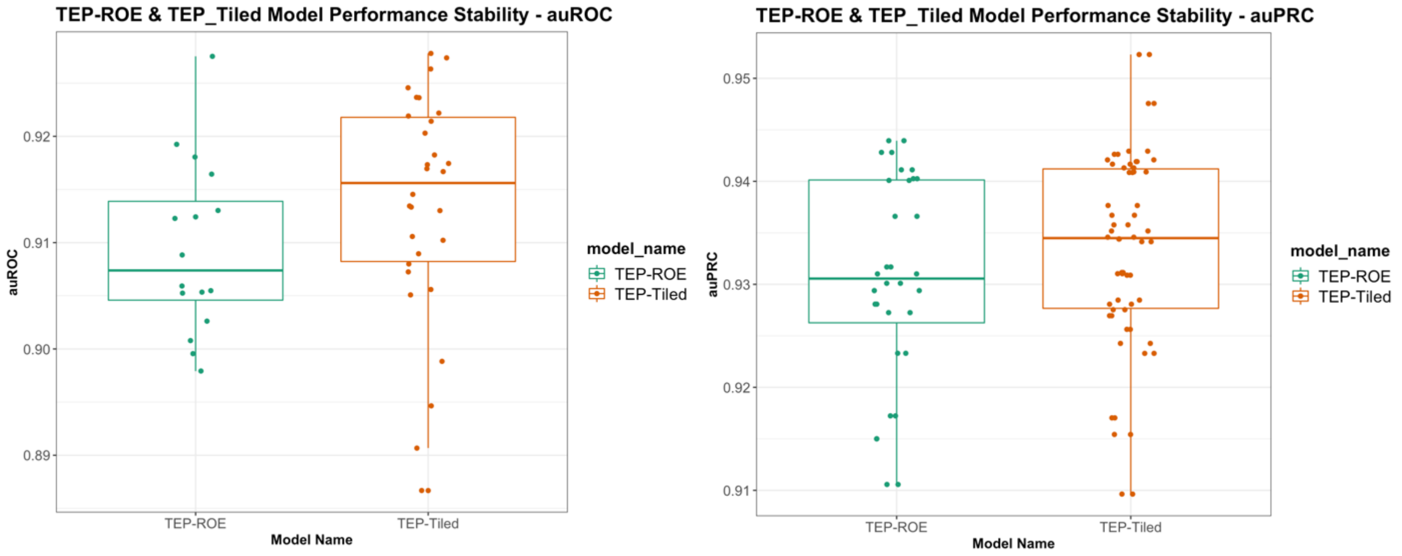

**Supplementary Figure 9:** Plots displaying auROC and auPRC for 30 model-runs each of the TEP-ROE and TEP-Tiled models. Training datasets were randomly partitioned 30 times and 5-fold cross-validation was performed, followed by testing on independent held-out samples. The auROC and auPRC were computed for each of 30 runs, for each model type. The TEP-Tiled model shows a slightly higher median auROC than the TEP_ROE model, however the variability in the Tiled model's performance is also higher— both in auROC and auPRC.

#### Supplementary Figure 10: Feature rank variability TEP-ROE model

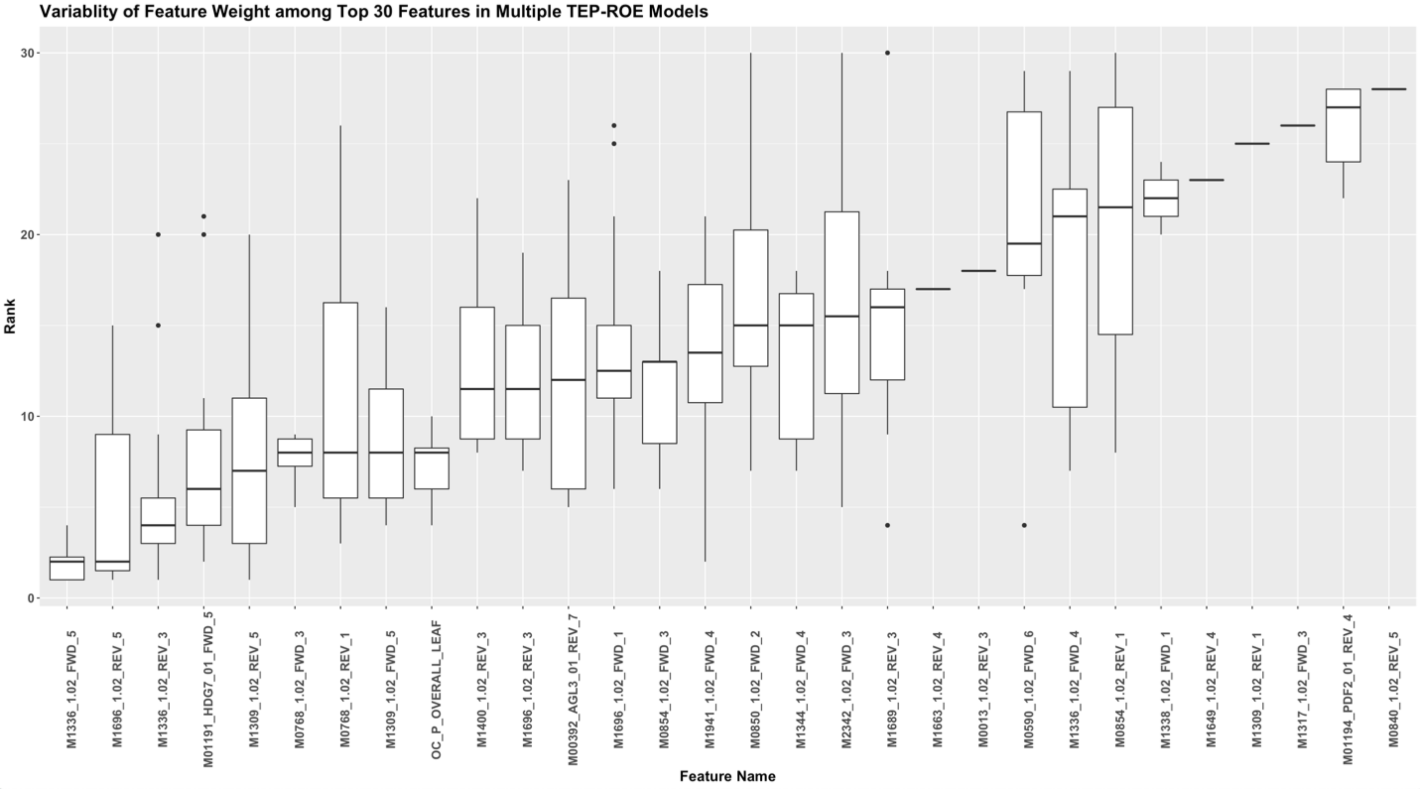

**Supplementary Figure 10:** Plot showing the variability in feature weight for the top 30 features over 30 runs of the TEP-ROE model. The top 30 most heavily weighted features from ROE model were considered, along with the rank of features in all 30 model-runs. This plot shows that some features display a substantially higher rank variability.

#### Supplementary Figure 11: Feature rank variability TEP-Tiled model

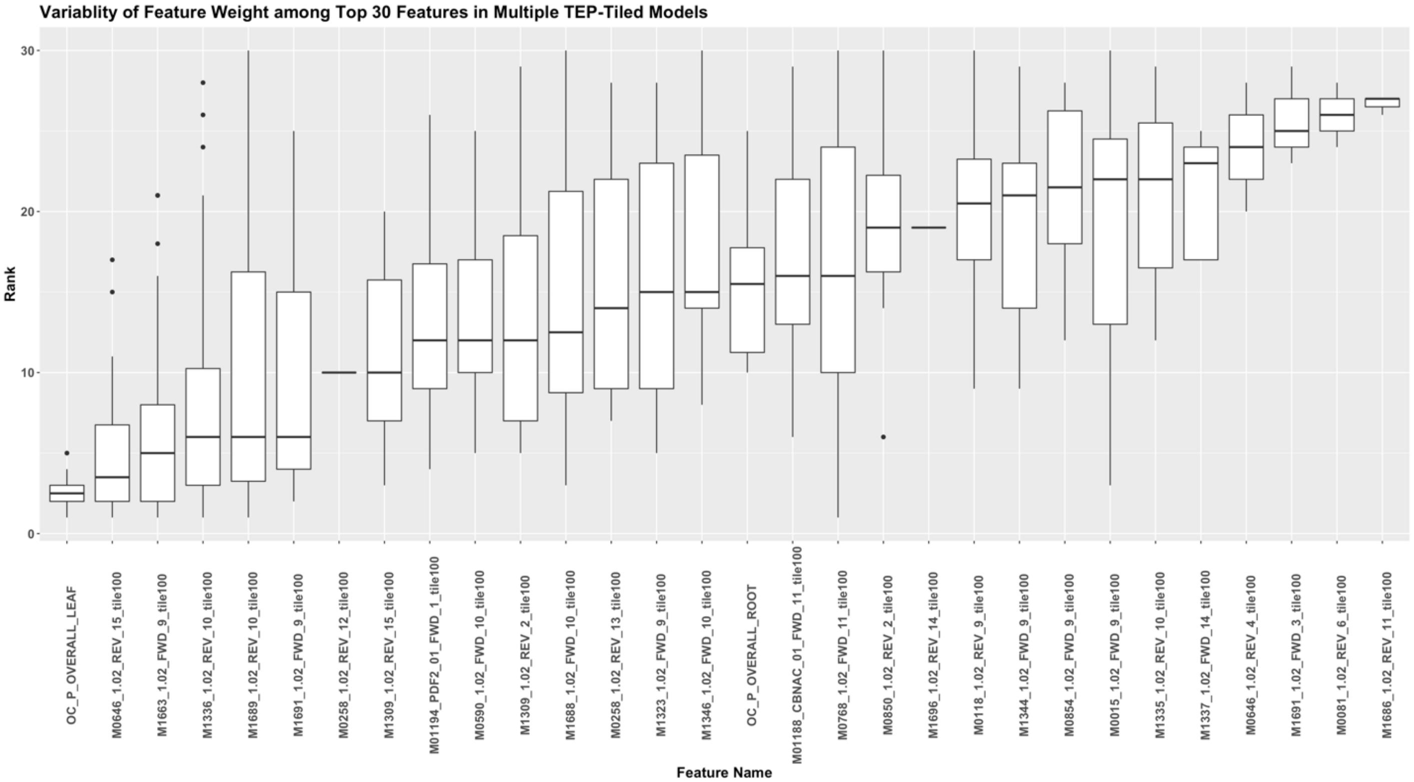

**Supplementary Figure 11:** Plot showing the variability in feature weight for top 30 features over 30 runs of the TEP-ROE model. The top 30 most heavily weighted features from tiled model were considered, along with the rank of features in all 30 model-runs. This plot shows that some features display a substantially higher rank variability.

#### Supplementary Figure 12: Feature removal performance plot for TEP-ROE model

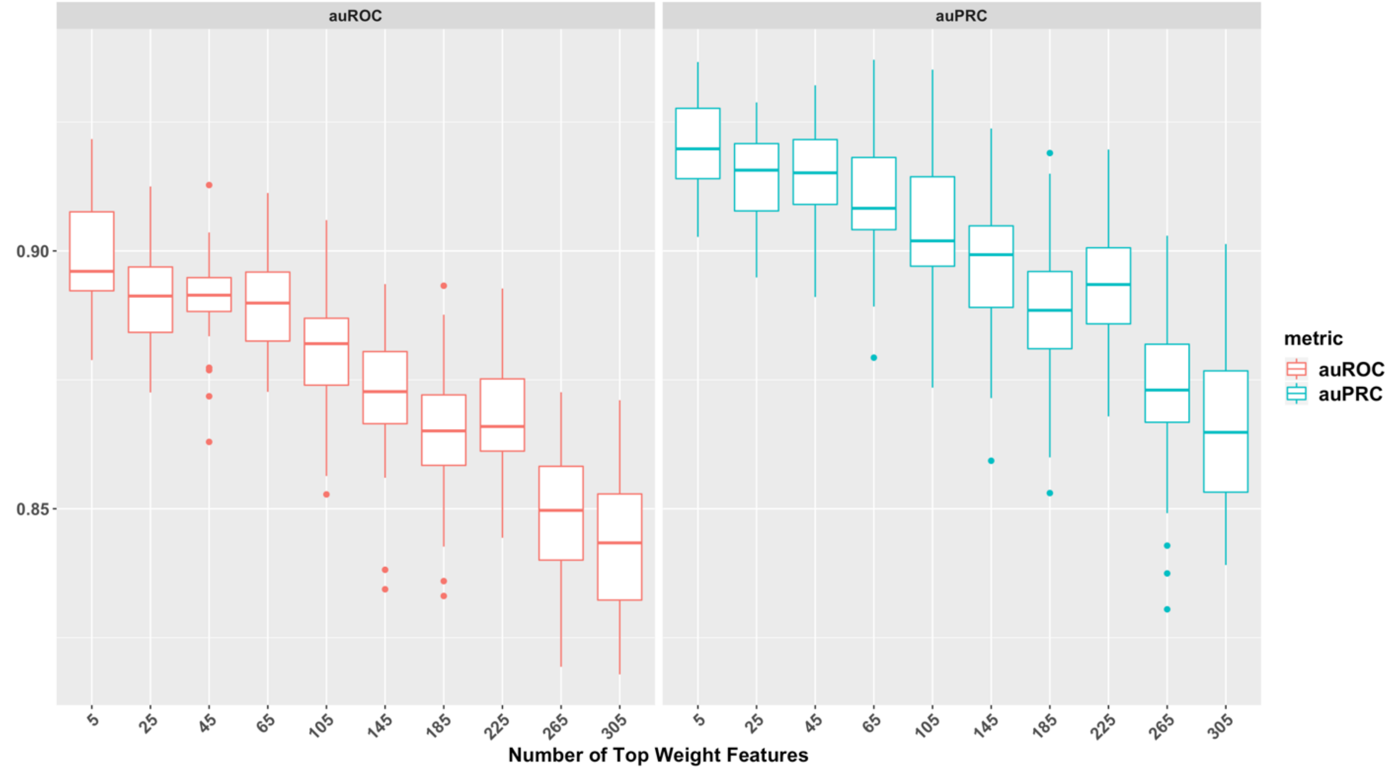

**Supplementary Figure 12:** Plots showing the drop in auROC and auPRC of the TEP-ROE model's performance after removing top weighted features from the training data. Sets of highly weighted PWMs (TF binding domain profiles) and their associated features were removed in multiple stages (top 5, 25, 45, .., 345) from training data. The TEP-ROE model's performance after removing each set of PWMs was recorded. The decrease in both auROC and auPRC indicates that the PWM sets are contributing unique information to the model, as other features do not appear to 'bubble up' to compensate for their removal in a way that maintains model performance.

**Supplementary Figure 13: Feature removal performance plot for TEP-Tiled model**

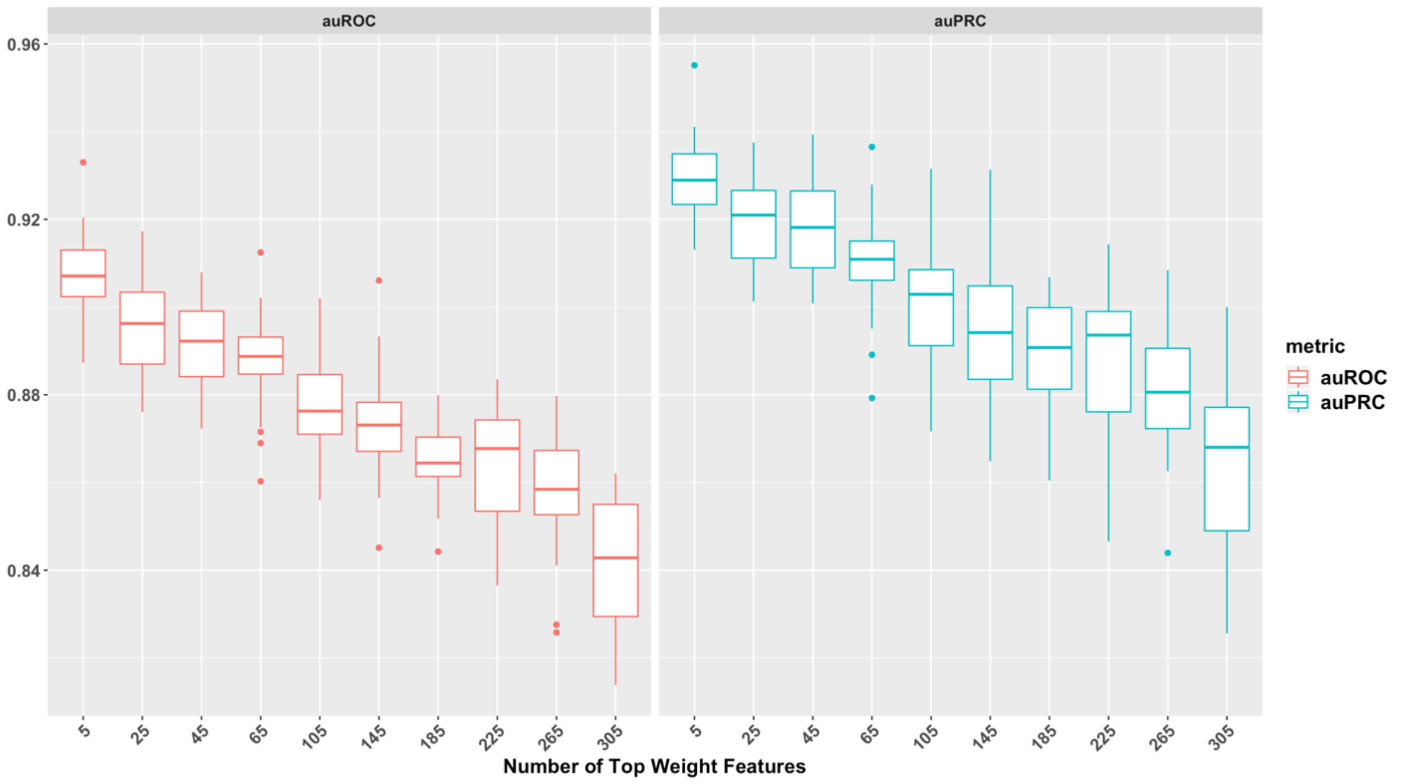

**Supplementary Figure 13:** Plots showing the drop in auROC and auPRC of the TEP-Tiled model's performance after removing top weighted features from the training data. Sets of highly weighted PWMs (TF binding domain profiles) and their associated features were removed in multiple stages (top 5, 25, 45, .., 345) from training data. The TEP-Tiled model's performance after removing each set of PWMs was recorded. The decrease in both auROC and auPRC indicates that the PWM sets are contributing unique information to the model, as other features do not appear to 'bubble up' to compensate for their removal in a way that maintains model performance.

**Supplementary Figure 14: Top-weighted feature comparison between TEP-Tiled and enhancer model**

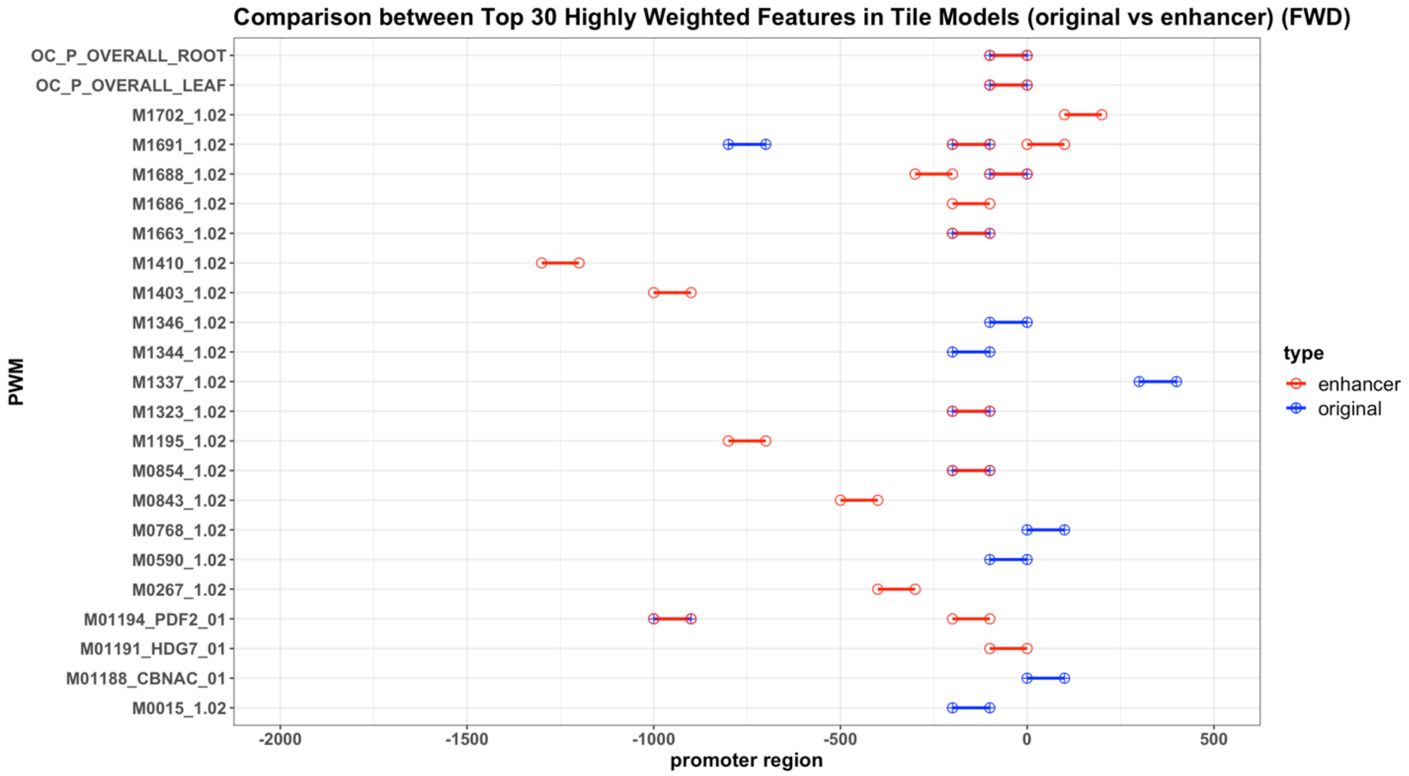

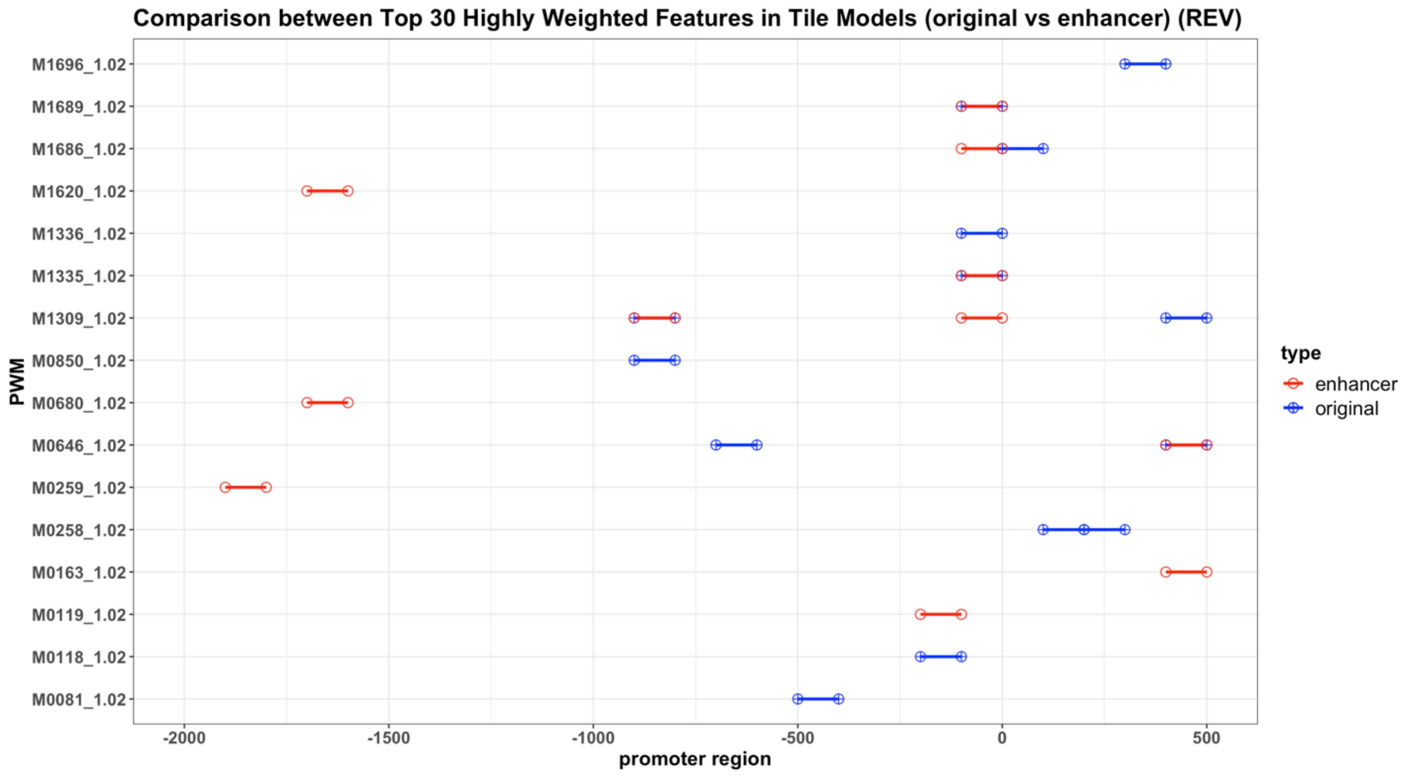

**Supplementary Figure 14:** Map of top-weighted feature comparison between the [TSS - 1 kb, TSS + 500 nt] Tiled model (TEP-Tiled, the "original model") and a [TSS - 2 kb, TSS + 500 nt] version of the Tiled model ("enhancer model”). Features in the TEP-Tiled original model are shown in blue, and in the enhancer model in red. This map shows fairly high overall agreement between the two models despite the large additional array of features associated with upstream tiles that the enhancer model could identify as important. Agreement between each model’s set of top-30 most important features is close to 90%, and the vast majority of these features lie within [TSS - 1 kb, TSS + 500 nt] for both models. The few features in the enhancer model that are located further than 1kb upstream of the TSS are not among top10 most heavily weighted features (see **Supplementary Table 1**).

#### Supplementary Figure 15: TEP-ROE rank correlation plot

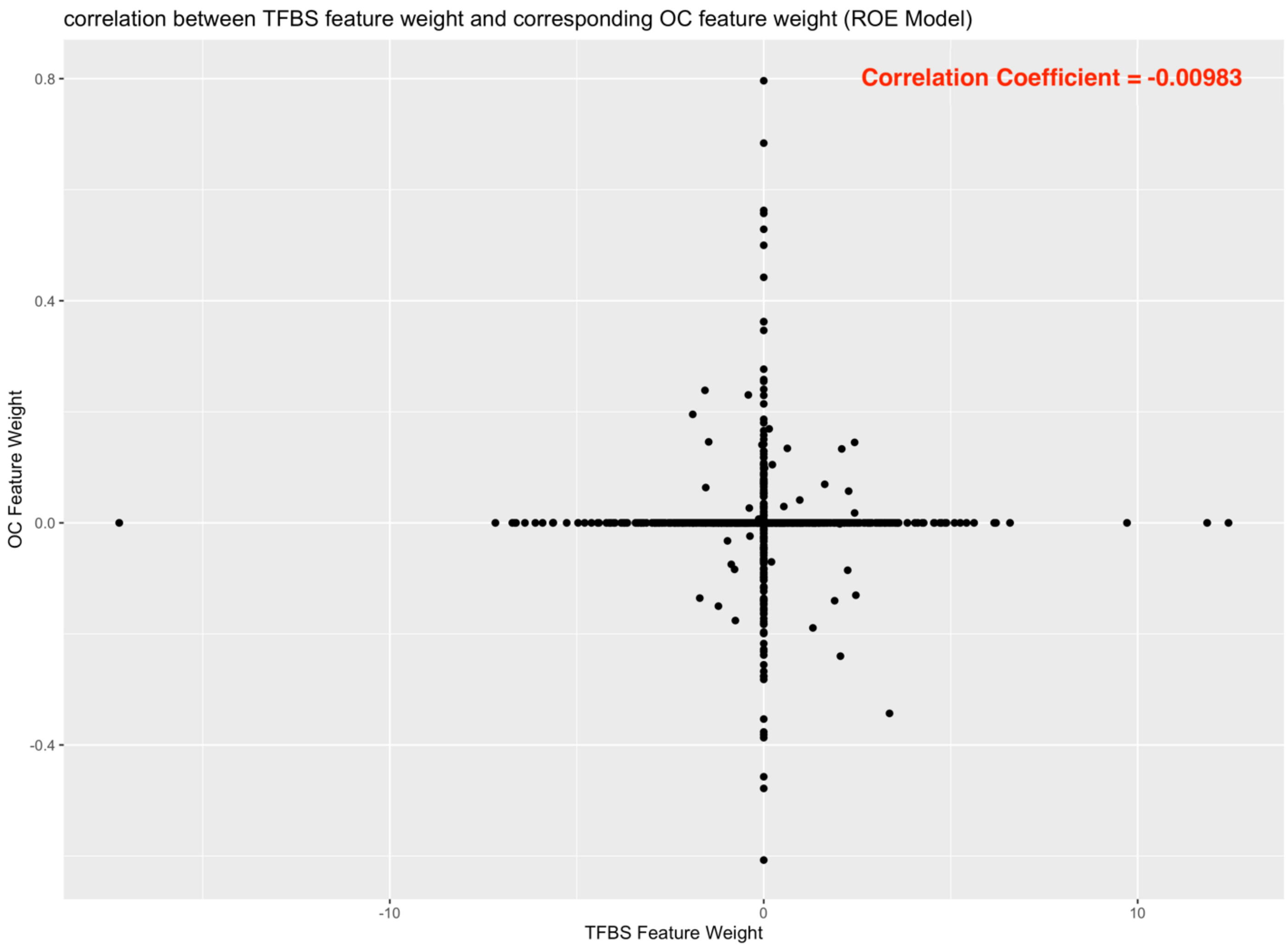

**Supplementary Figure 15**: Plot showing the correlation between TFBS and OC feature weights in the TEP-ROE model. This figure shows that TFBS features have no correlation with OC features in terms of model importance. The apparent visual positive correlation along the x and y axis derives from features with near-0 coefficients (these features were deemed 'unimportant' and essentially set to 0 by the regularization process in model training).

**Supplementary Figure 16: TEP-Tiled rank correlation plot**

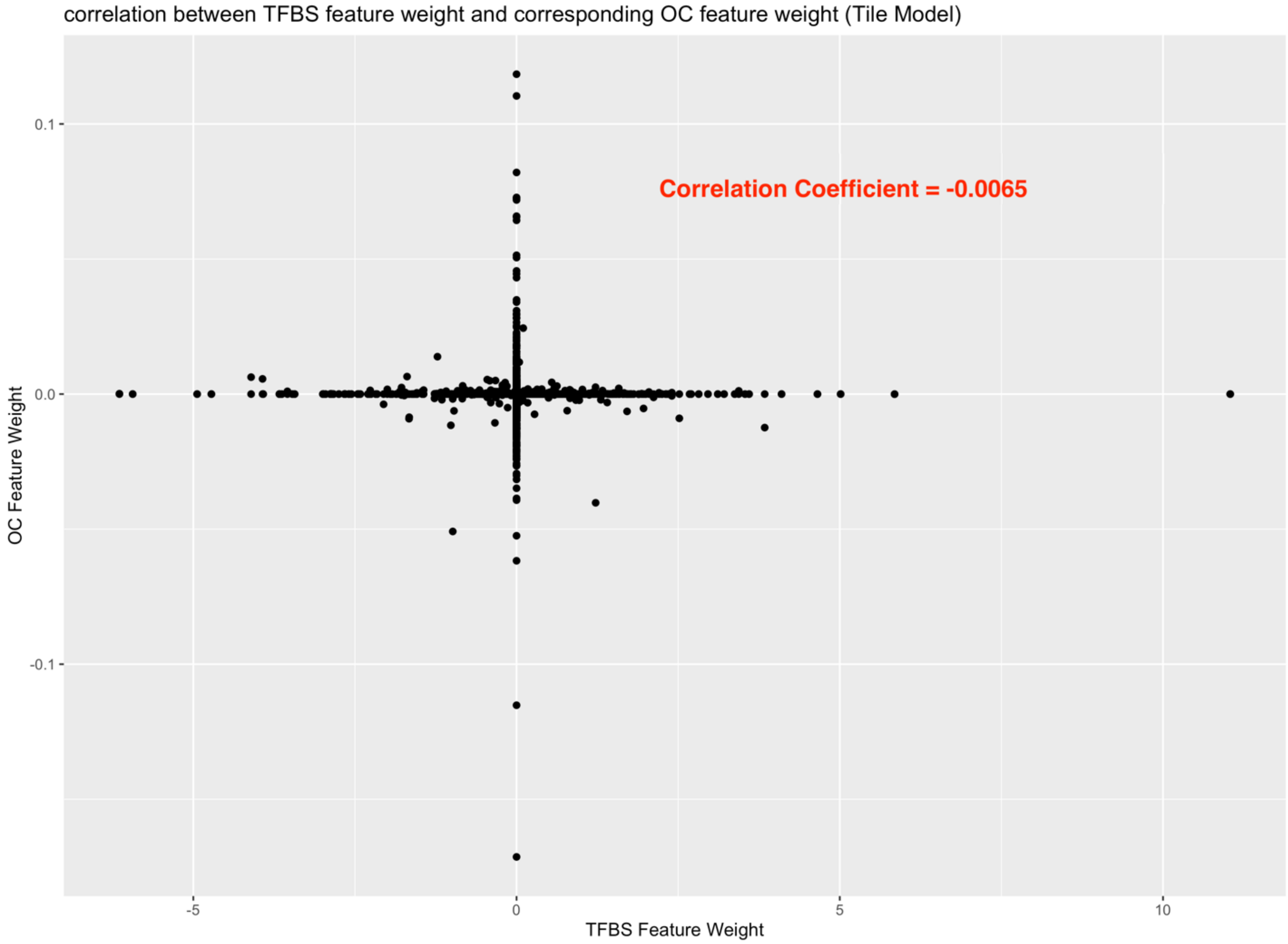

**Supplementary Figure 16**: Plot showing the correlation between TFBS and OC feature weights in the TEP-Tiled model. This figure shows that TFBS features have no correlation with OC features in terms of model importance. The apparent visual positive correlation along the x and y axis derives from features with near-0 coefficients (these features were deemed 'unimportant' and essentially set to 0 by the regularization process in model training).

**Supplementary Figure 17: ROE feature products vs openness**

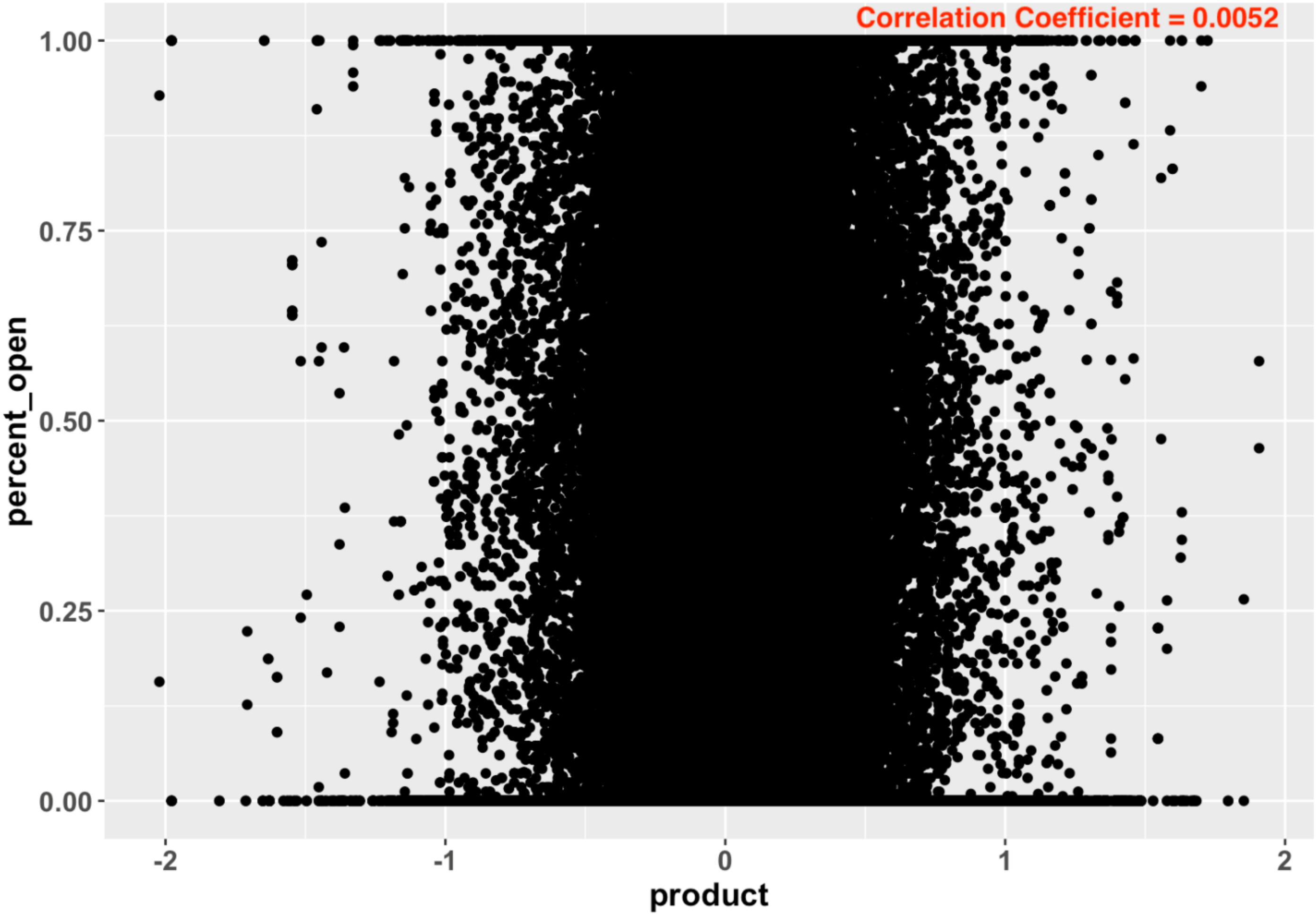

**Supplementary Figure 17:** Plot showing the correlation between products of TFBS feature value and TFBS model weight on the x-axis, and chromatin % openness on the y-axis, for the top 20 TFBS features in the TEP-ROE model. Each point represents an instance of the product vs. % openness in an individual promoter region. % openness was computed as the percentage of nucleotides within the TFBS feature region that are open (accessible). The correlation value is close to zero, indicating little/no relationship between TFBS feature importance and chromatin accessibility in the TFBS regions of individual promoters.

**Supplementary Figure 18: Tiled feature products vs openness**

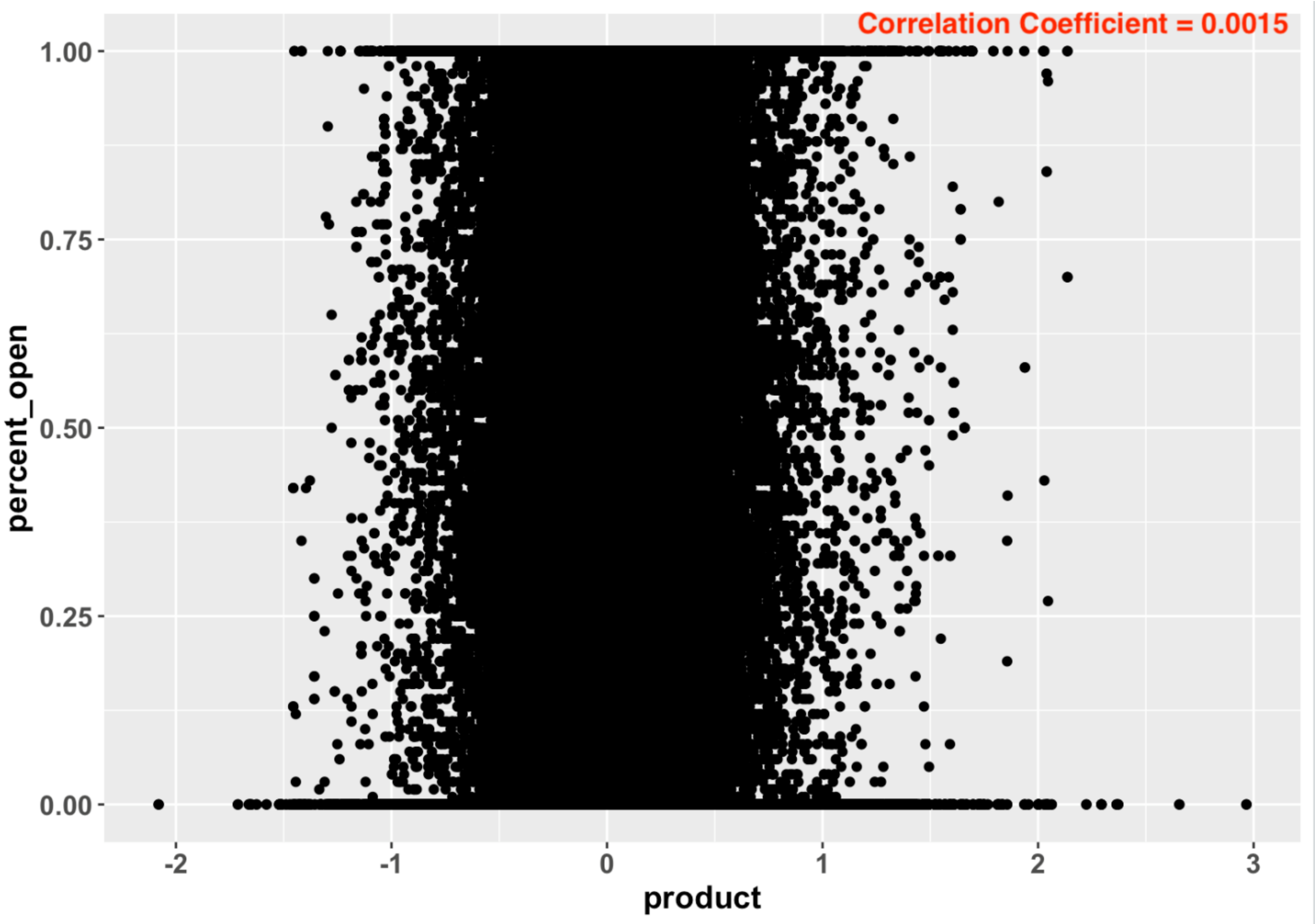

**Supplementary Figure 18:** Plot showing the correlation between products of TFBS feature value and TFBS model weight on the x-axis, and chromatin % openness on the y-axis, for the top 20 TFBS features in the TEP-Tiled model. Each point represents an instance of the product vs. % openness in an individual promoter region. % openness was computed as the percentage of nucleotides within the TFBS feature region that are open (accessible). The correlation value is close to zero, indicating little/no relationship between TFBS feature importance and chromatin accessibility in the TFBS regions of individual promoters.

#### Supplementary Figure 19: TEP-ROE model construction

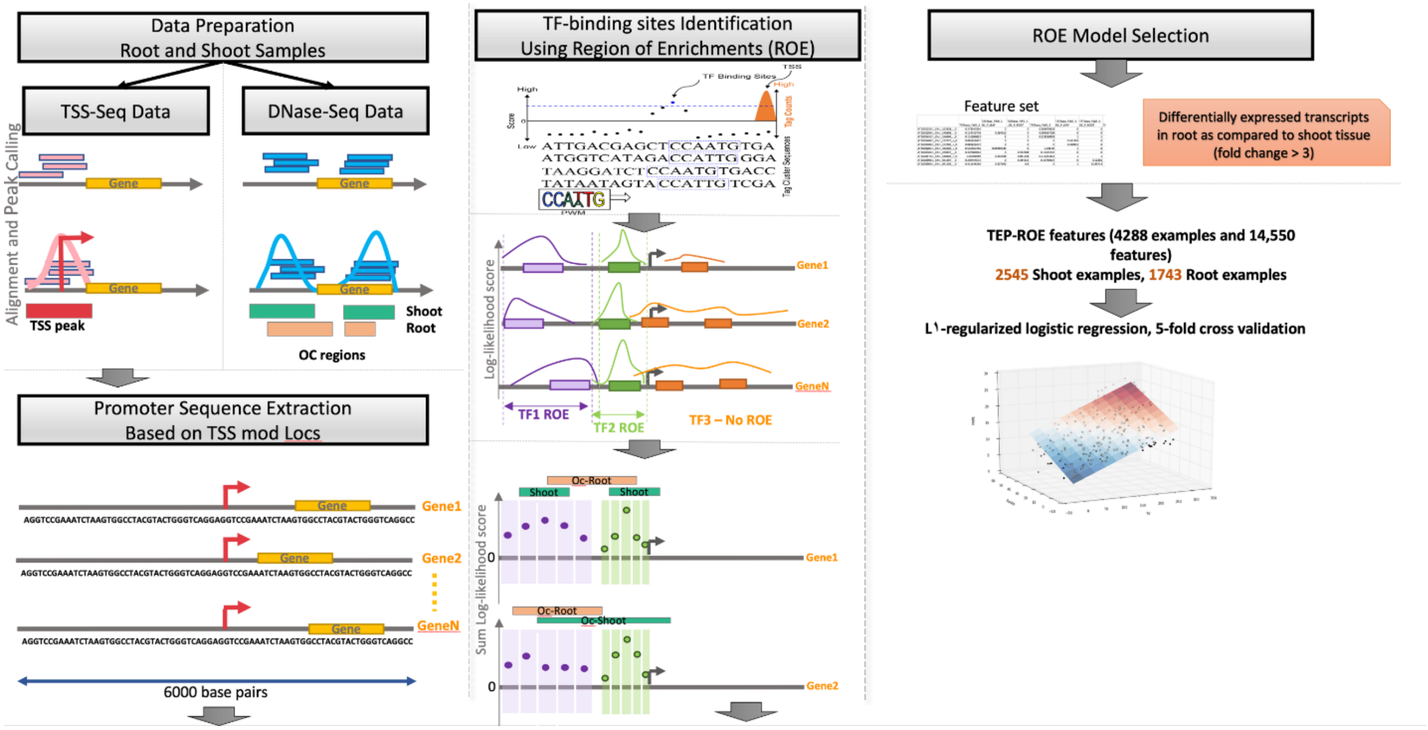

**Supplementary Figure 19:** The modeling process begins with raw dataset processing for TSS-seq, DNase-seq and RNA-seq datasets. This includes mapping reads to genome, calling peaks for both OC-openness and TSS-peak identification, and detecting differentially expressed transcripts to define class labels (root vs shoot). 6kb sequences were extracted (TSS - 3 kb, TSS + 3 kb, centered at each TSS mode), and TFBS log-likelihood summed scores (log-likelihood values above zero) were computed within this region for all PWMs (TF binding profiles). A Region of Enrichment (ROE) was identified for each TF (if present), and model features were generated within ROE regions. The TEP-ROE model training dataset contains 4288 samples, including 2545 shoot-expressed TSS promoters and 1743 root expressed TSS promoters. Finally, L1-regularized logistic regression was used to train and test the model. Using 5-fold cross-validation, the optimal regularization parameter was computed. Final auROC and auPRC metrics were reported on an independent held-out test set comprising 20% of the dataset.

#### Supplementary Figure 20: TEP-Tiled model construction

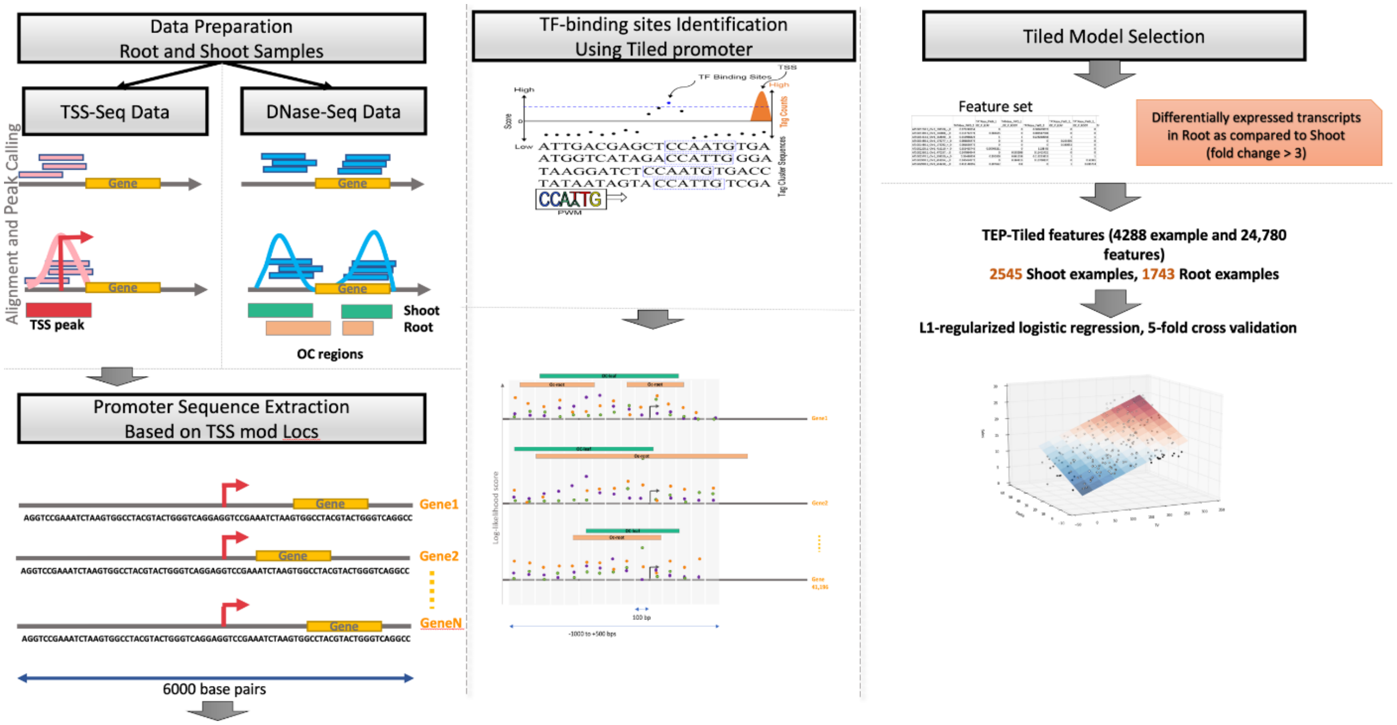

**Supplementary Figure 20:** The modeling process begins with raw dataset processing for TSS-seq, DNase-seq and RNA-seq datasets. This includes mapping reads to the genome, calling peaks for both OC-openness and TSS-peak identification, and detecting differentially expressed transcripts to define class labels (root vs shoot). 6kb sequences were extracted (TSS - 3 kb, TSS + 3 kb centered at each TSS mode), and TFBS log-likelihood summed scores (log-likelihood values above zero) were computed within this region for all PWMs (TF binding profiles). The region from 1000 nt upstream to 500 nt downstream of the TSS mode was divided into 100nt-wide tiles, and TFBS feature scores (log-likelihood sum scores) were computed within each tile. Model features were generated using TFBS scores and % OC openness in each tile (feature generation process within each Tile is identical to that performed within each ROE region in the ROE model). The training data in TEP-Tiled model contains 4288 samples, including 2545 shoot-expressed promoters and 1743 root expressed promoters. Finally, L1-regularized logistic regression was used to train and test the model. Using 5-fold cross-validation, the optimal regularization parameter was computed. Final auROC and auPRC metrics were reported on an independent held-out test set comprising 20% of the dataset.

**Supplementary Figure 21: Feature product sums for hard-codedness evaluation**

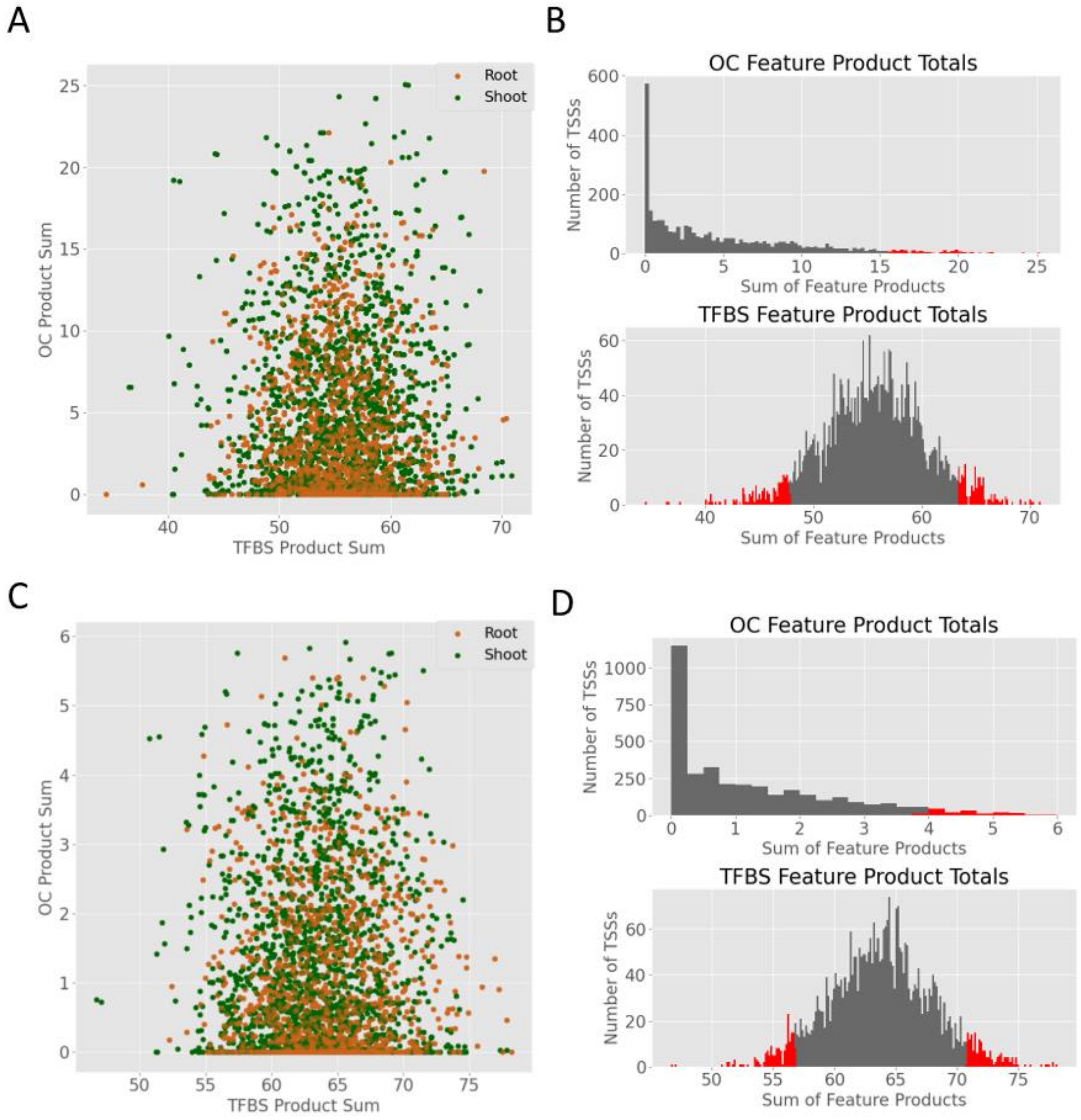

**Supplementary Figure 21:** A) Dot-plot displaying the sums of the TFBS feature products (feature value * feature weight assigned by model) and sums of the OC feature products (feature value * feature weight assigned by model) for each correctly classified TSS from the TEP-ROE model. Green dots indicate shoot; brown dots indicate root. B) Top histogram is a distribution of OC feature product sums from the TEP-ROE model. Bottom histogram is a distribution of TFBS feature product sums from the TEP-ROE model. The red bars represent TSSs that fell into the 5$^{th}$ and 95$^{th}$ percentiles. C) Dot-plot demonstrating the TFBS feature product sums and the OC feature product sums for each correctly classified TSS from the TEP-Tiled model. Green dots indicate shoot; brown dots indicate root. B) Top histogram is a distribution of OC feature product sums from the TEP-Tiled model. Bottom histogram is a distribution of TFBS feature product sums from the TEP-Tiled model. The red bars represent TSSs that fell into the 5$^{th}$ and 95$^{th}$ percentiles.

### Supplementary Tables

**Supplementary Table 1: Number of TSS peaks and their mapped locations**

| | No. TSS peaks | Mapped Location | | |
|---|---|---|---|---|
| | | <250 | 5’ utr | tss |
| Root | 26,040 | 2223 | 6047 | 17,770 |
| Shoot | 24,595 | 1636 | 5152 | 17,807 |

**Supplementary Table 1**: Table showing the number of total valid TSS peaks identified in each TSS-seq dataset, along with their mapped locations in the TAIR10 genome. The JAMM peak finder was used to identify TSS peaks from nanoCage-XL root and shoot samples. Only peaks having more than 50 reads and located in the region immediately upstream of gene body (within 500 nt) are considered “valid TSS peaks”.

**Supplementary Table 2: Number of TSS peaks and covered number of transcripts**

| Tissue | #TSS peaks mapped to individual Transcripts | | | | |
|---|---|---|---|---|---|
| | 1 Transcript | 2 Transcripts | 3 Transcripts | 4 Transcripts | 5 Transcripts |
| Root | 21,260 | 1950 | 236 | 35 | 4 |
| Shoot | 21,229 | 1416 | 128 | 30 | 6 |

**Supplementary Table 2:** Table showing the number of TSS peaks associated with one or more transcripts. Most TSS peaks are assigned to unique transcripts (first column left, “1 Transcript”). However, there are few TSS peaks in the vicinity of more than one transcripts in TAIR10 genome (within 500 nt).

**Supplementary Table 1: Basic statistics on TSS-seq, RNA-seq and OC data**

| | ROOT | SHOOT | TOTAL |
|---|---|---|---|
| No. Of TSS peaks (No. of Promoters) | 26,040 | 24,595 | 50,635 |
| No. of Transcripts with mapped TSS peak upstream | 23,487 (67% coverage) | 22,809 (65% coverage) | 25,751 (73%) |
| No. of Genes with mapped TSS peak upstream | 16,841(62%) | 16,086 (59%) | 18,580 (68%) |
| No. of closed Promoters | 656 (1.3%) | 639 (1.2%) | 123 closed in both (0.24%) |
| No. of DE Transcripts | 575 | 1037 | 1612 |
| No. of DE Genes | 562 | 963 | 1525 |
| No. of Tissue-specific Promoters | 1743 | 2545 | 4288 |

**Supplementary Table 3:** Total number of TAIR10 transcripts is 35,176 transcripts, which are associated with 27,206 protein coding genes. For simplicity in the row descriptions, a “Promoter” is equated to the [TSS - 3 kb, TSS + 3 kb] region surrounding a TSS. The first row shows the total number of TSS peaks used to extract [TSS - 3 kb, TSS + 3 kb] regions centered at TSS peak mode. The second row in the table shows the number of transcripts with TSS peaks upstream of the TAIR10 gene body (within 500 nt). The third row shows the number of TSS peaks covering TAIR10-annotated protein-coding genes. 68% of total TAIR annotated genes are assigned with at least one TSS peak in at least one of the tissues. The fourth row shows the number of TSS peaks with closed chromatin over the [TSS - 3 kb, TSS + 3 kb] region. Only 1% of the [TSS - 3 kb, TSS + 3 kb] regions, out of ~50,635 TSSs, are closed in one of the two tissues, and less than 0.3% are closed in both tissues. The last three rows show RNA-seq related information. A transcript is considered a tissue-specific or differentially expressed (DE) transcript if it expressed in both tissues (the expression value in one tissue is greater than 300 and the minimum expression value in the other tissue is greater than 30) and has log-fold-change above 3 (computed using RSEM). Read abundance was calculated on transcript level; we report differentially expressed genes as the number of genes that have at least one DE transcript. Finally, 4288 tissue specific TSSs and their [TSS - 3 kb, TSS + 3 kb] surrounding regions were used to train/test the ML model. Since the surrounding sequences are extracted from the set of TSS peaks, the TSSs assigned to DE transcripts are considered as tissue-specific (and their surrounding regions are considered Tissue-specific promoters).

**Supplementary Table 2: Basic RNA-seq expression statistics**

| | No. Transcripts (%Coverage) | No. Genes (%Coverage) |
|---|---|---|
| Low/No-expression in both tissues | 2,355 (7%) | 1,117 (4%) |
| Expressed in both tissues | 13,789 (40%) | 12,447 (46%) |
| Not-Differentially expressed | 12,177 (35%) | 10,922 (40%) |

**Supplementary Table 4:** Low or No-expression transcripts are those for which the mean normalized expression value is less than 30 in both tissue types. Genes that have low or no expression are those whose transcripts are all low/non-expressing. Transcripts expressed in both tissues are those for which the mean normalized expression values were greater than 300 in one tissue and greater than 30 in the other tissue. Percent coverage is out of 35,176 TAIR10 transcripts (for No. of Transcripts), or 27,206 protein coding genes (for No. of Genes).

**Supplementary Table 5: Top weighted features for 3PEAT root and shoot models**

| Shoot PWMs | Shoot coeffs | Root coeffs | Root PWMs |
|---|---|---|---|
| GAcontent | 0.902913 | 0.554167 | GAcontent |
| GCcontent | 0.506326 | 0.451861 | GCcontent |
| CAcontent | 0.359322 | 0.450633 | GA |
| M00502_TEIL_01 | 0.284998 | 0.375893 | M00702_SPF1_Q2 |
| M01126_BPC1_Q2 | 0.230441 | 0.257856 | TATAbox |
| RAV1-A_binding_site_motif | 0.221136 | 0.222318 | CAcontent |
| M00503_ATHB5_01 | 0.206872 | 0.207115 | Inr |
| M00355_PBF_01 | 0.201728 | 0.195353 | TEF-box_promoter_motif |
| Inr | 0.195206 | 0.189543 | RAV1-A_binding_site_motif |
| TATAbox | 0.190123 | 0.179117 | Bellringer_replumless_pennywise_BS1_IN_AG |
| M01136_DOF_Q2 | 0.18255 | 0.168393 | M00355_PBF_01 |
| Bellringer_replumless_pennywise_BS1_IN_AG | 0.177122 | 0.166626 | MYB4_binding_site_motif |
| M00439_C1_Q2 | 0.17623 | 0.15994 | M00314_GEN_INI3_B |
| Y_Patch | 0.171425 | 0.158837 | M01006_AGP1_01 |
| M00506_LIM1_01 | 0.165361 | 0.154919 | M01126_BPC1_Q2 |
| M01135_GAMYB_Q2 | 0.148444 | 0.152979 | M01050_ARR10_01 |
| GA | 0.146563 | 0.14662 | M00439_C1_Q2 |
| TEF-box_promoter_motif | 0.136267 | 0.137381 | M00653_OCSBF1_01 |
| MYB4_binding_site_motif | 0.131022 | 0.130946 | M01054_BHLH66_01 |
| SORLIP2 | 0.12592 | 0.128882 | Y_Patch |
| M00344_RAV1_02 | 0.122411 | 0.125033 | M00952_PCF5_01 |
| M00702_SPF1_Q2 | 0.119607 | 0.122956 | M00502_TEIL_01 |
| M01057_ERF2_01 | 0.113987 | 0.120953 | SORLIP2 |
| M01164_SQUA_01 | 0.113332 | 0.120847 | BoxII_promoter_motif |
| M00653_OCSBF1_01 | 0.109638 | 0.119398 | AtMYC2_BS_in_RD22 |
| M00315_GEN_INI_B | 0.100816 | 0.107276 | GCbox |
| M01050_ARR10_01 | 0.099942 | 0.102441 | M00506_LIM1_01 |
| Hexamer_promoter_motif | 0.098984 | 0.098304 | M01057_ERF2_01 |
| M01006_AGP1_01 | 0.093032 | 0.097558 | M00344_RAV1_02 |
| ATHB2_binding_site_motif | 0.088824 | 0.09406 | MYB3_binding_site_motif |
| M00440_CG1_Q6 | 0.088758 | 0.092642 | ATHB2_binding_site_motif |
| SORLIP5 | 0.087368 | 0.091277 | AG_BS_in_SPL_NOZ |
| T-box_promoter_motif | 0.085916 | 0.088917 | M01135_GAMYB_Q2 |
| BoxII_promoter_motif | 0.084432 | 0.084906 | M00635_GT1_Q6 |
| M01194_PDF2_01 | 0.083086 | 0.084657 | M01164_SQUA_01 |
| GCbox | 0.079116 | 0.081594 | M00503_ATHB5_01 |
| RAV1-B_binding_site_motif | 0.076355 | 0.081415 | M01194_PDF2_01 |
| DRE-like_promoter_motif | 0.075151 | 0.080094 | JASE2_motif_in_OPR1 |
| TELO-box_promoter_motif | 0.074697 | 0.079486 | DRE-like_promoter_motif |

| | | | |
|---|---|---|---|
| M00700_ROM_Q2 | 0.072984 | 0.079288 | M00353_DOF2_01 |
| M00438_ARF_Q2 | 0.07115 | 0.076477 | M01136_DOF_Q2 |
| M01188_CBNAC_01 | 0.070803 | 0.076344 | M00654_OSBZ8_Q6 |
| M00952_PCF5_01 | 0.069698 | 0.076107 | CCA1_binding_site_motif |
| M00370_CPRF3_Q2 | 0.067499 | 0.075843 | Hexamer_promoter_motif |
| E2F_DP_BS_in_AtCDC6 | 0.066471 | 0.074837 | M01133_AG_Q2 |
| M01130_PBF_Q2 | 0.066048 | 0.072843 | M00313_GEN_INI2_B |
| SORLREP3 | 0.062399 | 0.072721 | EveningElement_promoter_motif |
| AG_BS_in_SPL_NOZ | 0.062081 | 0.071576 | T-box_promoter_motif |
| M00376_TGA1A_Q2 | 0.060794 | 0.067971 | M00404_MADSB_Q2 |
| M01021_ID1_01 | 0.059488 | 0.066099 | AtMYB2_BS_in_RD22 |

**Supplementary Table 5.** Top 50 most heavily weighted features for the 3PEAT nanoCAGE-XL root-trained model and shoot-trained model, with their respective model coefficient weights.

**Supplementary Table 6: Cross-tissue model performance**

| Tissue of Training Dataset | Tissue of Test Dataset | |
|---|---|---|
| | Shoot | Root |
| Root | auROC: 0.98<br>auPRC:0.79 | auROC:0.98<br>auPRC:0.80 |
| Shoot | auROC: 0.98<br>auPRC: 0.80 | auROC:0.98<br>auPRC: 0.78 |

**Supplementary Table 6.** 3PEAT model performance outcomes for TSS location prediction in a given tissue sample. The training dataset is the collection of genomic sites (locations that are highly expressed TSSs in a tissue, or are not TSSs in a tissue) on which a classifier model was trained; the test dataset is the collection of genomic sites on which a model was tested to yield a performance measure.

**Supplementary Table 7: Cross-tissue 3PEAT-style GO-analysis for misclassified TSSs**

| GO-term | Description | p-value |
|---|---|---|
| GO: 0010449 | root meristem growth | 0.024373259 |
| GO: 0010102 | lateral root morphogenesis | 0.045790547 |
| GO: 0010101 | post-embryonic root morphogenesis | 0.045790547 |
| GO: 0010449 | root meristem growth | 0.03363946 |
| GO: 2000280 | regulation of root development | 0.029422773 |
| GO: 0010101 | post-embryonic root morphogenesis | 0.043180084 |
| GO: 0010102 | lateral root morphogenesis | 0.043180084 |
| GO: 0010449 | root meristem growth | 0.022479093 |
| GO: 0090057 | root radial pattern formation | 0.047136064 |
| GO: 0090057 | root radial pattern formation | 0.044521176 |
| GO: 0010101 | post-embryonic root morphogenesis | 0.040432343 |
| GO: 0010102 | lateral root morphogenesis | 0.040432343 |
| GO: 0010449 | root meristem growth | 0.021048648 |

**Supplementary Table 7:** GO-term enrichment analysis of the TSSs misclassified by the 3PEAT-style single tissue models. All of the significant terms for the misclassified TSSs are related to root development. Only terms with a p-value < 0.05 are reported here.

**Supplementary Table 8: TEP-Tiled vs TEP-ROE models top feature comparison**

| No. Top Features | No. of PWMs (TEP-ROE) | No. of PWMs (TEP-Tiled) | No. of Common PWMs | No. of Distinct PWMs | % common |
|---|---|---|---|---|---|
| 2 | 1 | 1 | 0 | 2 | 0.00000 |
| 5 | 3 | 4 | 1 | 6 | 14.28571 |
| 7 | 5 | 6 | 2 | 8 | 20.00000 |
| 10 | 7 | 9 | 4 | 9 | 30.76923 |
| 12 | 9 | 11 | 4 | 13 | 23.52941 |
| 15 | 12 | 13 | 5 | 16 | 23.80952 |
| 20 | 17 | 16 | 6 | 21 | 22.22222 |
| 50 | 36 | 40 | 15 | 50 | 23.07692 |
| 100 | 75 | 72 | 28 | 92 | 23.33333 |
| 150 | 99 | 92 | 43 | 112 | 27.74194 |
| 200 | 122 | 111 | 57 | 127 | 30.97826 |
| 300 | 151 | 144 | 82 | 147 | 35.80786 |
| 400 | 171 | 161 | 112 | 155 | 41.94757 |
| 500 | 182 | 173 | 135 | 144 | 48.38710 |
| 1000 | 189 | 189 | 157 | 101 | 60.85271 |

**Supplementary Table 8**: Differences among top weighted features between TEP-ROE and TEP-Tiled models. For a given number of top features from both the TEP-ROE and TEP-Tiled models, the number of PWMs representing only the features that agree between the two models was calculated; this number can be found in the “%common” column. As the number of features examined increases, top-feature agreement between the two models also increases.

#### Supplementary Table 9: Top 30 features of enhancer-covering Tiled model

| feature | pwm | strand | win | coef | type | left | right |
|---|---|---|---|---|---|---|---|
| OC_P_OVERALL_LEAF | OC_P_OVERALL_LEAF | FWD | 20 | 7.216611 | enhancer | -100 | 0 |
| M1663_1.02_FWD_19 | M1663_1.02 | FWD | 19 | 5.885361 | enhancer | -200 | -100 |
| M1691_1.02_FWD_19 | M1691_1.02 | FWD | 19 | -5.864451 | enhancer | -200 | -100 |
| M1309_1.02_REV_12 | M1309_1.02 | REV | 12 | 5.223288 | enhancer | -900 | -800 |
| M1686_1.02_REV_20 | M1686_1.02 | REV | 20 | -5.126922 | enhancer | -100 | 0 |
| M1689_1.02_REV_20 | M1689_1.02 | REV | 20 | -4.728434 | enhancer | -100 | 0 |
| M1691_1.02_FWD_21 | M1691_1.02 | FWD | 21 | -4.722934 | enhancer | 0 | 100 |
| M1323_1.02_FWD_19 | M1323_1.02 | FWD | 19 | -4.649934 | enhancer | -200 | -100 |
| M01194_PDF2_01_FWD_11 | M01194_PDF2_01 | FWD | 11 | 4.556978 | enhancer | -1000 | -900 |
| M1688_1.02_FWD_20 | M1688_1.02 | FWD | 20 | -4.449704 | enhancer | -100 | 0 |
| OC_P_OVERALL_ROOT | OC_P_OVERALL_ROOT | FWD | 20 | -4.440923 | enhancer | -100 | 0 |
| M0843_1.02_FWD_16 | M0843_1.02 | FWD | 16 | -4.135187 | enhancer | -500 | -400 |
| M1410_1.02_FWD_8 | M1410_1.02 | FWD | 8 | -4.005008 | enhancer | -1300 | -1200 |
| M1620_1.02_REV_4 | M1620_1.02 | REV | 4 | 3.864812 | enhancer | -1700 | -1600 |
| M0267_1.02_FWD_17 | M0267_1.02 | FWD | 17 | -3.863250 | enhancer | -400 | -300 |
| M1686_1.02_FWD_19 | M1686_1.02 | FWD | 19 | -3.746127 | enhancer | -200 | -100 |
| M0854_1.02_FWD_19 | M0854_1.02 | FWD | 19 | -3.693703 | enhancer | -200 | -100 |
| M1335_1.02_REV_20 | M1335_1.02 | REV | 20 | -3.687715 | enhancer | -100 | 0 |
| M1309_1.02_REV_20 | M1309_1.02 | REV | 20 | 3.676828 | enhancer | -100 | 0 |
| M0119_1.02_REV_19 | M0119_1.02 | REV | 19 | -3.535757 | enhancer | -200 | -100 |
| M0163_1.02_REV_25 | M0163_1.02 | REV | 25 | -3.526353 | enhancer | 400 | 500 |
| M01194_PDF2_01_FWD_19 | M01194_PDF2_01 | FWD | 19 | 3.507581 | enhancer | -200 | -100 |
| M1195_1.02_FWD_13 | M1195_1.02 | FWD | 13 | -3.506908 | enhancer | -800 | -700 |
| M0680_1.02_REV_4 | M0680_1.02 | REV | 4 | -3.460753 | enhancer | -1700 | -1600 |
| M1702_1.02_FWD_22 | M1702_1.02 | FWD | 22 | -3.440453 | enhancer | 100 | 200 |
| M1403_1.02_FWD_11 | M1403_1.02 | FWD | 11 | -3.437669 | enhancer | -1000 | -900 |
| M01191_HDG7_01_FWD_20 | M01191_HDG7_01 | FWD | 20 | 3.436033 | enhancer | -100 | 0 |
| M0646_1.02_REV_25 | M0646_1.02 | REV | 25 | -3.334678 | enhancer | 400 | 500 |
| M0259_1.02_REV_2 | M0259_1.02 | REV | 2 | -3.279065 | enhancer | -1900 | -1800 |
| M1688_1.02_FWD_18 | M1688_1.02 | FWD | 18 | -3.198103 | enhancer | -300 | -200 |

**Supplementary Table 9**: Top 30 features from the Tiled enhancer model, which computes features over tiles in the region [TSS - 2 kb, TSS + 500 nt]. The first column contains the full feature name. The “coef” column contains the feature’s model weight. A positive weight indicates shoot class association, and a negative weight indicates root class association. The right-most two columns show the genome coordinates (in nt) of the feature’s tile relative to TSS mode.

**Supplementary Table 10: Putatively "hardcoded" promoter examples from TEP-ROE**

| TSS ID | Tissue | Top TFBS | Associated TFs |
|---|---|---|---|
| AT1G31050.1_Chr1_11079176_-_0 | Root | M1940_1.02_FWD_5 | IDD1, IDD4, IDD6, IDD12, MGP, JKD |
| AT1G54940.1_Chr1_20481654_+_0 | Root | M0263_1.02_FWD_4 | bZIP17, bZIP49 |
| AT1G78660.2_Chr1_29585876_+_0 | Root | M0844_1.02_FWD_4 | ATHB-22 |
| AT2G16980.1_Chr2_7376364_+_0 | Root | M0142_1.02_REV_7 | "AT-hook containing protein" |
| AT2G16980.1_Chr2_7376366_+_0 | Root | M0010_1.02_FWD_5 | ERF9, ERF10, ERF110, ATERF14, ATABI4, RAP2.6 |
| AT2G16980.2_Chr2_7376364_+_0 | Root | M0142_1.02_REV_7 | "AT-hook containing protein" |
| AT2G16980.2_Chr2_7376366_+_0 | Root | M0010_1.02_FWD_5 | ERF9, ERF10, ERF110, ATERF14, ATABI4, RAP2.6 |
| AT4G08555.1_Chr4_5448141_+_0 | Root | M0371_1.02_REV_6 | ZFP8 |
| AT4G35380.1_Chr4_16819824_+_0 | Root | M0582_1.02_REV_4 | CAMTA2 |
| AT5G36970.1_Chr5_14605243_-_0 | Root | M1576_1.02_REV_1 | YAB3, YAB5 |
| AT3G09162.1_Chr3_2808252_-_0 | Shoot | M1344_1.02_FWD_4 | KUA1 |
| AT3G14330.1_Chr3_4782437_-_0 | Shoot | M2343_1.02_REV_3 | BZR1 |
| AT3G14330.1_Chr3_4782444_-_0 | Shoot | M2343_1.02_REV_3 | BZR1 |
| AT4G26555.1_Chr4_13406181_-_0 | Shoot | M1309_1.02_REV_5 | MYBD, MYBH |
| AT4G26555.1_Chr4_13406188_-_0 | Shoot | M1309_1.02_REV_5 | MYBD, MYBH |
| AT5G13730.1_Chr5_4430798_-_0 | Shoot | M0370_1.02_FWD_6 | ZAT9, STZ, AZF3 |
| AT5G60040.2_Chr5_24173327_+_0 | Shoot | M0119_1.02_REV_4 | AHL13, HAP3 |

**Supplementary Table 10:** Table containing TSS IDs that have putatively hard-coded promoters, computed from weights and feature values from the TEP-ROE model. Also listed is the top weighted TFBS for each TSS and the TFs that are associated with it.

| TSS ID | Tissue | Top TFBS | Associated TF |
|---|---|---|---|
| AT1G12160.1_Chr1_4126070_+_0 | Root | M0118_1.02_REV_9_tile100 | HMGA |
| AT1G33280.1_Chr1_12072606_+_0 | Root | M0142_1.02_FWD_1_tile100 | “AT-hook containing protein” |
| AT1G45015.1_Chr1_17021589_+_0 | Root | M0118_1.02_REV_9_tile100 | HMGA |
| AT1G45015.2_Chr1_17021589_+_0 | Root | M0118_1.02_REV_9_tile100 | HMGA |
| AT1G53680.1_Chr1_20038359_+_0 | Root | M0142_1.02_REV_14_tile100 | “AT-hook containing protein” |
| AT1G62990.1_Chr1_23337372_+_0 | Root | M0015_1.02_FWD_9_tile100 | DREB2, DREB2C, DREB2D |
| AT1G68150.1_Chr1_25543988_+_0 | Root | M0646_1.02_REV_15_tile100 | DOF1.5 |
| AT2G16970.1_Chr2_7369541_+_0 | Root | M0118_1.02_REV_9_tile100 | HMGA |
| AT2G16980.1_Chr2_7376364_+_0 | Root | M0118_1.02_REV_9_tile100 | HMGA |
| AT2G16980.1_Chr2_7376366_+_0 | Root | M0118_1.02_REV_9_tile100 | HMGA |
| AT2G16980.2_Chr2_7376364_+_0 | Root | M0118_1.02_REV_9_tile100 | HMGA |
| AT2G16980.2_Chr2_7376366_+_0 | Root | M0118_1.02_REV_9_tile100 | HMGA |
| AT3G05770.1_Chr3_1712207_-_0 | Root | M01126_BPC1_Q2_REV_12_tile100 | BPC1 |
| AT3G09270.1_Chr3_2849273_-_0 | Root | M0118_1.02_REV_9_tile100 | HMGA |
| AT4G18550.1_Chr4_10226979_-_0 | Root | M0118_1.02_REV_9_tile100 | HMGA |
| AT5G12420.1_Chr5_4026782_-_0 | Root | M0118_1.02_REV_9_tile100 | HMGA |
| AT5G40730.1_Chr5_16301102_+_0 | Root | M1663_1.02_FWD_9_tile100 | TCP1, TCP10, TCP13 |
| AT5G45920.1_Chr5_18622757_+_0 | Root | M0015_1.02_REV_15_tile100 | DREB2, DREB2C, DREB2D |
| AT5G45920.1_Chr5_18622778_+_0 | Root | M1274_1.02_REV_15_tile100 | ASL5, LBD3, LBD4 |
| AT5G65160.1_Chr5_26034000_-_0 | Root | M0080_1.02_REV_2_tile100 | EDF3 |
| AT1G51805.1_Chr1_19225648_-_0 | Shoot | M0118_1.02_REV_9_tile100 | HMGA |
| AT1G51805.2_Chr1_19225648_-_0 | Shoot | M0118_1.02_REV_9_tile100 | HMGA |
| AT1G52000.1_Chr1_19336089_-_0 | Shoot | M0372_1.02_REV_11_tile100 | ZAT1, ZAT4, ZAT9, AZF2 |
| AT1G64860.1_Chr1_24098017_+_0 | Shoot | M0118_1.02_REV_9_tile100 | HMGA |
| AT1G64860.2_Chr1_24098017_+_0 | Shoot | M0118_1.02_REV_9_tile100 | HMGA |
| AT1G76960.1_Chr1_28920925_-_0 | Shoot | M0118_1.02_REV_9_tile100 | HMGA |
| AT1G79040.1_Chr1_29736063_+_0 | Shoot | M01188_CBNAC_01_FWD_11_tile100 | CBNAC |
| AT3G01060.1_Chr3_18779_-_0 | Shoot | M0583_1.02_REV_10_tile100 | ATGRP2B, CSDP2 |
| AT3G01060.2_Chr3_18779_-_0 | Shoot | M0583_1.02_REV_10_tile100 | ATGRP2B, CSDP2 |
| AT3G03341.1_Chr3_790618_-_0 | Shoot | M0142_1.02_FWD_1_tile100 | “AT-hook containing protein” |
| AT3G51820.1_Chr3_19219005_-_0 | Shoot | M1274_1.02_REV_15_tile100 | ASL5, LBD3, LBD4 |
| AT3G51820.1_Chr3_19219011_-_0 | Shoot | M1274_1.02_REV_15_tile100 | ASL5, LBD3, LBD4 |
| AT3G52150.1_Chr3_19342053_+_0 | Shoot | M0372_1.02_REV_11_tile100 | ZAT1, ZAT4, ZAT9, AZF2 |
| AT3G52150.1_Chr3_19342057_+_0 | Shoot | M0118_1.02_REV_9_tile100 | HMGA |
| AT3G52150.2_Chr3_19342053_+_0 | Shoot | M0372_1.02_REV_11_tile100 | ZAT1, ZAT4, ZAT9, AZF2 |
| AT3G52150.2_Chr3_19342057_+_0 | Shoot | M0118_1.02_REV_9_tile100 | HMGA |
| AT4G17560.1_Chr4_9780335_+_0 | Shoot | M0118_1.02_REV_9_tile100 | HMGA |
| AT4G17560.1_Chr4_9780336_+_0 | Shoot | M0118_1.02_REV_9_tile100 | HMGA |
| AT4G26950.2_Chr4_13534252_-_0 | Shoot | M1274_1.02_REV_15_tile100 | ASL5, LBD3, LBD4 |
| AT5G13730.1_Chr5_4430798_-_0 | Shoot | M1309_1.02_REV_15_tile100 | MYBD, MYBH |
| AT5G24150.1_Chr5_8175404_-_0 | Shoot | M0118_1.02_REV_9_tile100 | HMGA |
| AT5G24150.2_Chr5_8175404_-_0 | Shoot | M0118_1.02_REV_9_tile100 | HMGA |

#### Supplementary Table 11: Putatively “hardcoded” promoter examples from TEP-Tiled

**Supplementary Table 11:** Table containing TSS IDs that have putatively hard-coded promoters, computed from weights and feature values from the TEP-Tiled model. Also listed is the top weighted TFBS for each TSS and the TFs that are associated with it.

**Supplementary Table 12: GO enrichment analyses for "hardcoded" promoter examples**

| GO-Term | Name | p-value | Model Type |
|---|---|---|---|
| GO:0015904 | tetracycline transmembrane transport | 0.000608 | TEP-ROE |
| GO:0046900 | tetrahydrofolylpolyglutamate metabolic process | 0.001823 | TEP-ROE |
| GO:0006855 | drug transmembrane transport | 0.001823 | TEP-ROE |
| GO:0015893 | drug transport | 0.002127 | TEP-ROE |
| GO:0032012 | regulation of ARF protein signal transduction | 0.00243 | TEP-ROE |
| GO:0032774 | RNA biosynthetic process | 0.002451 | TEP-ROE |
| GO:0046578 | regulation of Ras protein signal transduction | 0.002733 | TEP-ROE |
| GO:0051056 | regulation of small GTPase mediated signal transduction | 0.002733 | TEP-ROE |
| GO:0046483 | heterocycle metabolic process | 0.005007 | TEP-ROE |
| GO:1900865 | chloroplast RNA modification | 0.005763 | TEP-ROE |
| GO:0006725 | cellular aromatic compound metabolic process | 0.005993 | TEP-ROE |
| GO:0016554 | cytidine to uridine editing | 0.00667 | TEP-ROE |
| GO:1901360 | organic cyclic compound metabolic process | 0.006758 | TEP-ROE |
| GO:0016070 | RNA metabolic process | 0.006912 | TEP-ROE |
| GO:0015850 | organic hydroxy compound transport | 0.007576 | TEP-ROE |
| GO:0016553 | base conversion or substitution editing | 0.008482 | TEP-ROE |
| GO:0034654 | nucleobase-containing compound biosynthetic process | 0.010517 | TEP-ROE |
| GO:0045492 | xylan biosynthetic process | 0.011194 | TEP-ROE |
| GO:0006760 | folic acid-containing compound metabolic process | 0.011194 | TEP-ROE |
| GO:0042558 | pteridine-containing compound metabolic process | 0.011796 | TEP-ROE |
| GO:0009059 | macromolecule biosynthetic process | 0.012875 | TEP-ROE |
| GO:0009863 | salicylic acid mediated signaling pathway | 0.013599 | TEP-ROE |
| GO:0090304 | nucleic acid metabolic process | 0.014502 | TEP-ROE |
| GO:0006352 | DNA-templated transcription, initiation | 0.016598 | TEP-ROE |
| GO:0045491 | xylan metabolic process | 0.017496 | TEP-ROE |
| GO:1902531 | regulation of intracellular signal transduction | 0.017795 | TEP-ROE |
| GO:0034641 | cellular nitrogen compound metabolic process | 0.018046 | TEP-ROE |
| GO:0018130 | heterocycle biosynthetic process | 0.019841 | TEP-ROE |
| GO:0019438 | aromatic compound biosynthetic process | 0.024284 | TEP-ROE |
| GO:0006139 | nucleobase-containing compound metabolic process | 0.024824 | TEP-ROE |
| GO:0070592 | cell wall polysaccharide biosynthetic process | 0.02495 | TEP-ROE |
| GO:0070589 | cellular component macromolecule biosynthetic process | 0.026138 | TEP-ROE |
| GO:0044038 | cell wall macromolecule biosynthetic process | 0.026138 | TEP-ROE |
| GO:0071482 | cellular response to light stimulus | 0.028214 | TEP-ROE |
| GO:0006575 | cellular modified amino acid metabolic process | 0.028806 | TEP-ROE |
| GO:1901362 | organic cyclic compound biosynthetic process | 0.029586 | TEP-ROE |
| GO:0071478 | cellular response to radiation | 0.02999 | TEP-ROE |
| GO:0098656 | anion transmembrane transport | 0.032058 | TEP-ROE |
| GO:0010410 | hemicellulose metabolic process | 0.033533 | TEP-ROE |
| GO:0010383 | cell wall polysaccharide metabolic process | 0.043214 | TEP-ROE |
| GO:0050794 | regulation of cellular process | 0.047963 | TEP-ROE |
| GO:0015711 | organic anion transport | 0.048458 | TEP-ROE |
| GO:0015850 | organic hydroxy compound transport | 0.000144 | TEP-Tiled |
| GO:0006638 | neutral lipid metabolic process | 0.000285 | TEP-Tiled |

| | | | |
|---|---|---|---|
| GO:0006639 | acylglycerol metabolic process | 0.000285 | TEP-Tiled |
| GO:0006749 | glutathione metabolic process | 0.000451 | TEP-Tiled |
| GO:0009407 | toxin catabolic process | 0.000493 | TEP-Tiled |
| GO:0006352 | DNA-templated transcription, initiation | 0.000704 | TEP-Tiled |
| GO:0046462 | monoacylglycerol metabolic process | 0.000709 | TEP-Tiled |
| GO:0052651 | monoacylglycerol catabolic process | 0.000709 | TEP-Tiled |
| GO:0046340 | diacylglycerol catabolic process | 0.000709 | TEP-Tiled |
| GO:0009404 | toxin metabolic process | 0.00081 | TEP-Tiled |
| GO:0019748 | secondary metabolic process | 0.001285 | TEP-Tiled |
| GO:0071461 | cellular response to redox state | 0.001418 | TEP-Tiled |
| GO:0015904 | tetracycline transmembrane transport | 0.001418 | TEP-Tiled |
| GO:0009058 | biosynthetic process | 0.001494 | TEP-Tiled |
| GO:0010029 | regulation of seed germination | 0.00183 | TEP-Tiled |
| GO:1900140 | regulation of seedling development | 0.001953 | TEP-Tiled |
| GO:0006790 | sulfur compound metabolic process | 0.001969 | TEP-Tiled |
| GO:0071482 | cellular response to light stimulus | 0.002038 | TEP-Tiled |
| GO:0006575 | cellular modified amino acid metabolic process | 0.002124 | TEP-Tiled |
| GO:0010270 | photosystem II oxygen evolving complex assembly | 0.002127 | TEP-Tiled |
| GO:0080005 | photosystem stoichiometry adjustment | 0.002127 | TEP-Tiled |
| GO:0071478 | cellular response to radiation | 0.002302 | TEP-Tiled |
| GO:0098754 | detoxification | 0.002393 | TEP-Tiled |
| GO:0046461 | neutral lipid catabolic process | 0.002835 | TEP-Tiled |
| GO:0046464 | acylglycerol catabolic process | 0.002835 | TEP-Tiled |
| GO:0080148 | negative regulation of response to water deprivation | 0.003543 | TEP-Tiled |
| GO:1901362 | organic cyclic compound biosynthetic process | 0.003586 | TEP-Tiled |
| GO:0051775 | response to redox state | 0.00425 | TEP-Tiled |
| GO:0046339 | diacylglycerol metabolic process | 0.00425 | TEP-Tiled |
| GO:0006855 | drug transmembrane transport | 0.00425 | TEP-Tiled |
| GO:0044249 | cellular biosynthetic process | 0.004261 | TEP-Tiled |
| GO:0046503 | glycerolipid catabolic process | 0.004956 | TEP-Tiled |
| GO:0015893 | drug transport | 0.004956 | TEP-Tiled |
| GO:0008150 | biological_process | 0.005045 | TEP-Tiled |
| GO:1901576 | organic substance biosynthetic process | 0.005138 | TEP-Tiled |
| GO:0046486 | glycerolipid metabolic process | 0.006851 | TEP-Tiled |
| GO:0104004 | cellular response to environmental stimulus | 0.007546 | TEP-Tiled |
| GO:0071214 | cellular response to abiotic stimulus | 0.007546 | TEP-Tiled |
| GO:0015918 | sterol transport | 0.008482 | TEP-Tiled |
| GO:0009987 | cellular process | 0.008497 | TEP-Tiled |
| GO:0044237 | cellular metabolic process | 0.009157 | TEP-Tiled |
| GO:1901001 | negative regulation of response to salt stress | 0.009185 | TEP-Tiled |
| GO:0034641 | cellular nitrogen compound metabolic process | 0.010425 | TEP-Tiled |
| GO:0048829 | root cap development | 0.010591 | TEP-Tiled |
| GO:0071704 | organic substance metabolic process | 0.011325 | TEP-Tiled |
| GO:0008152 | metabolic process | 0.01244 | TEP-Tiled |
| GO:0032774 | RNA biosynthetic process | 0.013147 | TEP-Tiled |
| GO:0018130 | heterocycle biosynthetic process | 0.014533 | TEP-Tiled |
| GO:1901259 | chloroplast rRNA processing | 0.014797 | TEP-Tiled |
| GO:2001141 | regulation of RNA biosynthetic process | 0.015672 | TEP-Tiled |

| | | | |
|---|---|---|---|
| GO:0010187 | negative regulation of seed germination | 0.017592 | TEP-Tiled |
| GO:0019432 | triglyceride biosynthetic process | 0.017592 | TEP-Tiled |
| GO:0051252 | regulation of RNA metabolic process | 0.018883 | TEP-Tiled |
| GO:0010207 | photosystem II assembly | 0.018986 | TEP-Tiled |
| GO:0010192 | mucilage biosynthetic process | 0.018986 | TEP-Tiled |
| GO:0019438 | aromatic compound biosynthetic process | 0.019332 | TEP-Tiled |
| GO:0019915 | lipid storage | 0.019683 | TEP-Tiled |
| GO:0046460 | neutral lipid biosynthetic process | 0.019683 | TEP-Tiled |
| GO:0046463 | acylglycerol biosynthetic process | 0.019683 | TEP-Tiled |
| GO:0010025 | wax biosynthetic process | 0.020379 | TEP-Tiled |
| GO:0006629 | lipid metabolic process | 0.02054 | TEP-Tiled |
| GO:2000652 | regulation of secondary cell wall biogenesis | 0.021074 | TEP-Tiled |
| GO:0010166 | wax metabolic process | 0.021074 | TEP-Tiled |
| GO:0019219 | regulation of nucleobase-containing compound metabolic process | 0.021157 | TEP-Tiled |
| GO:0006641 | triglyceride metabolic process | 0.021769 | TEP-Tiled |
| GO:1901570 | fatty acid derivative biosynthetic process | 0.021769 | TEP-Tiled |
| GO:0010556 | regulation of macromolecule biosynthetic process | 0.022065 | TEP-Tiled |
| GO:0016126 | sterol biosynthetic process | 0.022464 | TEP-Tiled |
| GO:0010191 | mucilage metabolic process | 0.022464 | TEP-Tiled |
| GO:0048868 | pollen tube development | 0.023158 | TEP-Tiled |
| GO:1901000 | regulation of response to salt stress | 0.023158 | TEP-Tiled |
| GO:0010089 | xylem development | 0.024545 | TEP-Tiled |
| GO:2000070 | regulation of response to water deprivation | 0.02593 | TEP-Tiled |
| GO:0031326 | regulation of cellular biosynthetic process | 0.025991 | TEP-Tiled |
| GO:0009889 | regulation of biosynthetic process | 0.027707 | TEP-Tiled |
| GO:0019761 | glucosinolate biosynthetic process | 0.028003 | TEP-Tiled |
| GO:0016144 | S-glycoside biosynthetic process | 0.028003 | TEP-Tiled |
| GO:0019758 | glycosinolate biosynthetic process | 0.028003 | TEP-Tiled |
| GO:0047484 | regulation of response to osmotic stress | 0.029384 | TEP-Tiled |
| GO:0010109 | regulation of photosynthesis | 0.029384 | TEP-Tiled |
| GO:0015995 | chlorophyll biosynthetic process | 0.030073 | TEP-Tiled |
| GO:0048856 | anatomical structure development | 0.030401 | TEP-Tiled |
| GO:1901568 | fatty acid derivative metabolic process | 0.030762 | TEP-Tiled |
| GO:1903338 | regulation of cell wall organization or biogenesis | 0.032826 | TEP-Tiled |
| GO:0048580 | regulation of post-embryonic development | 0.033762 | TEP-Tiled |
| GO:0006779 | porphyrin-containing compound biosynthetic process | 0.034885 | TEP-Tiled |
| GO:0044271 | cellular nitrogen compound biosynthetic process | 0.036955 | TEP-Tiled |
| GO:0033014 | tetrapyrrole biosynthetic process | 0.037625 | TEP-Tiled |
| GO:1901360 | organic cyclic compound metabolic process | 0.038226 | TEP-Tiled |
| GO:0051171 | regulation of nitrogen compound metabolic process | 0.039126 | TEP-Tiled |
| GO:0006694 | steroid biosynthetic process | 0.041039 | TEP-Tiled |
| GO:0015994 | chlorophyll metabolic process | 0.042401 | TEP-Tiled |
| GO:0009834 | plant-type secondary cell wall biogenesis | 0.042401 | TEP-Tiled |
| GO:1901659 | glycosyl compound biosynthetic process | 0.043081 | TEP-Tiled |
| GO:0080090 | regulation of primary metabolic process | 0.044226 | TEP-Tiled |
| GO:0006518 | peptide metabolic process | 0.045872 | TEP-Tiled |
| GO:0044248 | cellular catabolic process | 0.045971 | TEP-Tiled |
| GO:0048519 | negative regulation of biological process | 0.046071 | TEP-Tiled |

| | | | |
|---|---|---|---|
| GO:0006807 | nitrogen compound metabolic process | 0.047825 | TEP-Tiled |
| GO:0016125 | sterol metabolic process | 0.04986 | TEP-Tiled |

**Supplementary Table 12:** GO-term enrichment analysis for the genes associated with the putatively "hard-coded" promoters. Included in this table are the GO-term, the description of the term, the p-value (all $p < 0.05$), and an indication of which model the set "hard-coded" promoters came from. Both GO-term enrichment experiments are contained in this single table.

**Supplementary Table 13: "In-silico knockout" results for TEP-ROE model**

| shift_amount | feature_id | Post-knockout prob1 | Pre-knockout prob1 | tss_name |
|---|---|---|---|---|
| -0.525761141 | M0119_1.02_REV_4 | 0.803841825 | 0.278080684 | AT5G52040.4_Chr5_21130346_+_0 |
| -0.525761141 | M0119_1.02_REV_4 | 0.803841825 | 0.278080684 | AT5G52040.3_Chr5_21130346_+_0 |
| -0.525761141 | M0119_1.02_REV_4 | 0.803841825 | 0.278080684 | AT5G52040.2_Chr5_21130346_+_0 |
| -0.525761141 | M0119_1.02_REV_4 | 0.803841825 | 0.278080684 | AT5G52040.1_Chr5_21130346_+_0 |
| -0.515625035 | M0119_1.02_REV_4 | 0.81283711 | 0.297212075 | AT5G03700.1_Chr5_967421_-_0 |
| -0.504363411 | M0119_1.02_REV_4 | 0.781646375 | 0.277282964 | AT5G59590.1_Chr5_24010616_-_0 |
| -0.501480944 | M0119_1.02_REV_4 | 0.758376904 | 0.256895961 | AT5G11810.1_Chr5_3808743_+_0 |
| -0.500727139 | M0119_1.02_REV_4 | 0.731933861 | 0.231206722 | AT5G11810.1_Chr5_3808742_+_0 |
| -0.475457796 | M0119_1.02_REV_4 | 0.727074621 | 0.251616825 | AT1G59640.2_Chr1_21911147_-_0 |
| -0.475457796 | M0119_1.02_REV_4 | 0.727074621 | 0.251616825 | AT1G59640.1_Chr1_21911147_-_0 |
| 0.460226045 | M0011_1.02_REV_7 | 0.231872302 | 0.692098347 | AT5G62720.2_Chr5_25191868_+_0 |
| 0.460226045 | M0011_1.02_REV_7 | 0.231872302 | 0.692098347 | AT5G62720.1_Chr5_25191868_+_0 |
| -0.4578229 | M1696_1.02_REV_5 | 0.728059863 | 0.270236963 | AT3G16565.2_Chr3_5642734_-_0 |
| -0.4578229 | M1696_1.02_REV_5 | 0.728059863 | 0.270236963 | AT3G16565.1_Chr3_5642734_-_0 |
| 0.449305754 | M0011_1.02_REV_7 | 0.259173323 | 0.708479077 | AT5G62720.2_Chr5_25191858_+_0 |
| 0.449305754 | M0011_1.02_REV_7 | 0.259173323 | 0.708479077 | AT5G62720.1_Chr5_25191858_+_0 |
| -0.444530928 | M0119_1.02_REV_4 | 0.893084762 | 0.448553834 | AT5G03700.1_Chr5_967404_-_0 |
| -0.442292868 | M0119_1.02_REV_4 | 0.716685211 | 0.274392343 | AT5G28640.1_Chr5_10649583_-_0 |
| -0.431771759 | M0119_1.02_REV_4 | 0.905423334 | 0.473651575 | AT5G52040.4_Chr5_21130352_+_0 |
| -0.431771759 | M0119_1.02_REV_4 | 0.905423334 | 0.473651575 | AT5G52040.3_Chr5_21130352_+_0 |
| -0.431771759 | M0119_1.02_REV_4 | 0.905423334 | 0.473651575 | AT5G52040.2_Chr5_21130352_+_0 |
| -0.431771759 | M0119_1.02_REV_4 | 0.905423334 | 0.473651575 | AT5G52040.1_Chr5_21130352_+_0 |
| -0.430064813 | M0119_1.02_REV_4 | 0.646933728 | 0.216868915 | AT2G23030.1_Chr2_9806673_-_0 |
| 0.428419104 | M1309_1.02_REV_5 | 0.245943985 | 0.674363089 | AT2G45170.2_Chr2_18624291_+_0 |
| 0.428419104 | M1309_1.02_REV_5 | 0.245943985 | 0.674363089 | AT2G45170.1_Chr2_18624291_+_0 |
| -0.426744547 | M0119_1.02_REV_4 | 0.570128701 | 0.143384154 | AT1G59640.2_Chr1_21911140_-_0 |
| -0.426744547 | M0119_1.02_REV_4 | 0.570128701 | 0.143384154 | AT1G59640.1_Chr1_21911140_-_0 |
| 0.426717636 | M1309_1.02_REV_5 | 0.335557429 | 0.762275065 | AT2G45170.2_Chr2_18624284_+_0 |
| 0.426717636 | M1309_1.02_REV_5 | 0.335557429 | 0.762275065 | AT2G45170.1_Chr2_18624284_+_0 |
| -0.426571534 | M0119_1.02_REV_4 | 0.4749933 | 0.048421766 | AT5G15190.2_Chr5_4933562_-_0 |
| -0.426571534 | M0119_1.02_REV_4 | 0.4749933 | 0.048421766 | AT5G15190.1_Chr5_4933562_-_0 |
| -0.414274796 | M2347_1.02_REV_2 | 0.888924088 | 0.474649292 | AT3G14100.1_Chr3_4672952_+_0 |
| -0.404004807 | M0119_1.02_REV_4 | 0.827577491 | 0.423572685 | AT3G56380.1_Chr3_20905295_+_0 |

**Supplementary Table 13:** The results from our in silico knockout experiments for the TEP-ROE model, sorted by the absolute value of the change in probability of expression in shoot (|shift_amount|). The "feature_id" column contains the name of the feature that was zeroed out for the experiment, while the "tss_name" column contains the name of the TSS that crossed the model's decision boundary. "Pre-knockout prob1" and "Post-knockout prob1" are the pre- and post- knockout probabilities of expression in shoot for the given TSS (1 – prob_value gives probability of expression in root for the given TSS).

**Supplementary Table 14: “In-silico knockout” results for TEP-Tiled model**

| shift_amount | feature_id | Post-knockout prob1 | Pre-knockout prob1 | tss_name |
|---|---|---|---|---|
| -0.525761141 | M0119_1.02_REV_4 | 0.80384183 | 0.27808068 | AT5G52040.4_Chr5_21130346_+_0 |
| -0.525761141 | M0119_1.02_REV_4 | 0.80384183 | 0.27808068 | AT5G52040.3_Chr5_21130346_+_0 |
| -0.525761141 | M0119_1.02_REV_4 | 0.80384183 | 0.27808068 | AT5G52040.2_Chr5_21130346_+_0 |
| -0.525761141 | M0119_1.02_REV_4 | 0.80384183 | 0.27808068 | AT5G52040.1_Chr5_21130346_+_0 |
| -0.515625035 | M0119_1.02_REV_4 | 0.81283711 | 0.29721208 | AT5G03700.1_Chr5_967421_-_0 |
| -0.504363411 | M0119_1.02_REV_4 | 0.78164638 | 0.27728296 | AT5G59590.1_Chr5_24010616_-_0 |
| -0.501480944 | M0119_1.02_REV_4 | 0.7583769 | 0.25689596 | AT5G11810.1_Chr5_3808743_+_0 |
| -0.500727139 | M0119_1.02_REV_4 | 0.73193386 | 0.23120672 | AT5G11810.1_Chr5_3808742_+_0 |
| -0.475457796 | M0119_1.02_REV_4 | 0.72707462 | 0.25161682 | AT1G59640.2_Chr1_21911147_-_0 |
| -0.475457796 | M0119_1.02_REV_4 | 0.72707462 | 0.25161682 | AT1G59640.1_Chr1_21911147_-_0 |
| 0.460226045 | M0011_1.02_REV_7 | 0.2318723 | 0.69209835 | AT5G62720.2_Chr5_25191868_+_0 |
| 0.460226045 | M0011_1.02_REV_7 | 0.2318723 | 0.69209835 | AT5G62720.1_Chr5_25191868_+_0 |
| -0.4578229 | M1696_1.02_REV_5 | 0.72805986 | 0.27023696 | AT3G16565.2_Chr3_5642734_-_0 |
| -0.4578229 | M1696_1.02_REV_5 | 0.72805986 | 0.27023696 | AT3G16565.1_Chr3_5642734_-_0 |
| 0.449305754 | M0011_1.02_REV_7 | 0.25917332 | 0.70847908 | AT5G62720.2_Chr5_25191858_+_0 |
| 0.449305754 | M0011_1.02_REV_7 | 0.25917332 | 0.70847908 | AT5G62720.1_Chr5_25191858_+_0 |
| -0.444530928 | M0119_1.02_REV_4 | 0.89308476 | 0.44855383 | AT5G03700.1_Chr5_967404_-_0 |
| -0.442292868 | M0119_1.02_REV_4 | 0.71668521 | 0.27439234 | AT5G28640.1_Chr5_10649583_-_0 |
| -0.431771759 | M0119_1.02_REV_4 | 0.90542333 | 0.47365158 | AT5G52040.4_Chr5_21130352_+_0 |
| -0.431771759 | M0119_1.02_REV_4 | 0.90542333 | 0.47365158 | AT5G52040.3_Chr5_21130352_+_0 |
| -0.431771759 | M0119_1.02_REV_4 | 0.90542333 | 0.47365158 | AT5G52040.2_Chr5_21130352_+_0 |
| -0.431771759 | M0119_1.02_REV_4 | 0.90542333 | 0.47365158 | AT5G52040.1_Chr5_21130352_+_0 |
| -0.430064813 | M0119_1.02_REV_4 | 0.64693373 | 0.21686892 | AT2G23030.1_Chr2_9806673_-_0 |
| 0.428419104 | M1309_1.02_REV_5 | 0.24594399 | 0.67436309 | AT2G45170.2_Chr2_18624291_+_0 |
| 0.428419104 | M1309_1.02_REV_5 | 0.24594399 | 0.67436309 | AT2G45170.1_Chr2_18624291_+_0 |
| -0.426744547 | M0119_1.02_REV_4 | 0.5701287 | 0.14338415 | AT1G59640.2_Chr1_21911140_-_0 |
| -0.426744547 | M0119_1.02_REV_4 | 0.5701287 | 0.14338415 | AT1G59640.1_Chr1_21911140_-_0 |
| 0.426717636 | M1309_1.02_REV_5 | 0.33555743 | 0.76227506 | AT2G45170.2_Chr2_18624284_+_0 |
| 0.426717636 | M1309_1.02_REV_5 | 0.33555743 | 0.76227506 | AT2G45170.1_Chr2_18624284_+_0 |
| -0.426571534 | M0119_1.02_REV_4 | 0.4749933 | 0.04842177 | AT5G15190.2_Chr5_4933562_-_0 |
| -0.426571534 | M0119_1.02_REV_4 | 0.4749933 | 0.04842177 | AT5G15190.1_Chr5_4933562_-_0 |
| -0.414274796 | M2347_1.02_REV_2 | 0.88892409 | 0.47464929 | AT3G14100.1_Chr3_4672952_+_0 |
| -0.404004807 | M0119_1.02_REV_4 | 0.82757749 | 0.42357268 | AT3G56380.1_Chr3_20905295_+_0 |

**Supplementary Table 14:** The results from our in silico knockout experiments for the TEP-Tiled model, sorted by the absolute value of the change in probability of expression in shoot (|shift_amount|). The “feature_id” column contains the name of the feature that was zeroed out for the experiment, while the “tss_name” column contains the name of the TSS that crossed the model’s decision boundary. “Pre-knockout prob1” and “Post-knockout prob1” are the pre- and post- knockout probabilities of expression in shoot for the given TSS (1 – prob_value gives probability of expression in root for the given TSS).
